# Behavioural choice emerges from nonlinear all-to-all interactions between drives

**DOI:** 10.1101/2020.03.12.989574

**Authors:** Stephen C. Thornquist, Michael A. Crickmore

## Abstract

Under the right conditions any drive can overcome nearly any other, yet studies of behavioural selection predominantly focus on only one, or occasionally two behaviours. We present an experimental and computational framework that captures and explains the resolution of conflicts between several competing motivations. We characterize neurons that integrate information from all rival drives to generate an aggregate signal that urges male Drosophila to transition out of mating. Experimental investigation of these Drive Integrating Neurons (DINs) revealed time-varying, supralinear interactions among competing drives that stimulate the DINs and induce a change in behaviour. Extending these findings to model the interactions between all of an animal’s motivations led to the surprising prediction that, under many conditions, all-to-all interactions actually buffer the dominant drive against challengers. We experimentally validated this prediction, showing that weak drives for a variety of tertiary goals can have a profound stabilizing effect on the ongoing behaviour. These results emerge only if non-linear integration of other motivations occurs for each of an animal’s drives, suggesting the potential universality of this mechanism. Our findings emphasize the interconnectedness of motivational systems and the consequent importance of considering the full motivational state of an animal to understand its behaviour.

## INTRODUCTION

Animals often have multiple unmet needs, and attempting to satisfy one generally precludes pursuing the others^1^. No one drive is strictly dominant; under the right conditions the pursuit of nearly any goal may be suppressed by another^2^. At some level behaviour-specific drive states must therefore affect the circuitry underlying many other behaviours^3^, and this information must be integrated to arrive at a consensus. The ethologist Konrad Lorenz used the metaphor of a “great parliament of instincts” to describe the behaviour of animals^2^, and the philosopher and mathematician Bertrand Russell noted in his Nobel Prize acceptance speech that “If you wish to know what men [sic] will do, you must know…the whole system of their desires with their relative strengths”^4^. Nearly all studies on the interactions between competing motivations, in contrast, focus on the resolution between just two drives in conflict. Here we establish an experimental and computational framework for examining the many interactions between simultaneous drive states that must be considered to understand naturalistic decision-making.

The mating duration of *Drosophila melanogaster* provides a clear and quantitative readout of the interplay between competing drives: to switch behaviours the male must first terminate the mating. If undisturbed, copulation will last ∼23 minutes; if a dangerous situation arises, the male may truncate the mating to flee, depending on both the severity of the threat and how far the mating has progressed^5^. For the first several minutes after initiating a mating, he will sacrifice his (and his partner’s) life in the face of a potentially lethal threat to ensure successful fertilization^6^, but his persistence (or propensity to sustain the mating when challenged) decreases as time passes, reflecting the increasing likelihood that the goals of mating have been achieved. Here we show how the changing properties of eight male-specific neurons^5^ (hereafter referred to as Drive Integrating Neurons, or DINs, **Figure 1a**) steer the resolution of multiple conflicting drives during mating.

**Figure 1:**
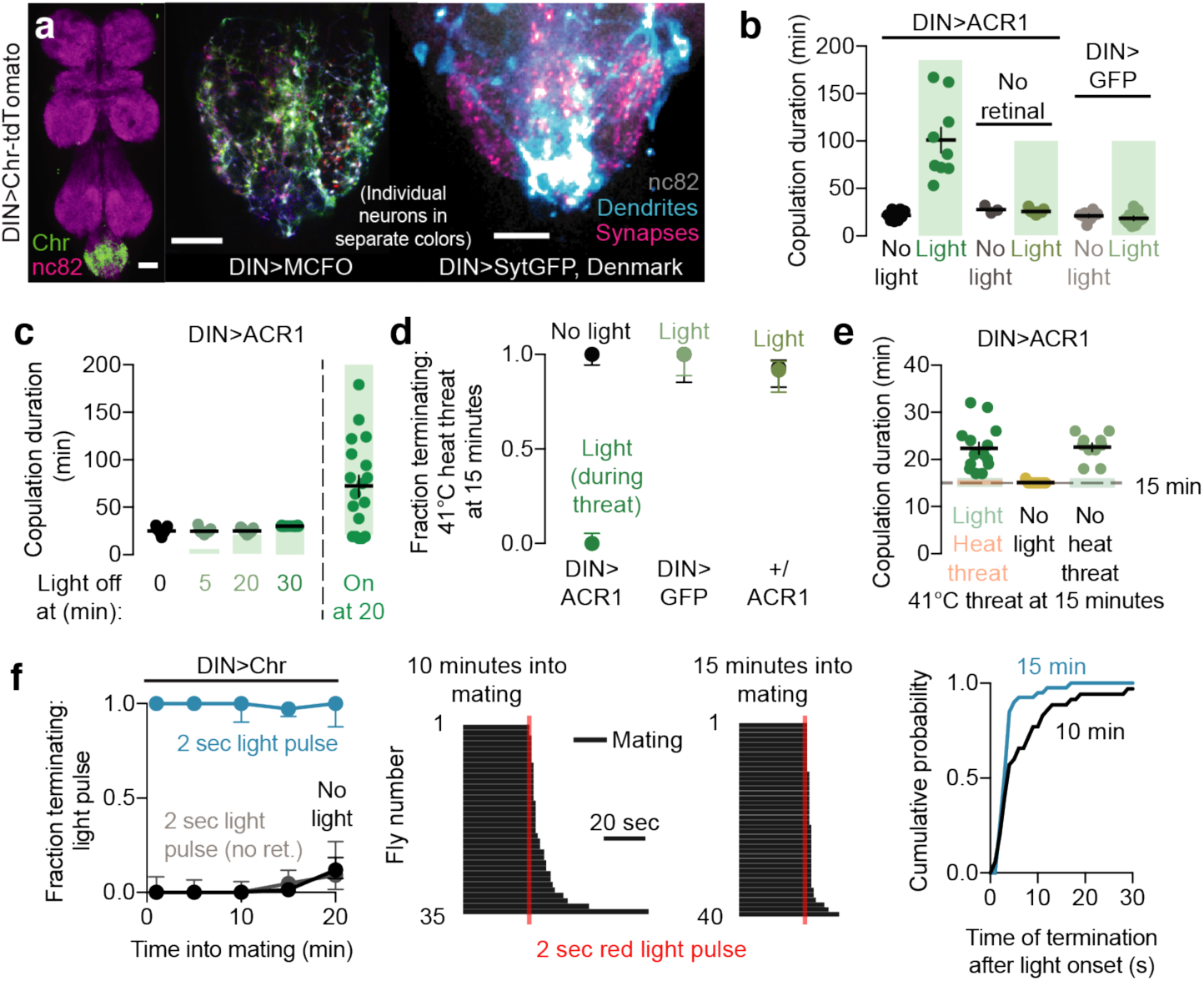
Drive Integrating Neurons (DINs) control the decision to terminate mating. **a)** Morphology of the DINs. Left: The DINs (labeled by NP2719-Gal4)^5^ are restricted to the abdominal ganglion of the ventral nervous system. Middle: Individual DINs together fill the lateral portions of the abdominal ganglion (image of a single optical section). Right: DIN dendrites selectively cover the midline tracts of the abdominal ganglion (blue) where most inputs from other parts of the nervous system converge^7^, and send projections to the local circuitry of the abdominal ganglion (magenta). Scale bars in all figures are 20 µm. **b)** Silencing the DINs using ACR1 and green light results in extremely long matings. Error bars here and in all other figures (unless otherwise noted) represent 67% credible intervals, chosen to resemble the standard error of the mean. For the number of samples in each experiment, see **Supplementary Table 2**. For statistical tests, see **Supplementary Table 3**. **c)** Electrical activity in the DINs is only necessary around the time of termination to end the mating: silencing from the beginning until near the natural end of mating does not affect copulation duration (third column), while silencing that begins just before the natural time of termination prolongs the mating by hours (last column). Matings in which the DINs are silenced through the normal ∼23 minute termination time are ended seconds after the light is turned off (fourth column). **d-e)** Acute silencing of the DINs prevents termination in response to heat threats (d). Transiently silencing the DINs during a heat threat prevents early termination but otherwise has no impact on the overall duration of mating (e). **f)** Acute optogenetic stimulation of the DINs causes termination within seconds. Left: Two seconds of stimulation is sufficient to terminate copulation regardless of how far the mating has progressed. “No ret.” refers to flies that were not fed retinal, the obligate chromophore for CsChrimson’s light sensitivity, showing that light *per se* does not cause termination of mating. Middle and Right: ethograms and cumulative distribution plot demonstrating the response to 2 seconds of optogenetic DIN stimulation delivered at 10 and 15 minutes into mating. Each black stripe in the ethograms represents a single mating.

## RESULTS

### Drive Integrating Neurons (DINs) control the decision to terminate mating

Constitutively silencing the DINs with tetanus toxin extends the average duration of mating from ∼23 minutes to ∼1.5 hours^5^. To examine their moment-to-moment function during mating and in response to threats, we conditionally silenced the DINs with GtACR1 (ACR1), a green-light gated chloride channel. While tonic optogenetic silencing extended copulation duration to a similar extent as tetanus toxin (**Figure 1b**), relaxing the inhibition at 30 minutes caused near-immediate termination of mating (**Figure 1c**). Inversely, turning on the light just before the normal time of termination (at 20 minutes) most often caused matings to last well over an hour (**Figure 1c**). These results show that electrical activity in the DINs is not required for tracking time during copulation but instead causes termination after the appropriate time has passed. Consistent with this interpretation, relaxing inhibition either shortly before the usual time of termination (20 minutes) or at 5 minutes into the mating allowed copulations to terminate at the appropriate time (**Figure 1c**). The temporal precision of these experiments shows that DIN activity is only required around the time of mating termination, overturning the idea derived from constitutive silencing experiments that these neurons promote a gradual decline in motivation to sustain the mating^5^.

In line with their requirement only around the moment of natural termination, transient DIN inhibition during the presentation of a threat caused the male to persist through severe challenges that would otherwise truncate nearly all late-stage matings (**Figure 1d** and **Video 1**), but did not extend copulation beyond its natural termination time (**Figure 1e**). The DINs are therefore specifically required to make the decision to terminate mating, as we confirmed using the heat-sensitive synaptic silencing tool UAS-Shibire^ts 8^ (**Extended Data Figure 1**), ruling out tool-specific artifacts (e.g. due to changes in the chloride equilibrium potential^9^). We conclude that DIN activity ends copulation in response to two types of triggering stimuli: (i) competing drives (e.g. survival in the case of heat threats); and (ii) the fulfillment of all mating goals at ∼23 minutes. At the level of DIN activity, there appears to be little, if any, difference between these two classes of demotivating conditions.

Using the red-light gated cation channel CsChrimson (Chr)^10^ we found that acute stimulation with a two-second pulse of red light induced termination of nearly 100% of matings (**Video 2** and **Figure 1f**) with a dismounting procedure resembling the response to threatening stimuli (**Video 3**). Termination induced by brief optogenetic stimulation occurred regardless of time into the mating (**Figure 1f**), and with a varying latency of up to 30 seconds after the stimulation pulse (**Figure 1f**), arguing against a startle or motor reflex. This latency was due to the sustained activity of the DINs, rather than a slow motor program: silencing the DINs immediately after optogenetic stimulation prevented termination (**Extended Data Figure 2a**). In our previous experiments, tonic thermogenetic activation of the DINs starting before copulation shortened matings, but did not cause immediate termination^5^. This was likely due to long timescale habituation, as we see a similar effect following mild optogenetic stimulation that commences before the initiation of copulation (**Extended Data Figures 2b**,**c**). The new, acute optogenetic activation and silencing experiments led us to name these neurons the Drive Integrating Neurons: they are the means through which competing drives (such as self-preservation) demotivate copulation in order to effect a change in behaviour.

### Demotivating stimuli integrate over an expanding time window as the mating progresses

Brief stimulation of the DINs (500 ms or 1 second) resulted in a probabilistic response to stimulation, like naturalistic demotivating conditions, but showed no reliable difference in termination probability if delivered at 10 or 15 minutes into mating (**Figure 2a**). Naturalistic demotivating conditions, in contrast, become more disruptive as the mating progresses^5^. This led us to the idea that only longer-lasting stimuli, such as sustained heat threats, generate responses that are enhanced as the mating progresses. Such a phenomenon could arise if the demotivation circuitry accumulates inputs over a relatively short time window early in mating, with an expanding window as the mating progresses. In models that integrate linearly over time (schematized in **Figure 2b**), a short integration window gives rise to similar peaks and only slightly more cumulative activation for longer stimuli of fixed intensity (**Extended Data Figure 4a**). If the time constant of integration (τ) is increased, sustained input can integrate over a longer time, increasing activation levels for a stimulus of the same intensity. Increasing the time constant has a strong effect on cumulative and peak output only when it was previously shorter than the duration of the stimulus (**Figure 2b**). *Vice versa*, increasing the duration of a demotivating stimulus only substantially enhances the output when the time constant is comparatively long (**Figure 2b**).

**Figure 2:**
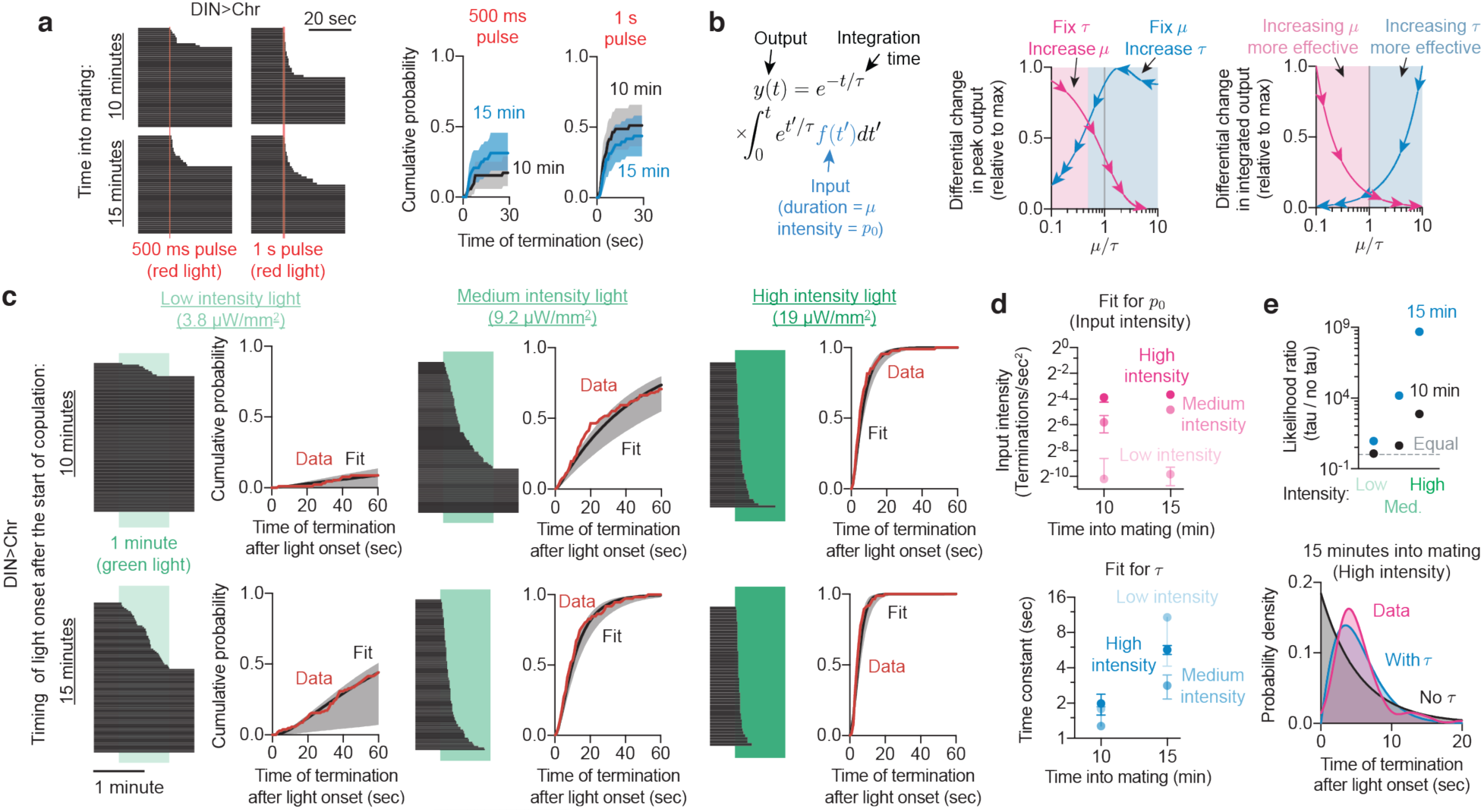
The DINs integrate inputs over time to oppose copulation. **a)** The response to brief pulses (500 ms and 1 sec) of DIN stimulation also shows little, if any, potentiation at 10 vs. 15 minutes into mating. **b)** A schematized system that performs temporal integration shows how increasing the time constant can potentiate the output of sustained inputs. Left: The output of this system is the summed response of every instant of input, where the effect of each moment of input decays with an exponential time constant of τ. Center and right: Both the peak (center) and integrated outputs (right) of the system are much more sensitive to changes in μ (the stimulus duration) when μ ≪ τ, whereas they are much more sensitive to changes in τ when μ ≫ τ. **c)** Termination times of DIN>Chr flies exposed to green light for sixty seconds (intensity indicated above graphs). Fitting the cumulative distribution to the model in **Extended Data Figure 3a** reveals a close fit. Solid black line: maximum likelihood fit. Error bars: pointwise 95% coverage intervals sampled according to estimated covariance of the parameters (**Extended Data Figure 3c**). **d)** Parameter estimates for τ (time constant) and *p*_0_ (intensity) across timepoints and conditions. *p*_0_ is sensitive to stimulation intensity but not time into mating, while τ scales with time into mating, but not stimulation intensity. Error bars show the square root of the estimated parameter variance using the Cramér-Rao bound. **e)** Temporal integration is necessary to explain the behaviour of flies during sustained optogenetic stimulation, as a model predicting no temporal integration (no τ) ascribes a much lower likelihood to the data sets observed (top plot). Temporal integration is also needed to explain the increasing probability of termination as the stimulus goes on (bottom plot, data fit with a kernel density estimate).

In the simplest such model, the instantaneous probability of responding to a stimulus behaves like a linear dynamical system with time constant τ. This enabled us to fit the cumulative distribution of actual times of termination to the equation

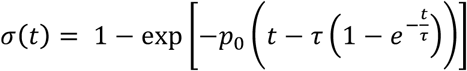

(where *p*_0_is the strength of the demotivating input) to estimate the parameters of the model (**Extended Data Figure 3a**, see Methods for more information). We reduced the strength of optogenetic activation by using green light, which penetrates tissue less effectively than red^11^, allowing us to stimulate the DINs for longer durations without immediately ending the mating. A shorter τ caps the peak termination probability for long threats of fixed intensity, whereas extending integration to longer timescales leads to an increased output and, therefore, increased termination probability.

To assess the overall fit of the data we plotted cumulative termination probabilities, which show that the data is well fit by a linear integration process in time, with τ ≈ 1-2 seconds at 10 minutes and ≈3-10 seconds at 15 minutes (**Figures 2c, d, and Extended Data Figures 3b,c**). Importantly, without either constraint being explicitly imposed, the model fit only predicted a change in the overall strength of input when the intensity of the light was changed (*p*_0_ ≈ 10^−3^ for low intensity light, 0.02 for medium intensity, and 0.1 for high intensity) and did not predict a change in time constant with different light intensities at the same point in mating (**Figure 2d**). The fit is orders of magnitude better when temporal integration is included than if the DINs are assumed not to integrate over time (**Figure 2e**). This analysis provides quantitative estimates for the expanding time constant as mating progresses, arguing that the response to competing drives is increased as the mating progresses by lengthening the integration time of the DINs. We emphasize that this model is descriptive: it does not claim that the parameters τ and *p*_0_ truly exist in some physical form. Instead, it provides a means to describe and analyze how the DINs integrate information over time and allows us to assess the demotivating impact of a memory-like component within this decision-making circuitry.

A temporal window of integration should potentiate responses not just to sustained challenges, but also to discrete inputs separated in time. We therefore stimulated the DINs with paired 500 ms excitatory pulses separated by 0-to-30 seconds at 10 or 15 minutes into mating. When the two DIN pulses were supplied in near-immediate succession, we found an augmentation of the second pulse at both 10 and 15 minutes (**Extended Data Figure 4b**). When the pulses were spaced out, augmentation was still evident with at longer inter-pulse intervals later in mating, closely matching the effects seen with sustained stimulation.

### Motivating inputs limit the ability of competing drives to activate the DINs

We used this quantitative analysis to ask whether inputs that motivate sustained copulation act by preventing stimulation of the DINs, either by decreasing the perceived intensity of the challenges or by shortening the time constant of integration. During the first five minutes of a mating, males will almost never terminate in response to even the most severe threats^5^. The duration of this period of insurmountably high motivation is determined by a molecular timer^6^ housed in four male-specific neurons that produce the neuropeptide Corazonin (Crz) (**Extended Data Figure 5a**)^12,13^. Silencing the Crz neurons dramatically extends the period of high motivation^6^, causing matings to last over an hour^13^. The DINs are functionally downstream of this switch in motivation, as two seconds of strong optogenetic DIN stimulation overrides Crz silencing and terminates matings whenever it is applied (**Extended Data Figure 5b**). Inversely, optogenetic activation of the Crz neurons very early in mating renders the male immediately responsive to threats, but this effect is prevented by silencing the DINs, and Crz stimulation cannot prevent the long mating duration caused by DIN silencing (**Figures 3a,b**). Sustained low-intensity DIN stimulation at 3 minutes into mating revealed an inferred time constant of ∼1.0 seconds (**Extended Data Figures 5c,d**), the smallest value resolvable by our approach (**Extended Data Figure 6, Supplementary Note 1**) and shorter than that seen at 10 minutes, indicating that the ability of the DINs to integrate inputs over time increases from the beginning to the end of the mating. The increasing integration time as the mating progresses does not require the Crz neurons, since silencing them had no effect on the time constant (**Figure 3c** and **Extended Data Figure 5c**). Instead, silencing the Crz neurons reduced the input intensity perceived by the DINs, *p*_0_, by nearly a factor of 10 (**Figure 3c** and **Extended Data Figure 5c**), rendering the DINs effectively inaccessible to demotivating inputs. These results reveal two adjustable properties of the DINs that determine the motivation to sustain mating at any moment: a time constant of integration that increases over the entire 23-minute mating, and a superimposed restriction on other drives’ ability to access the DINs for the first 6 minutes (**Extended Data Figure 5e**).

**Figure 3:**
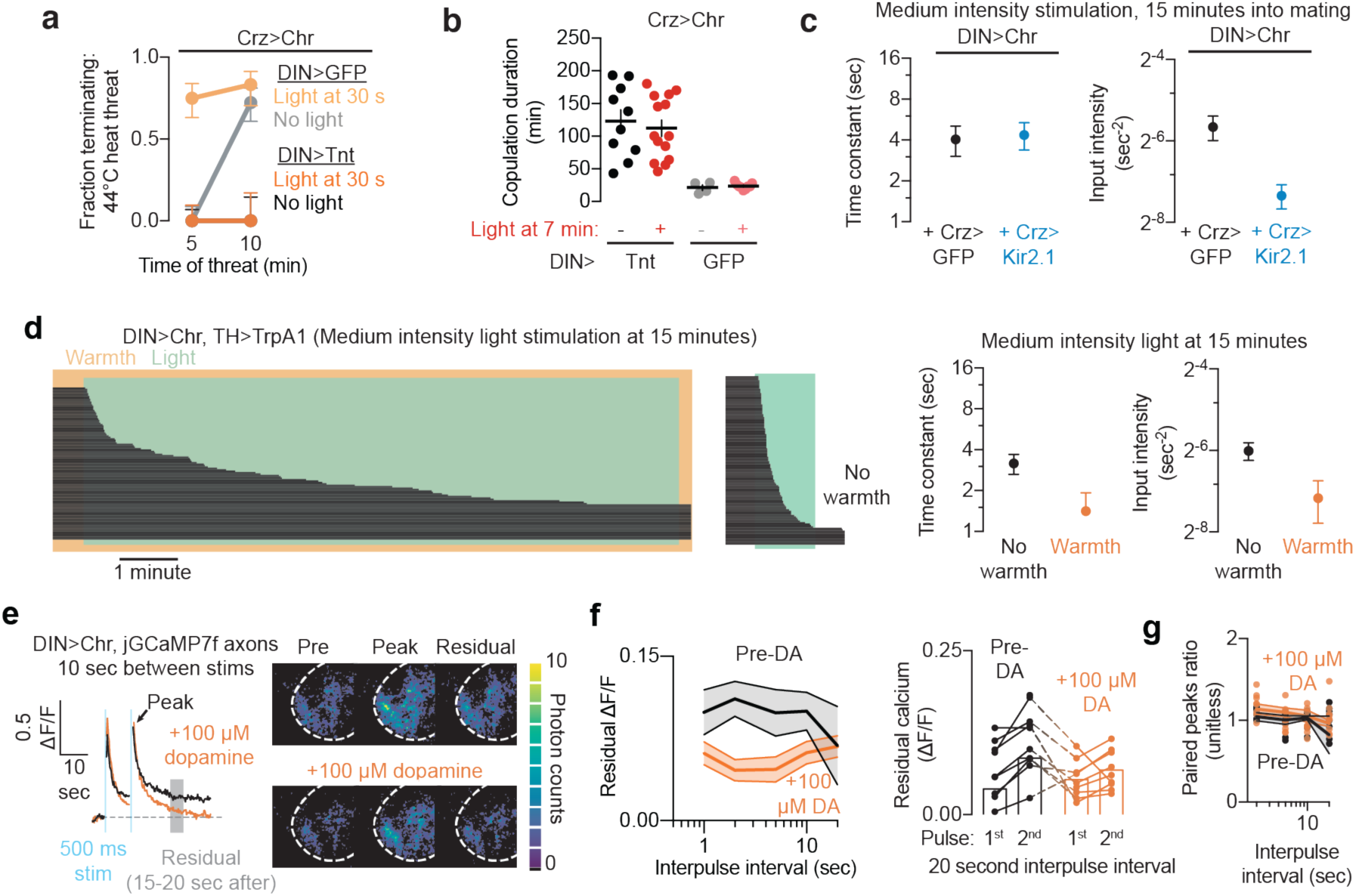
Motivating inputs increase persistence by restricting integration within the DINs. **a)** Silencing the DINs with tetanus toxin (Tnt) prevents Crz neuronal stimulation from reducing the motivation to sustain matings. **b)** Stimulating the Crz neurons does not prevent the long mating durations seen with DIN silencing. **c)** Silencing the Crz neurons reduces the response to sustained stimulation of the DINs by selectively decreasing the gain on the input (∼8-fold), leaving the time constant of integration largely unaffected. Stimulation used corresponds to the “medium intensity” condition in **Figure 2.** **d)** Far left: Sustained stimulation of the dopaminergic neurons with TrpA1 protects the mating against optogenetic stimulation of the DINs at 15 minutes into mating. Center left: Flies in which the dopaminergic neurons are not stimulated show little change in their response to optogenetic stimulation. Center right and far right: Dopamine reduces both the time constant of integration and the input intensity experienced by the DINs. Fitting is restricted to the first 60 seconds of optogenetic stimulation due to the habituation described in **Extended Data Figure 2b,c**. **e)** Application of dopamine results in more rapid clearance of calcium after optogenetic stimulation. (Left) Example traces of fluorescence summed across the imaged region. (Right) Absolute intensity of individual pixels (measured as number of photons detected within a 250 ms image acquisition) before (pre), immediately after (peak), and 20 seconds after (residual) the second optogenetic stimulation shown on the left. Dashed white line indicates approximate outline of the abdominal ganglion. **f)** Summary of residual calcium experiments as in **e**, shown for varying intervals between pulses. Measurements are taken as an average value between 15 and 20 seconds after the second pulse. (Left) Residual calcium following the second pulse was diminished by dopamine application (orange). (Right) Residual calcium is greater after the second pulse than after the first with a 20 s interpulse interval. In the absence of dopamine, the residual-calcium-mediated fluorescence of the second pulse is approximately double that of the first pulse (black), while after dopamine application the ratio is smaller (orange). **g)** The peak response of the second pulse is not affected by the application of dopamine.

Stimulating or silencing the dopaminergic neurons of the ventral nervous system (VNS) bidirectionally modulates the male’s propensity to sustain the mating in the face of threats^5^ (**Extended Data Figures 5f-h**). To ask whether dopaminergic activity alters the response to threats by acting on the DINs, as opposed to adjusting representations at earlier processing stages, we thermogenetically stimulated the dopamine neurons while providing weak and prolonged optogenetic DIN stimulation. Elevating dopaminergic activity at 15 minutes dramatically reduced the sensitivity to DIN stimulation, causing males to persist through several minutes of DIN stimulation that would normally cause termination after only a few seconds (**Figure 3d** and **Extended Data Figures 5i,l**). Fitting the cumulative distribution function revealed a 55% reduction in the time constant of integration, and a similar decrement in the intensity of input perceived by the DINs (**Figure 3d**). While our understanding of the neurons and signaling systems that motivate the male to sustain copulation remains incomplete, these results demonstrate that motivating inputs promote the stability of the ongoing behaviour by adjusting two properties of demotivating nodes: decreasing their accessibility to competing drives and decreasing the time over which inputs can integrate to promote behavioural switching.

Temporal integration could be implemented in many ways, but because direct optogenetic stimulation of the neurons was integrated across time, (**Figures 2c-e**), it seemed unlikely that the effect arises from changing dendritic responses to synaptic input. We therefore focused on axonal mechanisms by which the DINs could potentiate their response to inputs. Nearly all known presynaptic potentiation phenomena involve changes in axonal calcium levels during or after excitation^14^. We expressed the high-sensitivity fluorescent indicator jGCaMP7f^15^ in the DINs to measure changes in intracellular calcium in their axons after optogenetic stimulation. A single 500 ms pulse of excitation transiently evoked a large response in the neurons (**Extended Data Figure 5m**), but also resulted in a sustained increase in axonal calcium that persisted for tens of seconds, much longer than the off-kinetics of the reporter itself^15^. As predicted by a model in which residual calcium mediates the augmented behavioural response, sustained elevations of calcium were enhanced following a second stimulating pulse (**Figures 3e,f**), with no discernible effect on peak calcium (**Figure 3g**). Application of dopamine to the bath nearly abolished the sustained elevations in calcium after optogenetic excitation (**Figures 3e,f**), again consistent with a model in which motivating cues prevent integration at the DINs by expunging calcium. As has long been the case for synaptic augmentation on this timescale, the causes and consequences of this lingering calcium are difficult to test without knowledge of the mechanisms that regulate its removal. Nevertheless, these results point to residual calcium as a promising candidate for the shifting, memory-like effects of temporal integration in the demotivating neurons.

### The DINs synergistically pool demotivating inputs across modalities

Every threatening, damaging, or otherwise demotivating stimulus to which we have subjected a mating pair (short of forcible separation) requires the activity of the male’s DINs to elicit a termination response^5^. We sought optogenetically-tractable behaviours that could oppose copulation to test whether the principles derived from direct DIN stimulation apply to other drives. Stimulating neurons that drive grooming behaviour terminates mating with increasing efficacy as the mating progresses (**Figure 4a**), and required DIN activity to do so (**Figure 4a**). Grooming itself, whether induced by optogenetic stimulation (**Video 4**) or application of baking flour (**Video 5**), was suppressed during mating, but was initiated rapidly upon termination, showing that this paradigm resulted in a genuine competition between the two behaviours. Grooming behaviour showed the same characteristics of temporal integration as direct DIN stimulation (**Extended Data Figures 4c-g, Supplementary Discussion 1**). Since demotivating stimuli of each modality converge at the DINs, and since paired DIN stimulations produce a synergistic response greater than the sum of their independent probabilities, we predicted that multimodal competing inputs would combine to generate a stronger termination response than when delivered alone, or even than their independent sum, when delivered together. Confirming this prediction, combining optogenetic activation of the grooming neurons with a heat threat at 10 minutes into the mating (**Figure 4b, Video 6**) revealed termination probabilities greater than predicted if the two stimuli were acting independently.

**Figure 4:**
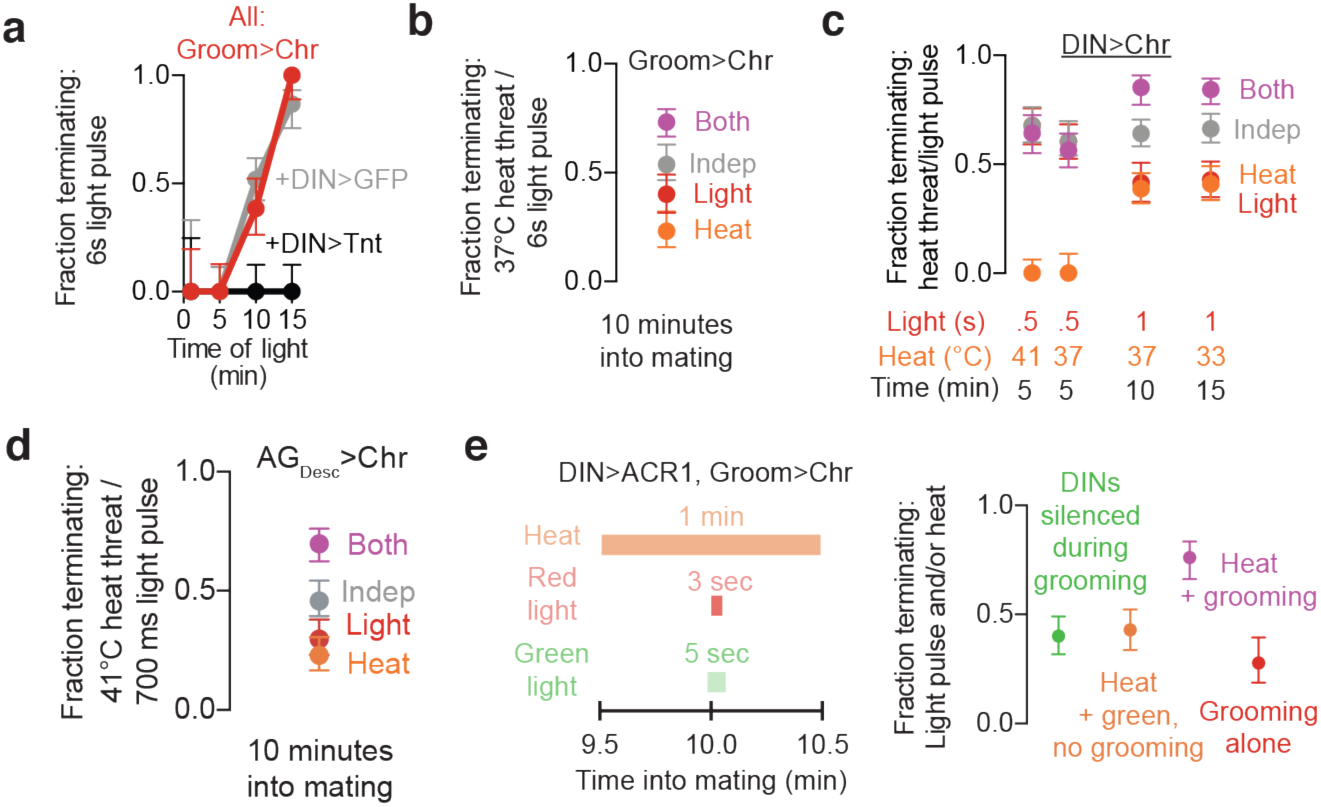
The DINs synergistically integrate inputs across drives, modalities, and time to demotivate copulation. **a)** Six seconds of optogenetic grooming neuron stimulation causes termination with increasing propensity as the mating progresses. Grooming-induced termination is prevented at all time points by blocking DIN output with tetanus toxin (Tnt). **b)** Stimulation of grooming neurons (6s; red) during a heat threat (60s; orange) results in a higher probability of terminating the mating (purple) than would be expected if the pathways did not interact (grey). **c)** Direct optogenetic stimulation of the DINs synergizes with heat threats to terminate mating, but only after the time of Crz activation (6 min into mating^6^), ruling out thermal effects on CsChrimson itself. **d)** Pairing optogenetic stimulation of AG_Desc_ with heat threats during mating increases termination probability more than would be expected if the two pathways did not interact. **e)** Silencing the DINs selectively during stimulation of the grooming neurons prevents any potentiation of the response to a threat, ruling out integrative effects upstream of the DINs.

We next performed a series of experiments pairing direct DIN stimulation with heat (**Figure 4c**). When heat and DIN stimulation were paired before the Crz-neuron mediated switch in motivation at ∼6 minutes into mating, we saw no contribution of a strong heat threat to termination probability (**Figure 4c**, also see **Supplementary Note 2**). This rules out any enhancing effect of temperature on CsChrimson activation itself^16^, and corroborates our finding that access to the DINs by real-world demotivating stimuli is blocked before the Crz switch is thrown. After the activity of the Crz neurons, we supplied heat and light stimuli that, when applied individually, terminate ∼50% of matings. At 10 minutes into mating, a 37°C threat gave a similar termination probability as a 33°C threat at 15 min (**Figure 4c**), reflecting the increasing responsiveness to real threats later in mating. Simultaneous presentation of real-world and optogenetic stimulation of the DINs at either 10 or 15 minutes caused higher termination probabilities than would be expected from their summed independent action (**Figure 4c**). These experiments show that competing inputs from diverse stimuli are pooled at the DINs, where they synergize to promote behavioural switching.

To extend these findings, we sought other impulses that could compete with copulation. We screened a collection of split-Gal4 lines that label small sets of neurons with cell bodies in the brain and that send projections to the VNS^7^ for lines that could oppose the motivation to copulate **(Extended Data Figure 4i).** The response to prolonged stimulation of the most effective of these, AG_Desc_ (for Abdomingal Ganglion-projecting Descending neurons) (**Extended Data Figure 4j,k**), showed augmentation over a time course that resembled that seen with the grooming neurons (**Extended Data Figure 4l**). As with grooming neuron stimulation, pairing excitation of the AG_Desc_ with heat threats resulted in a greater termination probability than could be explained by the two stimuli acting independently (**Figure 4d**).

These results point to drive integration at the DINs or elsewhere in the nervous system. To test integration by the DINs, themselves, we combined heat threats with brief grooming stimulation and silenced the DINs selectively during the stimulation of the grooming neurons (**Figure 4e**). This returned the termination probability to that of the heat threat alone (**Figure 4e**), arguing, together with the above results, that the integration of competing information occurs at the DINs.

### High-order interactions between drives can stabilize or destabilize the ongoing behaviour

The data presented above suggest three novel principles of motivational control over behavioral selection: i) synergistic effects of multimodal competing drive inputs on the ongoing behavior; ii) long-timescale integration of diverse inputs at behavior-specific demotivating neurons; and iii) that motivational cues prevent or limit integration of competing drive inputs. In this section we explore the implications of these principles assuming they hold across many or all behaviours.

We generated an integration-based mathematical model for high-order (i.e. supralinear) interactions between multiple drives. Drives are represented as evolving variables in a dynamical system using the principles described above, which we assessed both numerically and analytically (see Methods). The dominant (highest) drive is taken as the ongoing behaviour, which is switched if surpassed. Each drive has a demotivating node, like the DINs, that integrates inputs from all other drives using a fixed time constant (though the conclusions hold when τ is decreased by increasing motivation; **Extended Data Figures 7** and **8**). Excitation of the demotivating node is a monotonic function of the other drives, increasing as a single nonlinear function of a weighted sum of the inputs. Each drive, in isolation, acts in a consistent (linear) way, but changes in all drives synergistically impact all other drives. This is clearly an overly simplistic implementation, but it leads to interesting and testable predictions.

The model reproduces our experimental demonstration that synergistic integration of strong competing drives can produce behavioural switching in cases when merely additive effects would fall short (**Figure 5b, top row**, shown for several parameters in **Figure 5c**). Surprisingly, the model also predicts that weak competing drives often *enhance* the stability of the dominant drive in the face of a strong competitor (**Figure 5b, bottom row**). This occurs when the integrative effects of weak and dominant drives suppress a challenger more effectively than integration between the competing drives can suppress the dominant one. Importantly, this stabilizing effect follows from supralinear integration by other drives and is not predicted by supralinearity in only the dominant behaviour—that is, this stabilizing effect would likely not occur if DIN-like nodes do not exist for all relevant drives. These predictions hold across several complementary implementations: the numerical nonlinear dynamical system presented in **Figures 5b,c**, and **Extended Data Figure 9** (and examined analytically in the Methods); a rate-coding model of neurons in the presence of noise (**Extended Data Figure 7**) (showing robustness to noise and variability in exact wiring); and for ensembles of spiking neurons (**Extended Data Figure 8**).

**Figure 5:**
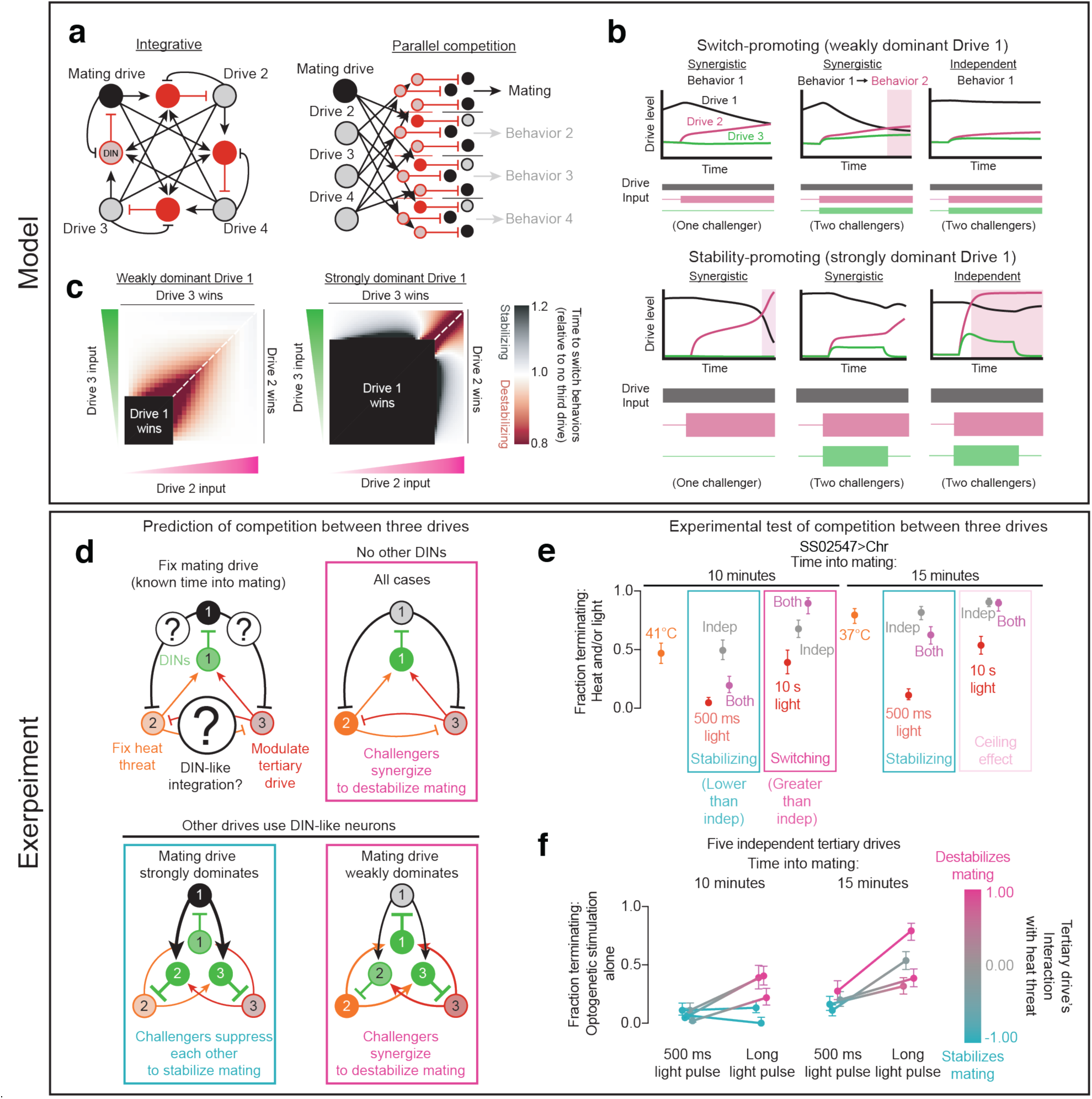
Nonlinear interactions between drives can predictably stabilize or destabilize the ongoing behaviour. **a)** Left: In an integrative circuit architecture, drive integrating neurons (red) pool information from all other drives (black). Motivating neurons prevent integration, while also stimulating the integrating neurons associated with competing behaviours. Right: Parallel computations would pit individual pairs of drives against each other, requiring a quadratically expanding number of nodes and connections as the number of drives increases, much greater than the linear growth of the integrative circuit. **b)** Synergistic integration can, depending on the full motivational state of the animal, either enhance or weaken the stability of the current behaviour. Top: A near-threshold stimulus exciting Drive 2 (magenta) is not capable of outcompeting a relatively weak dominant drive on its own (left), but synergistic integration with a weaker third drive (green) allows it to overcome the dominant drive (black) (middle panel), indicated by a change in the background colour. Independent action of the drives with the same stimulation would be less capable of weakening the dominant drive (right). Bottom: All-to-all synergistic integration prevents a challenger drive from eclipsing the dominant drive when paired with another weak drive. **c)** The same model as in **b**, with varying levels of input to the challenger drives. The time to switch behaviours (i.e. when Drive 2 overcomes Drive 1) is plotted as a function of the input to Drives 2 and 3. When the dominant drive is relatively weakly dominant (left; as in the top row of panel **b**), most interactions destabilize the dominant behaviour (blue regions). If the dominant behaviour is relatively strong (right), challenger drives may suppress each other more effectively than they cooperate to overcome the dominant drive (red regions). **d)** We test the predictions of different models of behavioural selection by putting three drives in conflict simultaneously: mating drive, the response to heat threats, and the response to optogenetic stimulation of tertiary drives. If the competing drives do not have corresponding drive-integrating neurons (top right), their synergy at the DINs should always destabilize mating. If the other drives have similar drive-integration neurons (numbered), synergistic interactions will show context-specific effects on mating drive (bottom row). **e)** As predicted by a DIN-based all-to-all model, tertiary drives can either stabilize or destabilize mating when confronted with a heat threat, depending on their relative intensity. A brief pulse of optogenetic stimulation stabilizes copulation behaviour (blue boxes), while increasing the stimulation intensity causes synergistic cooperation with the heat threat to oppose copulation (magenta box). A condition was labeled as “stabilizing” if the “both” condition showed a termination probability below the 95% credible interval of “independent” condition, “switching” if the termination probability was above the 95% credible interval, and “neither” if within the credible interval. **f)** These findings hold for nearly all lines tested and at either 10 (left) or 15 (right) minutes into mating. Each dot represents the strength of an optogenetic impulse on its own, and its colour indicates whether it stabilizes or destabilizes mating when presented in conjunction with a heat threat. Weak stimulation of individual lines can stabilize the ongoing behaviour (blue), but when the stimulation is strong enough to overpower the ongoing behaviour, synergistic effects with heat threats are observed (magenta). Raw data for all drives and explanation of “interaction” shown in **Extended Data Figure 10**.

To experimentally test the surprising prediction of stabilizing effects of weak tertiary drives (**Figure 5d**), we delivered heat threats during mating while also optogenetically activating the grooming neurons or the competing drive lines identified in **Extended Data Figure 4h,i**. 500 ms of optogenetic stimulation of these lines usually induced very low levels of termination and, as predicted, often caused lower-than-expected termination rates when paired with a heat threat (**Figure 5e,f** and **Extended Data Figure 10**). Remarkably, the combined terminating impact of heat and the tertiary drive was usually lower than the heat threat alone (e.g. **Figure 5e**, 500ms pulses). Though we emphasize that we do not know which drives are promoted when we stimulate most of these lines, the results are strikingly consistent with the prediction from the model, showing that a weak tertiary drive can dramatically decrease the effectiveness of a strong challenger (the heat threat). By increasing the duration of optogenetic stimulation, we found that if—and only if—increased stimulation turned these tertiary drives into strong challengers (i.e. they often overcame mating drive even when presented in isolation), they then synergized with heat to cause termination rates higher than would be expected from the independent action of the two stimuli (**Figure 5e,f** and **Extended Data Figure 10**). These results show that there is nothing intrinsically switch- or stability-promoting about individual competing drives; it is their relative intensities that determine their net influence on the ongoing behaviour. That this prediction was derived from assuming the generality of our findings across behaviours provides support for the wide applicability of our primary conclusion: that motivational control over decision-making is determined by integration at behaviour-specific demotivation neurons.

## Discussion

### The great parliament of instincts

Our results argue that Lorenz’s metaphor of a parliament of instincts may be useful beyond the immediate mental imagery it conjures: “it is a more or less complete system of interactions between many independent variables” in which “all imaginable interactions can take place between two impulses”^2^. Legislative decisions are not determined solely by the party with the largest representation, but often involve cooperation and antagonism between multiple factions whose allegiances may reverse with changes in context. Minority parties can be disruptive through the formation of coalitions or can reinforce the dominant party by uniting to suppress strong challengers.

Our proposed mechanistic instantiation of the parliamentary model for behavioural selection predicts that demotivating neurons like the DINs will be found to regulate many behaviours. The clearest analogs that we see in the literature are the parabrachial CGRP neurons of the mammalian hypothalamus. These neurons prevent feeding when activated, are stimulated by a wide variety of aversive cues, and are themselves suppressed by the AgRP neurons that motivate feeding behaviour^17–19^. Silencing the CGRP neurons leads to extended bouts of feeding^19^ arguing that they are required to demotivate feeding not just in response to competing drives, but also with the onset of satiety. Hunger is the most intensively studied mammalian motivation, and we expect that the ongoing circuit-level investigations into other behaviours will uncover demotivating nodes with converging inputs from many opposing drives.

### Temporal integration by demotivating neurons

Most adjustments to circuit functions over time and with experience have been found, or assumed, to result from changing synaptic weights. Here we show that a demotivating node operates over long motivational and decision-making timescales through a different mechanism for altering the response to fixed input: changing the time constant of integration. This mechanism has several theoretical advantages. For example, the representations of stimuli shorter than the timescale of integration are preserved, avoiding a potentiation of the response to noise and acting as a tunable low-pass filter.

While we cannot yet provide a detailed mechanistic description of the changing time constant of integration in the DINs, we find it useful to think about it in terms of the well-known phenomenon of synaptic augmentation^14^. Though not mechanistically understood itself, synaptic augmentation is thought to emerge from lingering calcium after an initial stimulus, generating a seconds-long period of increased transmitter release probability that decays as localized calcium is buffered or cleared. We also observe lingering calcium in the DINs, and find that the augmentation persists through electrical silencing, indicating that the memory-like trace is stored and adjusted biochemically. Synaptic augmentation lasts up to tens of seconds^20^, with a time constant that is independent of the strength of the initiating stimulus, also similar to the effects we observe here. In the DINs, augmentation is tuned by motivational inputs like dopamine to alter the impact of contemporaneous or long-lasting challenges as the male progresses through the mating. Targeted, functional genetic screening of the DINs will likely reveal the mechanisms that adjust this signal and implement its effects, information that may bring us to the verge of a thorough molecular explanation of motivation in this system.

## Author contributions

S.C.T. performed all experiments, with occasional assistance from M.A.C., and performed statistical and computational modeling. S.C.T. and M.A.C. designed the experiments, analyzed the data, and wrote the paper.a

## Acknowledgements

We thank: Dragana Rogulja for discussions, comments on the manuscript, and hosting us in her lab for the early stages of this project; Rachel Wilson’s lab for saline and advice on calcium imaging; Ryohei Yasuda and Long Yan for assistance building the two-photon microscope used in these experiments; Florian Engert for pointing us to similarities between parts of our model and an earlier proposal by Marvin Minsky; Ashna Singh, Megan Hoffman, Evan Zheng, Mary Dello Russo and Gabriel Verderame for assistance performing experiments; Barret Pfeiffer, David Anderson, and Gerry Rubin for sharing the UAS and LexAop2 Cs-Chrimson-tdTomato stocks before publication; Ofer Mazor and Pavel Gorelik (Harvard Medical School Neuroinstrumentation Core) for technical advice on designing the experimental apparatuses; and Jan Drugowitsch and members of the Crickmore and Rogulja Labs for comments on the manuscript. S.C.T. was supported by the National Science Foundation Graduate Research Fellowship (NSF Grant No. DGE1144152).

## References

1. Davis, W. J. Behavioural hierarchies. Trends Neurosci. 2, 5–7 (1979).

2. Lorenz, K. On Aggression. (Routledge Classics, 1966).

3. Burnett, C. J. et al. Hunger-Driven Motivational State Competition. Neuron 92, 187–201 (2016).

4. Russell, B. What Desires are Politically Important? (1950).

5. Crickmore, M. A. & Vosshall, L. B. Opposing dopaminergic and GABAergic neurons control the duration and persistence of copulation in Drosophila. Cell 155, 881–93 (2013).

6. Thornquist, S. C., Langer, K., Zhang, S. X., Rogulja, D. & Crickmore, M. A. CaMKII measures the passage of time to coordinate behavior and motivational state. Neuron (2020).

7. Namiki, S., Dickinson, M. H., Wong, A. M., Korff, W. & Card, G. M. The functional organization of descending sensory-motor pathways in Drosophila. Elife 1–50 (2018).

8. Baines, R. A., Uhler, J. P., Thompson, A., Sweeney, S. T. & Bate, M. Altered electrical properties in Drosophila neurons developing without synaptic transmission. J. Neurosci. 21, 1523–1531 (2001).

9. Mahn, M., Prigge, M., Ron, S., Levy, R. & Yizhar, O. Biophysical constraints of optogenetic inhibition at presynaptic terminals. Nat. Neurosci. 19, 554–6 (2016).

10. Klapoetke, N. C. et al. Independent optical excitation of distinct neural populations. Nat. Methods 11, 338–46 (2014).

11. Inagaki, H. K. et al. Optogenetic control of Drosophila using a red-shifted channelrhodopsin reveals experience-dependent influences on courtship. Nat. Methods 11, 325–32 (2014).

12. Veenstra, J. A. Isolation and structure of corazonin, a cardioactive peptide from the American cockroach. FEBS Lett. 250, 231–234 (1989).

13. Tayler, T. D., Pacheco, D. A., Hergarden, A. C., Murthy, M. & Anderson, D. J. A neuropeptide circuit that coordinates sperm transfer and copulation duration in Drosophila. Proc. Natl. Acad. Sci. U. S. A. 109, 20697–702 (2012).

14. Zucker, R. S. & Regehr, W. G. Short-Term Synaptic Plasticity. Annu. Rev. Physiol. 355–405 (2002). doi:10.1146/annurev.physiol.64.092501.114547

15. Dana, H. et al. High-performance GFP-based calcium indicators for imaging activity in neuronal populations and microcompartments. Nat. Methods 16, 434589 (2019).

16. Chater, T. E., Henley, J. M., Brown, J. T. & Randall, A. D. Voltage- and temperature-dependent gating of heterologously expressed channelrhodopsin-2. J. Neurosci. Methods 193, 7–13 (2010).

17. Campos, C. A., Bowen, A. J., Carolyn, W. & Palmiter, R. D. Encoding of danger by parabrachial CGRP neurons. Nature (2018). doi:10.1038/nature25511

18. Alhadeff, A. L. et al. A Neural Circuit for the Suppression of Pain by a Competing Need State. Cell 173, 140-152.e15 (2018).

19. Campos, C. A., Bowen, A. J., Schwartz, M. W. & Palmiter, R. D. Parabrachial CGRP Neurons Control Meal Termination. Cell Metab. 23, 811–820 (2016).

20. Magleby, B. Y. K. L. & Zengel, J. E. A quantitative description of tetanic and post-tetanic potentiation of transmitter release at the frog neuromuscular junction. J. Physiol. 183–208 (1975).

21. Lacin, H. et al. Neurotransmitter identity is acquired in a lineage-restricted manner in the Drosophila CNS. Elife 8, (2019).

22. Atkinson, J. & Birch, D. The Dynamics of Action. (1970).

23. Zhang, X. & Van Den Pol, A. N. Rapid binge-like eating and body weight gain driven by zona incerta GABA neuron activation. Science (80-.). 356, 853–859 (2017).

24. Liu, K. et al. Lhx6-positive GABA-releasing neurons of the zona incerta promote sleep. Nature 548, 582–587 (2017).

25. Zhao, Z. et al. Zona incerta GABAergic neurons integrate prey-related sensory signals and induce an appetitive drive to promote hunting. Nat. Neurosci. 22, 921–932 (2019).

26. Fadok, J. P. et al. A competitive inhibitory circuit for selection of active and passive fear responses. Nature 542, 96–99 (2017).

27. Han, W. et al. Integrated Control of Predatory Hunting by the Central Nucleus of the Amygdala. Cell 168, 311-324.e18 (2017).

28. Cai, H., Haubensak, W., Anthony, T. E. & Anderson, D. J. Central amygdala PKC-d+ neurons mediate the influence of multiple anorexigenic signals. Nat. Neurosci. 17, 1240–8 (2014).

29. Ma, C. et al. Sleep Regulation by Neurotensinergic Neurons in a Thalamo-Amygdala Circuit. Neuron 103, 323-334.e7 (2019).

30. Hong, W., Kim, D.-W. & Anderson, D. J. Antagonistic Control of Social versus Repetitive Self-Grooming Behaviors by Separable Amygdala Neuronal Subsets. Cell 158, 1348–1361 (2014).

31. Fadok, J. P., Markovic, M., Tovote, P. & Lüthi, A. New perspectives on central amygdala function. Curr. Opin. Neurobiol. 49, 141–147 (2018).

32. Allen, W. E. et al. Thirst regulates motivated behavior through modulation of brainwide neural population dynamics. Science (80-.). 364, (2019).

33. MacBean, I. T. & Parsons, P. A. Directional selection for duration of copulation in Drosophila melanogaster. Genetics 56, 233–239 (1967).

34. Nern, A., Pfeiffer, B. D. & Rubin, G. M. Optimized tools for multicolor stochastic labeling reveal diverse stereotyped cell arrangements in the fly visual system. Proc. Natl. Acad. Sci. U. S. A. 112, E2967–E2976 (2015).

35. Jaynes, E. T. Prior Probabilities. IEEE Trans. Syst. Sci. Cybern. 4, 227–241 (1968).

36. Jeffreys, H. An invariant form for the prior probability in estimation problems. Proc. R. Soc. Lond. A. Math. Phys. Sci. 453–461 (1945). doi:10.1098/rspa.1974.0120

37. Bender, C. M. & Orszag, S. A. Advanced Mathematical Methods for Scientists and Engineers: Asymptotic Methods and Perturbation Theory. (1999).

