## Supplementary Text and Figures for "Behavioural choice emerges from nonlinear all-to-all interactions between drives"

### SUPPLEMENTARY DISCUSSIONS

#### Supplementary Discussion 1

*Temporal integration when stimulating competing drives*

Like heat threats, a 6 second pulse of grooming stimulation terminated mating with increasing probability as mating progressed (**Figure 4a**). As expected, reducing the duration of the grooming stimulation to 3 seconds decreased the proportion of matings terminating at 10 or 15 minutes. However, unlike with a 6 second pulse, the response to 3 seconds of grooming stimulation was not much stronger when presented at 15 (compared to 10) minutes into mating (**Extended Data Figure 4c**). This result is consistent with the predictions of **Figure 2b**: only inputs lasting much longer than the time constant of integration become more effective as $\tau$ increases (**Extended Data Figure 4a**).

When we supplied paired 3-second stimulations of the grooming neurons 10 minutes into mating, we found no evidence for potentiation of the response to the second pulse regardless of the intervening time interval. At 15 minutes into mating, however, the two pulses interacted to produce a termination probability well above that predicted by their independent action (**Extended Data Figure 4d-f**). The enhanced response (which we refer to as augmentation by analogy to synaptic augmentation [14]) was roughly 2-fold if the pulses were separated by 5 seconds, but the effect was no longer clearly discernable if the inter-pulse interval was increased to 10 seconds. This augmentation was dependent on DIN activity, as silencing the DINs during the first pulse of grooming stimulation prevented the increased response to the second pulse (**Extended Data Figure 4g**). Silencing the DINs only during the interval between two grooming pulses did not affect integration (**Extended Data Figure 4h**), demonstrating that DIN voltage dynamics are required only for the receipt of demotivating input, not for its lingering effects.

The requirement for DIN activity not only for the response to demotivating stimuli, but also for the potentiation of a second stimulus indicates that the augmentation process occurs at or downstream of the DINs. Since DIN activity appears necessary for the response to all demotivating stimuli during mating, we expect these results to hold for any type of demotivating stimulus. The response to prolonged stimulation of the $AG_{Desc}$ (**Extended Data Figure 4f**), showed augmentation over a time course that resembled that seen with the grooming neurons and with the DINs (**Extended Data Figure 4b**). These results support and generalize the conclusion that the temporal window over which demotivating inputs accumulate to make behavioural decisions expands over time, using the same time constant for multiple forms of competing drives. This novel mechanism allows incremental adjustments of the response to incoming stimuli without otherwise altering how the stimuli themselves are represented and processed.

#### Supplementary Discussion 2

*Motivation in the ventral nervous system*

Many sensory and motor neurons relevant to copulation reside in, or project to, the abdominal ganglion, so only by comparison to the spinal cord is it surprising to find motivational circuit elements there as well. The spinal cord contains ~1% of human neurons, while the VNS contains ~20% of the fly's neurons[21]—and this percentage becomes much higher if the large optic lobes of the brain are not considered. We suggest that the presence of an exoskeleton in invertebrates may loosen neuronal packing constraints within the thorax, allowing, for example, ~1% of brain neurons to project to the VNS[7] compared to the <0.1% of mammalian brain neurons that descend to the spinal cord. We therefore believe that the invertebrate central nervous system should be considered as a functional continuum.

We are eager to identify the VNS neurons that are functionally downstream of the DINs for many reasons. One is practical: we are currently unable to monitor the activity of the DINs during mating because any invasive recording preparation would expose the male to extreme danger and cause termination. Silencing the downstream neurons that execute the termination of mating should remove this experimental constraint and allow a deeper understanding not just of the DINs themselves, but also of the motivating inputs such as the dopaminergic and Crz neurons that adjust the properties of the DINs to effect changes in decision making. Another reason is that we would like to know how the DINs effect the change in behaviour; we suspect the answer will be more than simply triggering a motor command. Computationally, we model the demotivating nodes as *de*-motivating: inhibiting the inputs that motivate the associated behaviour. This allows the dominant challenger to transition the animal into the next phase in the "continuous stream (of) change from one activity to another without pause from birth until death"[22].

*Resolution of conflicting results in other literature*

The parliamentary model may help explain several, apparently conflicting, recent studies of subcortical structures that receive converging inputs from all over the brain. Manipulating the same neuronal populations within these structures has different impacts on different behaviours, depending on the experimental context. For example the GABAergic neurons in the zona incerta control feeding[23], sleep[24], and hunting[25] and GABAergic neurons in the central amygdala drive defensive or fear behaviours[26], hunting[27], feeding[28], sleep[29], and more[30,31]. Some of this functional and contextual diversity may be explained if manipulating these populations shifts the relative intensities of latent drives—with indirect consequences for the tendency to perform the behaviour for which a motivating stimulus is experimentally provided. Similarly, the all-to-all network connectivity in our model may help explain why roughly half of the neurons across many brain regions respond to tasks designed to investigate only a tiny fraction of the animal's motivational, perceptual, and behavioural repertoires[32].

**EXTENDED DATA FIGURES**

#### Extended Data Figure 1

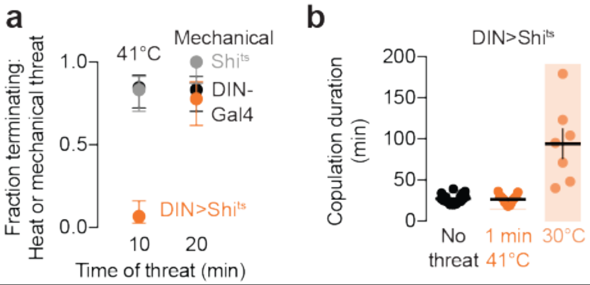

**Extended Data Figure 1 | Effects of silencing the DINs are not specific to GtACR1**
**a)** Expressing the temperature-sensitive synaptic silencing tool Shibire-ts ($Shi^{ts}$) in the DINs abolishes the response to heat threats (41°C heat), but mechanical stimulation (manipulating the vial in which the flies are mating) that does not activate the silencing effects of $Shi^{ts}$.can successfully cause termination
**b)** Acutely silencing the DINs with $Shi^{ts}$ during a heat threat does not affect overall copulation duration (middle column), but sustained silencing (right column) dramatically extends the mating.

#### Extended Data Figure 2

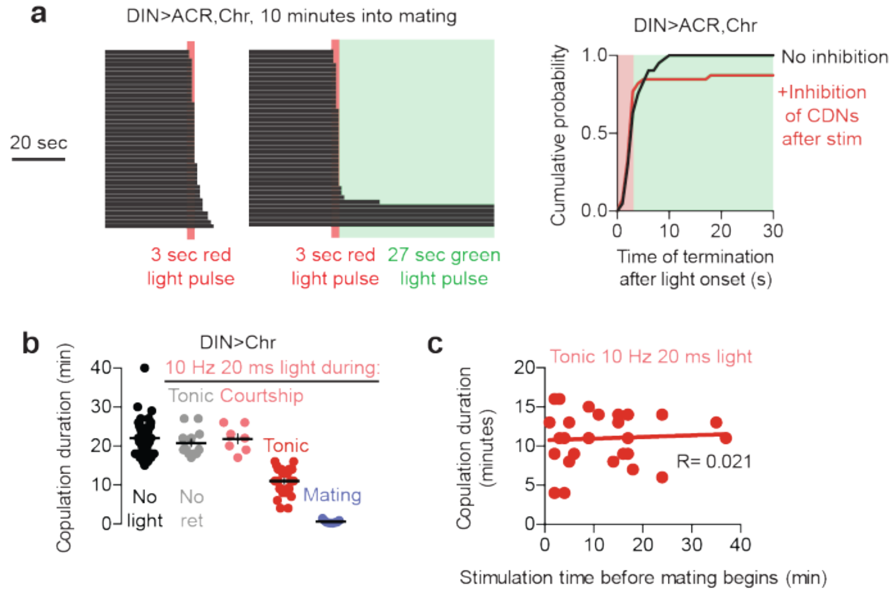

**Extended Data Figure 2 | Prolonged optogenetic stimulation of the DINs causes habituation, explaining previous thermogenetic results**

**a)** Stimulation of the DINs followed by immediate electrical silencing prevents the termination of mating during the period of silencing.

**b)** Optogenetic stimulation of the DINs using CsChrimson preceding the onset of mating does not affect copulation duration if only supplied during courtship (pink), but shortens copulation by several minutes if continued into the mating (i.e. flashing red lights throughout the duration of the experiment, “tonic”, red). These results closely resemble the results of thermogenetic activation in previous work that did not attribute immediate termination of the mating to DIN activation[5]. Providing the same optogenetic activation only after mating begins results in near-immediate termination of copulation (purple).

**c)** Habituation sets in within a few minutes of optogenetic stimulation, as there is no clear relationship between duration of optogenetic stimulation that induces habituation and the extent to which it shortens the mating.

**Extended Data Figure 3**

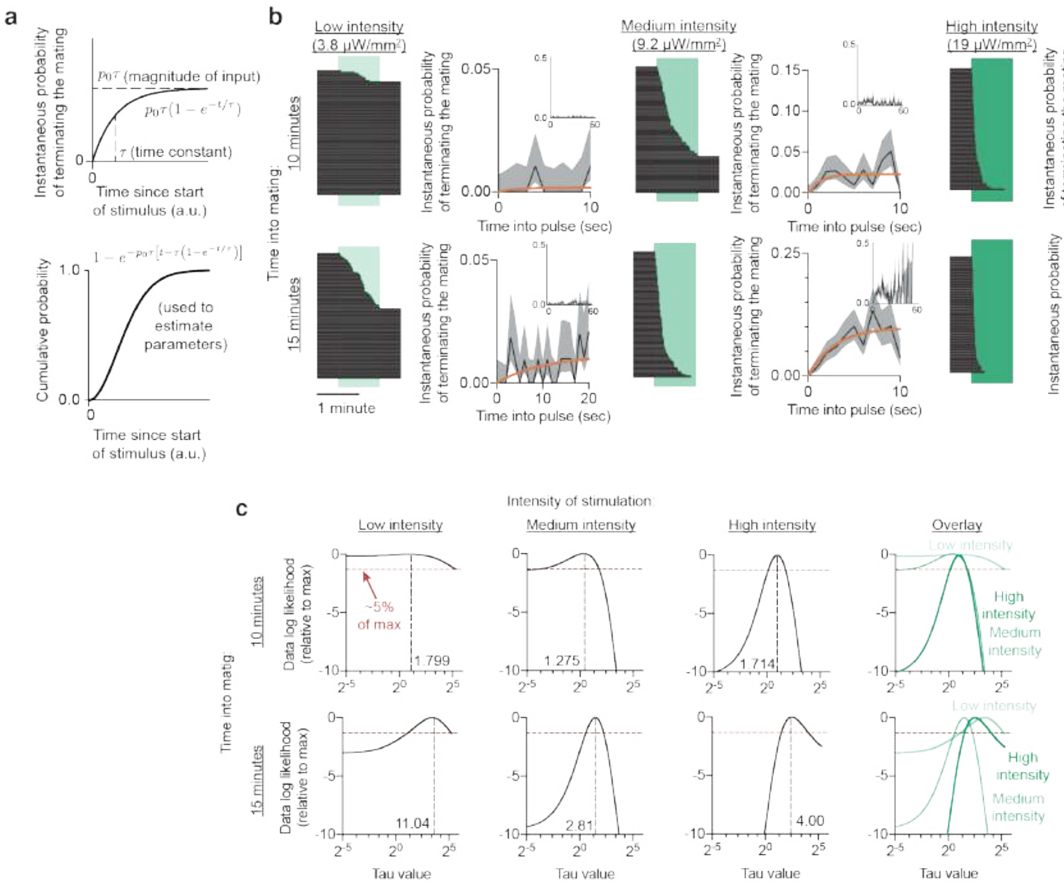

**Extended Data Figure 3 | Details of fitting the cumulative distribution in Figure 2**

**a)** If the instantaneous probability of terminating the mating in response to sustained stimulation ramps up as an exponential (left), then the cumulative probability of a mating ending by a particular time into a sustained stimulation follows the function $\sigma(t) = 1 - \exp\left(-p_0\tau\left(t - \tau\left(1 - \exp\left(-\frac{t}{\tau}\right)\right)\right)\right)$ (right).

**b)** Sustained optogenetic stimulation of the DINs results in an exponentially-ramping instantaneous termination probability that plateaus at values determined by the intensity of stimulation and time into mating. The horizontal black lines in the ethograms are individual matings. The orange lines in the instantaneous probability plots represent the maximum likelihood fit of the equation in **Figure 2c**. Insets show the entire 60 second pulse. The large fluctuations in instantaneous probability toward the end of highly demotivating pulses are caused by the small number of matings that endure to that point. Error bars on every point in the instantaneous probability plots (lumped into 1 second bins) are estimated independently from each other.

**c)** Likelihood of the observed data under the assumption of each $\tau$ (x-axis), assuming the maximum likelihood estimate for $p_0$ for that $\tau$, relative to the likelihood of the maximum likelihood estimate of $\tau$. These plots give a sense of how well our data is described by $\tau$ values other than the maximal one we present in **Figures 2d,e**. A log likelihood of $-1.301 = \log(0.05)$

corresponds to a $\tau$ value that would be 20 times less likely than the maximum likelihood estimate to produce the data we observed, if it were the true value.

**Extended Data Figure 4**

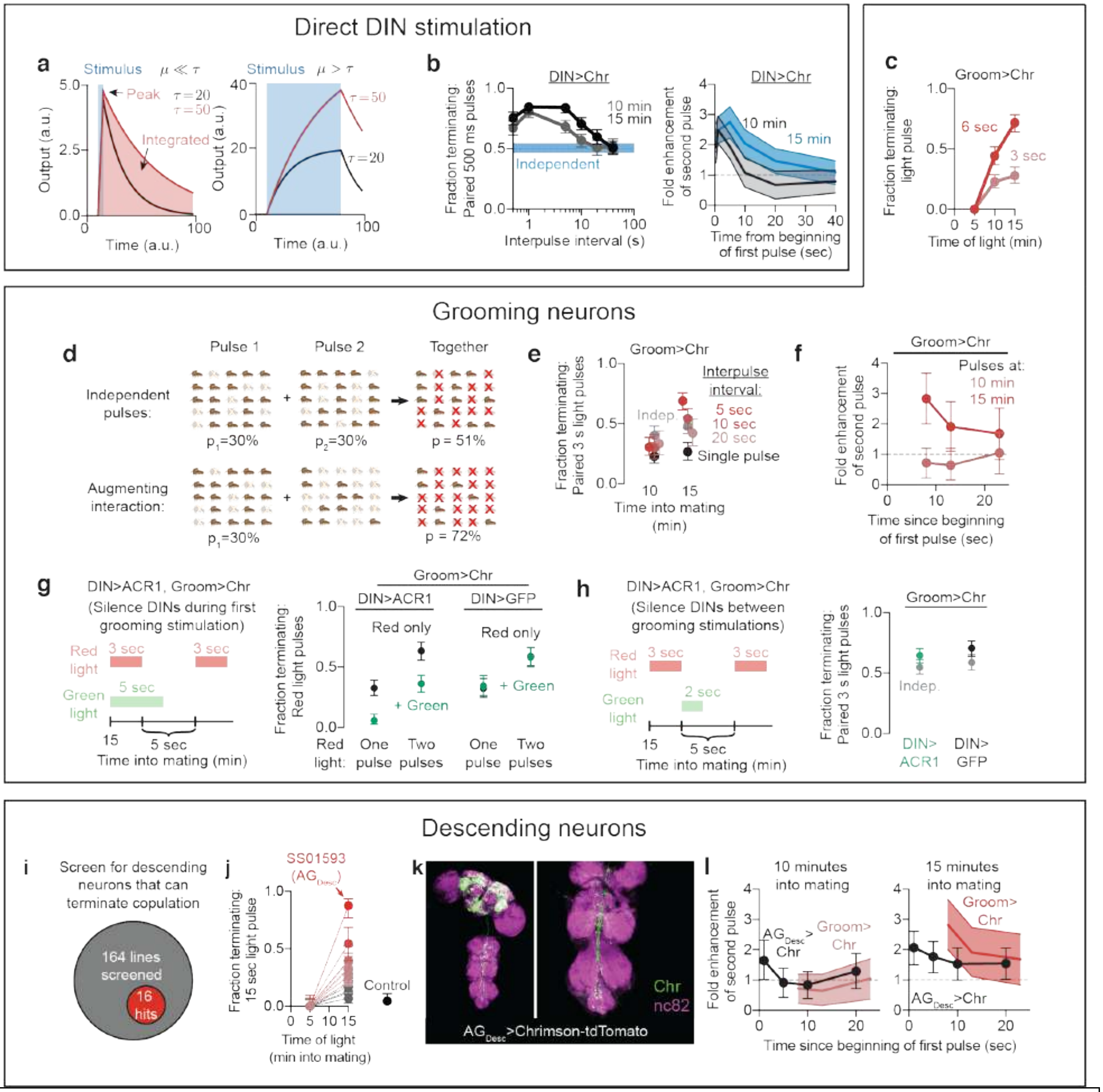

**Extended Data Figure 4 | The properties of temporal integration are the same across multiple demotivating impulses**

**a)** The model in **Figure 2b**, with the predictions for stimuli shorter than (left) and longer than (right) the timescale of integration.

**b)** A first pulse of optogenetic DIN stimulation potentiates the response to a second pulse, with an intervening timescale that increases with time into mating. Left: The probability of terminating the mating in response to two pulses of DIN stimulation is greater than expected if the pulses did not interact (estimated using the single pulse response value), with a longer interaction time at 15 minutes into mating than at 10 minutes. Right: The extent to, and timescale over, which the second pulse of DIN stimulation is potentiated.

**c)** Three seconds of grooming neuron stimulation terminates the mating less effectively than six seconds at 10 and 15 minutes into mating. The briefer (3 sec) demotivating challenge does not show a strong increase in potency when delivered later in mating.

**d)** Illustration of the probability of terminating the mating in response to two non-interacting (top) and synergistically interacting (bottom) pulses. In the non-interacting case, each pulse would terminate 30% of matings, and so the two acting together terminate 51% of matings (after subtracting the double-counted $9\% = 30\% \times 30\%$ that would have responded to either pulse). If the first pulse potentiates the second, then the overall proportion terminating the mating will be > 51% (in this example, 72% when the second pulse is augmented two-fold to 60%).

**e)** Two bouts of grooming neuron stimulation separated in time results in potentiation of the second pulse, but only if performed sufficiently close to the first pulse (5 seconds) and sufficiently late in mating (15 minutes). The predicted value of their independent interaction is indicated by a grey dot. Termination probabilities of the two-pulse experiment, coloured by time in between pulses.

**f)** Inferred potentiation of the second pulse, assuming that the first pulse is the same across treatments (i.e. single pulse responses were pooled between the 10 minute and 15 minute condition) from the data in **Extended Data Figure 4e** (red).

**g)** Silencing the DINs during a first demotivating stimulus prevents the potentiation of later stimuli

**h)** Silencing the DINs between two bouts of demotivating stimuli does not erase the memory of the first pulse or prevent the potentiation of later stimuli. The predicted values of two independent pulses of grooming stimulation are indicated by grey dots.

**i)** Of 164 descending-interneuron-labeling lines screened, 16 were able to terminate the mating when optogenetically stimulated.

**j)** 16 descending neuron lines showed varying efficacy in ending the mating when stimulated with a 15 second pulse of light at 15 minutes into mating. The control treatment is stimulation of DIN>GFP to control for the effect of the flash of light itself.

**k)** $Ag_{Desc}$ (SS01593), the most potent demotivating line from this collection, sends projections to the abdominal ganglion.

**l)** Paired pulses of $AG_{Desc}$ stimulation in rapid succession results in a greater termination probability than would be expected by their independent action, with a time course reminiscent of that over which grooming pulses are potentiated (red). Stimulation at 10 minutes into mating consisted of two 700 ms pulses of red light, while stimulation at 15 minutes was done with 300 ms pulses of red light.

#### Extended Data Figure 5

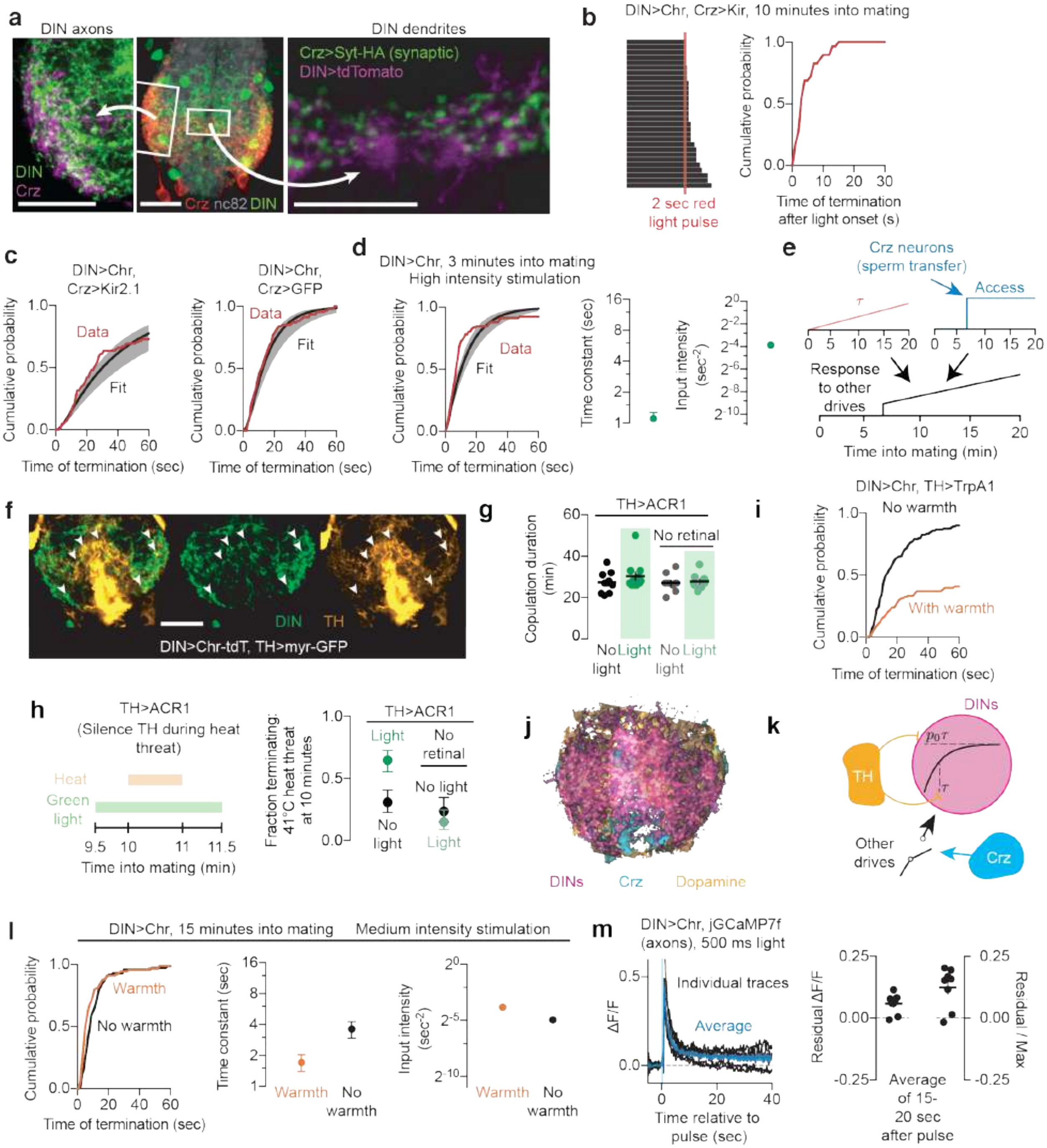

**Extended Data Figure 5 | Corazonin and dopamine releasing neurons differentially suppress temporal integration by the DINs**

**a)** The Crz neurons project throughout the abdominal ganglion, with processes closely apposed to those of the DINs, both near their axons (left) and dendrites (right), though synaptic connectivity cannot be concluded.

**b)** Optogenetic stimulation of the DINs while the Crz neurons are silenced results in termination of the mating, demonstrating that the DINs operate downstream of the Crz neurons in determining the motivational state of the fly.

**c)** Cumulative distribution functions used for estimating the parameters of **Figure 3c**.
**d)** The time constant of integration inferred from optogenetic stimulation of the DINs is shorter at 3 minutes into mating than at later times (compare to **Figure 2d**). High intensity green stimulation was used in these conditions to obtain a more reliable estimate of the anticipated shorter $\tau$, see **Extended Data Figure 6** for more details.
**e)** Model: The Crz neurons gate access to the DINs by permitting demotivating stimuli to access the DINs only after ~six minutes into mating. Before Crz activation, the input intensity to the DINs is so low that the male will persist through even lethal threats. After Crz activation, the persistence in mating continually decreases as a result of an expanding time constant of integration.
**f)** Dopaminergic neurons (orange) send projections throughout the abdominal ganglion, often forming varicosities near DIN (green) processes (indicated by white arrowheads). Images are obtained from a single optical plane.
**g)** Silencing the dopaminergic neurons does not affect overall copulation duration.
**h)** Acutely silencing the dopaminergic neurons enhances the response to threatening stimuli.
**i)** Cumulative distribution functions used for fitting the parameters in **Figure 3d**.
**j)** Overlaid maximum-intensity projections of the DINs (magenta), dopaminergic neurons (orange), and Crz neurons (cyan).
**k)** Schematized effects of the Crz and TH neurons on temporal integration by the DINs. Dopaminergic activity reduces both $\tau$ and $p_0$ to diminish the ability of competing drives to demotivate the mating. The Crz neurons permit competing drives to access the DINs by dramatically increasing the value of $p_0$ at ~6 minutes into mating.
**l)** Warmth alone decreases $\tau$ but increases $p_0$, showing that heat cannot account for the effects of stimulation the dopaminergic neurons.
**m)** Calcium levels within DIN axons remain elevated for tens of seconds after 500 ms optogenetic stimulation. (Left) Individual (black) and averaged (blue) response to a single pulse of optogenetic stimulation using CsChrimson. (Right) Almost all flies show long-lasting elevations in calcium corresponding to approximately 1/6 the observed maximal response to stimulation.

#### Extended Data Figure 6

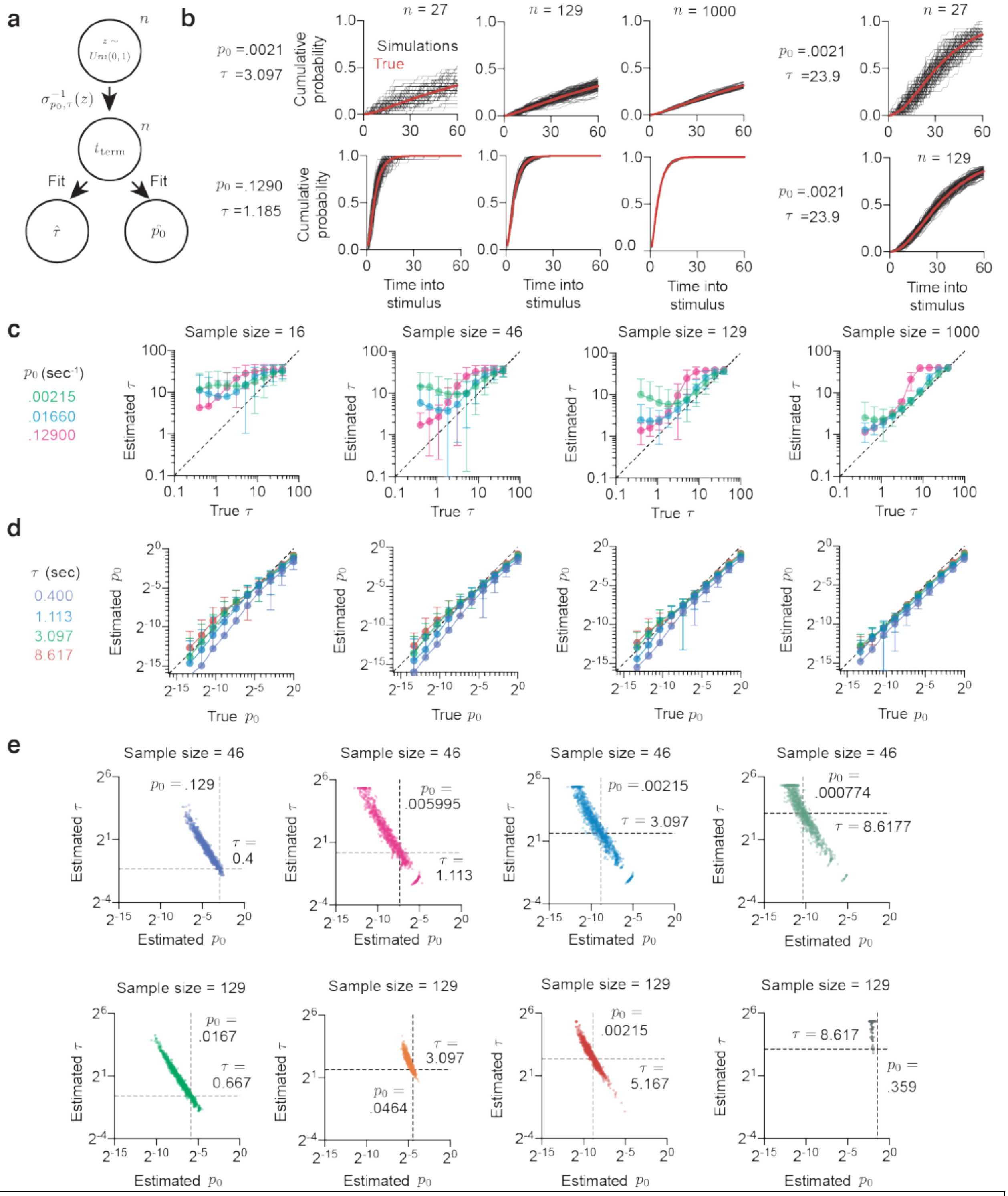

**Extended Data Figure 6 | Analysis of the methodology for fitting temporal integration and sensitivity to sample size**

**a)** Sampling scheme for generating data sets. $n$ samples were generated according to the given cumulative distribution function, $\sigma_{p_0,\tau}$, and these were used to fit estimates for the generating $\tau$

and $p_0$. The value of $n$ was varied logarithmically from 10 to 1000 to evaluate what sample size would be necessary to accurately estimate the parameters of the distribution.

**b)** The cumulative distribution function can be qualitatively reconstructed with samples of size ~ 100 across a wide range of cumulative distribution function shapes. Smaller sample sizes (e.g. ~30) are highly variable, especially when the overall number of flies terminating the mating during the stimulation is low (top row).

**c)** The sensitivity of the inference of the value of $\tau$ to sample size across a range of $p_0$ values. The closer a point is to the diagonal, the more likely the fitting procedure is to capture the correct $\tau$. The fitting procedure overestimates $\tau$ at low sample sizes, especially when the true value for $\tau$ is small. This may, to some extent, be explained by the fact that termination times are rounded to the nearest second (we find it is impossible to judge the time of termination more precisely than this value, given the complex motor sequence of terminating the mating). For larger sample sizes, the estimate is much better, so long as a large number of flies terminate the mating during the stimulation. When $p_0$ and $\tau$ are both small, however, the inference is considerably less reliable, because these conditions correspond to cases in which very few flies terminate the mating during the stimulus, providing very little information about $\tau$.

**d)** As in **c)**, but instead examining the sensitivity of the estimate of $p_0$. The parameter $p_0$ is easier to estimate, because even flies that do not terminate the mating during the stimulation are still informative about its value to some extent (see **Methods**). However, we find that $p_0$ is systematically underestimated due to the bias towards overestimating $\tau$ and the fact that the two estimates show substantial anticovariance (elaborated in **Extended Data Figure 6e**).

**e)** Covariance of $p_0$ and $\tau$ for various sample sizes. Dashed lines indicate the true parameter values, while independent points show individual sample estimates. The two parameters always anticovary, as indicated by the diagonal slant of each distribution. This reflects the fact that $p_0$ only appears in the cumulative distribution with $\tau$ in the form $p_0\tau$, and so this term is easier to fit than either value alone. Thus, if $\tau$ is overestimated, $p_0$ will tend to be underestimated to compensate. The multiplicative relationship is clear from the approximately linear covariance of the logarithm of the two parameters. When the data is more informative about $\tau$, i.e. many flies terminate the mating during the experiment, the cluster is much smaller (e.g. the orange data set). We therefore restricted our experiments to those conditions that would generate reliable estimates of the parameters, especially in cases where we expected $\tau$ or $p_0$ to be very small (e.g. **Extended Data Figure 5d**).

### Extended Data Figure 7

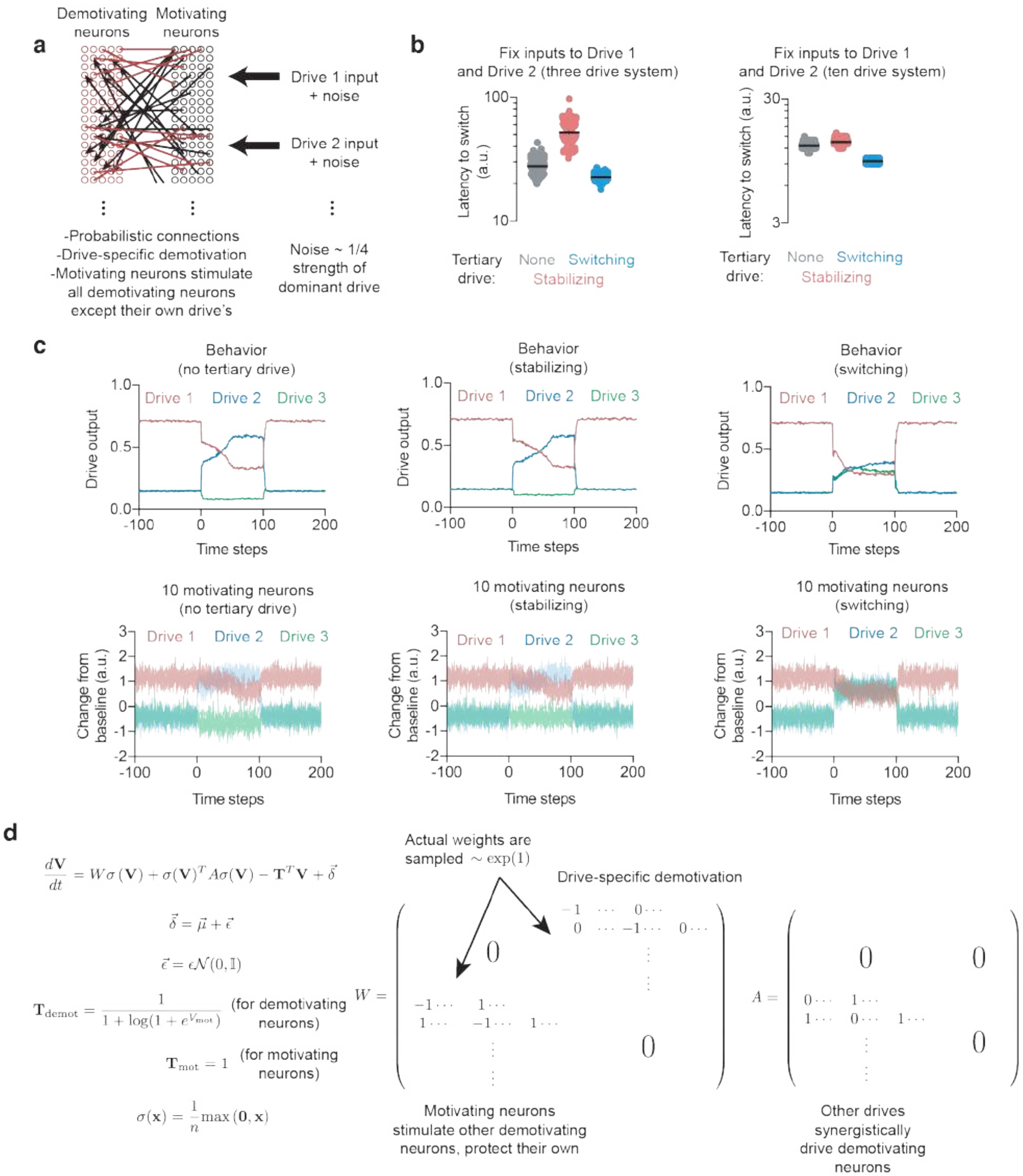

**Extended Data Figure 7 | Synergistic integration can destabilize or stabilize dominant drives in a probabilistically-connected linear rate-code network with noise**
**a)** The model is implemented as suggested in **Figure 5a**, with a population of demotivating and motivating neurons for each drive. Noise is on the scale of ~1/4 the magnitude of the input drives.

**b)** Left: In a three-drive system (as in **Figure 5**), the linear network recapitulates the stabilizing and switching regimes, with weak tertiary drives stabilizing the dominant behaviour (as measured by a longer time to switch behaviours) or destabilizing the dominant behaviour (as measured by a shortened time to switch behaviours). The values of Drive 1 and Drive 2 input are not adjusted across the three conditions. Right: Increasing the number of drives reveals the same phenomenon, though the system operates closer to the switching regime, because it constantly receives input from many drives.
**c)** Individual trials under the switching and stabilizing paradigms, in this instance using 3 drives. The top row represents the average drive output, normalized by a softmax function. The bottom row shows the firing rates of 10 motivating neurons for each drive.
**d)** Explicit mathematical form of the model. The linear weight matrix $W$ is sampled from an exponential distribution, while the synergistic matrix $A$ is fixed

.

#### Extended Data Figure 8

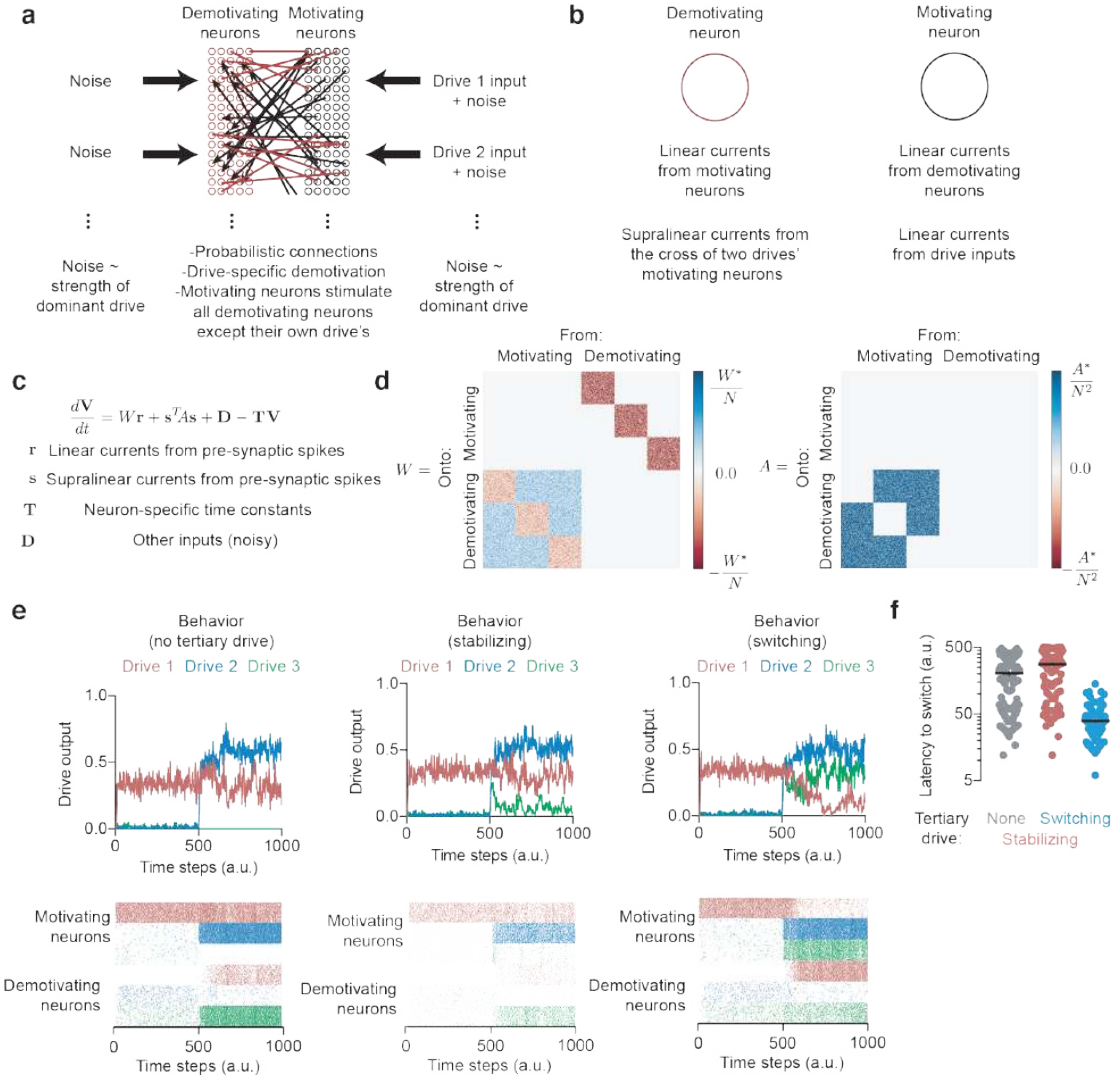

**Extended Data Figure 8 | Synergistic integration can destabilize or stabilize dominant drives in a spiking neural network**

**a)** The spiking neural network model is implemented like the linear rate-coding model, except the noise is of the order of the strength of the dominant drive to ensure substantive spontaneous activity.

**b)** Demotivating neurons experience linear and non-linear currents that sum to produce spikes when the voltage crosses a threshold, while motivating neurons receive only linear input currents.

**c)** The dynamics of the membrane voltage are determined entirely by the sum of the currents each neuron experiences.

**d)** Both the linear input current weights (represented by $W$) and the synergistic current weights (represented by $A$) are probabilistic, drawn from an exponential distribution and divided by either the total number of neurons (in the case of $W$) or the square of the total number of neurons (in the case of $A$).
**e)** A single example trial in each of the paradigms: no tertiary drive, stabilizing, and switching. Top row shows the average spike rate for motivating neurons in each drive. The bottom row shows spike rasters for all 600 neurons in the trial.
**f)** Latency to switch behaviours across 100 trials in each condition. All trials use the same average input for Drive 1 and Drive 2, and each condition fixes the average input to Drive 3 (but across each trial, each input has added Gaussian noise). The network can enter a stabilizing or destabilizing regime, dependent on the input to tertiary drives.

#### Extended Data Figure 9

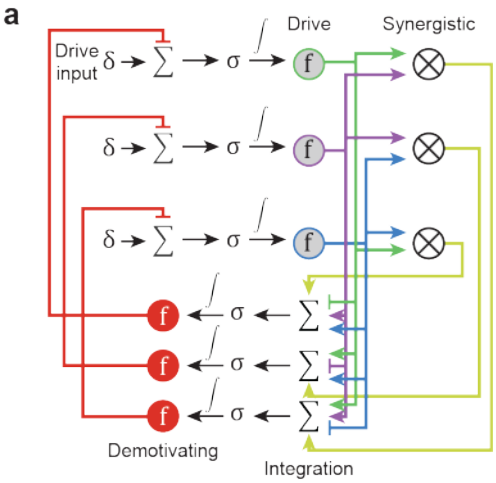

**Extended Data Figure 9 | Schematic of the nonlinear dynamical system implemented in Figure 5**

**a)** Each drive receives its own drive input, $\delta$, which is summed against inhibitory input (in red) and passed through a nonlinearity $\sigma$ to generate the drive value. Each drive value is passed through a synaptic nonlinearity $f$ to determine its input onto the demotivating neurons (in red). The input to the demotivating neurons is the sum of all opposing drives minus the value of each demotivating neuron's corresponding drive. In addition, each demotivating neuron receives a synergistic (circled X) term that is the product of the opposing drives. This input is passed through the same nonlinearity $\sigma$ to determine the level of demotivating inhibitory input onto each drive (which is passed through the synaptic nonlinearity $f$ to produce the inhibitory input experienced). In **Figure 5**, $\sigma$ is a sigmoid, while $f(x)$ takes the form $\log(a + b\exp(x))$ to prevent "negative" release from a synapse.

#### Extended Data Figure 10

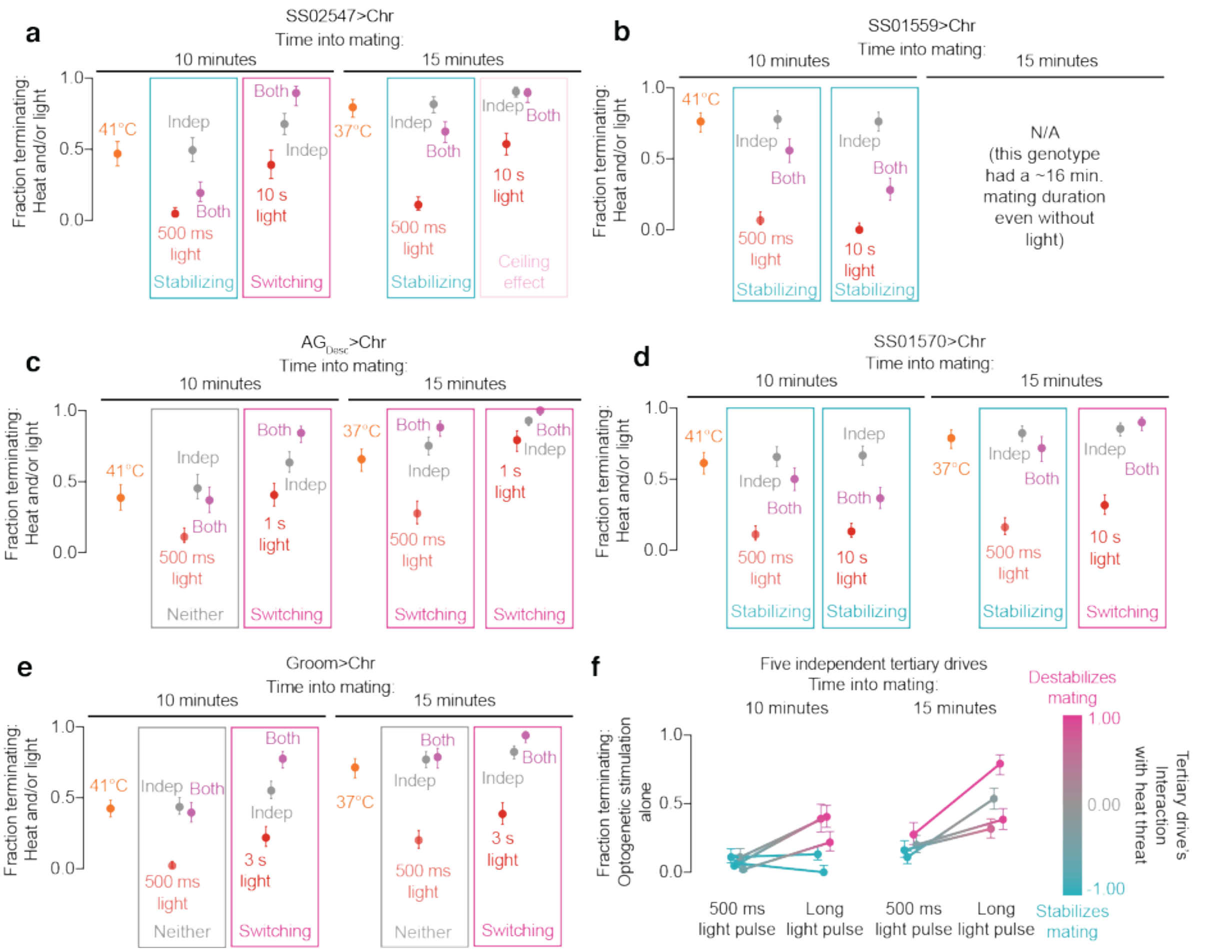

**Extended Data Figure 10 | Tertiary drives can either stabilize or destabilize the ongoing behaviour, depending on their relative intensities**

Note that panels **a** and **f** are reproduced from **Figure 5**.

**a-e)** By pairing weak or strong optogenetic stimulation of neurons labeled by challenger lines (red) with strong heat threats (orange), we find that challenger drives can either stabilize or destabilize mating when confronted with a heat threat, depending on their relative intensities. Weak tertiary drives stabilize copulation behaviour (blue boxes), while increasing the stimulation intensity of these same lines causes synergistic cooperation with the heat threat to oppose copulation (magenta boxes). These findings hold for multiple lines and at either 10 (left) or 15 (right) minutes into mating. The 15 minute condition for panel **b** was not tested because this line had a shorter-than-usual copulation duration. We did not find a stabilizing effect for the grooming neurons with 500 ms of stimulation in panel **e**. A condition was labeled as “stabilizing” if the “both” condition showed a termination probability below the 95% credible interval of “independent” condition, “switching” if the termination probability was above the 95% credible interval, and “neither” if within the credible interval.

**f)** Summary across all five genotypes within the figure. Each dot represents the strength of an optogenetic impulse on its own, and its colour indicates whether it stabilizes or destabilizes mating when presented in conjunction with a heat threat. Weak stimulation of individual driver lines can stabilize the ongoing behaviour (blue), but when the stimulation is strong enough to overpower the ongoing behaviour, synergistic effects with heat threats are observed (magenta). The "interaction" is computed from the p-value of an increase in termination probability as large as observed in the "both" condition, given that the "independent" condition is the true termination probability. The synergistic interaction $s$ is related to the p-value $p$ by $s = 1 - 2p$, so.that a highly synergistic interaction (where $p \approx 0$) results in $s \approx 1$ and a stabilizing interaction (where the termination probability is lower than nearly all independent cases, so that $p \approx 1$) results in $s \approx -1$.

#### VIDEO LEGENDS

**Video 1 | Silencing the DINs prevents the demotivation of mating in response to heat threats.**
Silencing the DINs with ACR1 beginning 14.5 minutes into mating prevents the male from terminating the mating in response to a one-minute 41°C heat threat beginning at 15 minutes after the onset of copulation (12 seconds into the video). The male does not appear obviously stuck or unable to disengage his genitalia, but is also not paralyzed as evidenced by his attempts to retain the mating posture even as the female struggles to escape.

**Video 2 | Stimulation of the DINs terminates the mating with variable latency.**
Several flies expressing CsChrimson in the DINs are exposed to 2 seconds of light after 10 minutes of copulation. Each fly responds with a variable latency, often several seconds after the optogenetic stimulation.

**Video 3 | The termination procedure in response to threats resembles that of stimulation of the DINs.**
A 41°C heat threat at 15 minutes into mating (beginning ~7 seconds into the video) results in termination behaviour resembling optogenetic stimulation of the DINs.

**Video 4 | Optogenetic stimulation of grooming neurons does not result in grooming behaviour during copulation.**
Sustained stimulation of the grooming neurons with CsChrimson (using the background white light) does not result in simultaneous grooming and mating, showing that grooming is actively suppressed by copulation. After the mating ends, the same fly grooms vigorously for several minutes, showing this is not due to desensitization of CsChrimson or insufficient activation of the neurons.

**Video 5 | Male flies suppress grooming during mating**
Wildtype flies sprinkled with baker's flour quickly clean themselves (see not mating flies), but a copulating male will suppress grooming (note the flour on the eyes). As soon as mating ends, he immediately cleans his eyes, even before his genitals; flies not exposed to flour clean their genitals immediately after mating.

**Video 6 | Heat and optogenetic stimulation of the grooming neurons synergize to demotivate copulation.**
18 successive matings (each well showing a different fly), aligned to the 9.5 minutes-into-mating timepoint, with either: 6 seconds of red light ~10 minutes into mating, 1 minute of 37°C heat beginning 9.5 minutes into mating (the beginning of the video), or both together. The threats synergize to demotivate copulation more effectively than if the two acted independently.

#### SUPPLEMENTARY NOTE 1

There is some ambiguity about the exact moment at which copulation ends, and so we round our data to the nearest second. This somewhat confounds the estimation of $\tau$, as it is far more challenging to confidently resolve values less than 1 second. When testing the robustness of the fitting algorithm in **Extended Data Figure 6**, all generated data was also rounded to the nearest second to emulate this effect. This results in an average estimate of $\approx$1 second even when $\tau$ is much smaller than 1 in **Extended Data Figure 6c**. However, if $\tau \geq 1$ second, the fitting algorithm reliably estimates appropriate values as long as the stimulus is not so strong that flies terminate mating immediately (i.e. $p_0$ is not too large).

#### SUPPLEMENTARY NOTE 2

In **Figure 4c** we use a 0.5 s pulse of DIN stimulation at 5 minutes to terminate ~50% of the matings, whereas later in mating we use 1 s pulses to achieve roughly the same effect. The increased sensitivity to intense stimulation early in mating is surprising since the response to real world threats increases over time and is essentially zero before ~6 minutes (see heat threat data in **Figure 4c**). This inconsistency is only apparent for brief red light pulses (intense stimulation), and not for sustained green light (weaker stimulation; compare to **Extended Data Figure 5d**). While we do not understand this discrepancy, the low input intensity perceived by the DINs before the Crz neurons signal at ~6 minutes (**Figure 5c**) likely means that the DINs never naturally receive intense stimulation very early in mating.

#### METHODS

##### Fly stocks

Flies were maintained on conventional cornmeal-agar-molasses medium under 12 hour light/12 hour dark cyclesxs at 25 °C. Unless otherwise stated, males were collected 0-6 days after eclosion and group-housed away from females for 3-6 days before testing. Flies expressing CsChrimson and all experimental controls for experiments involving CsChrimson were housed with rehydrated potato food (Carolina Bio Supply Formula 4-24 Instant Drosophila Medium, Blue) coated with all-trans-retinal (Sigma Aldrich R2500) diluted to 50 mM in ethanol for at least 3 days, unless marked as "no retinal." These vials were kept inside aluminum foil sheaths to prevent degradation of the retinal due to light exposure. Virgin females used as partners for copulation assays were generated by heat-shocking a UAS-CsChrimson-mVenus stock with a hs-hid transgene integrated on the Y-chromosome (Bloomington stock #55135) in a 37 °C water bath for 90 minutes. This stock was selected for mating partners because the females are highly receptive to courtship, resulting in a large number of mating pairs shortly after initiation of assays, and because it has been shown that copulation duration is robust to variations in the female's genetic background [33]. Virgins were group-housed for 3-13 days before use. Experiments with CsChrimson-expressing flies were not performed at specific times relative to the light-dark cycle of the incubator because these animals were housed in constant dark conditions to preserve all-trans-retinal integrity. We did not observe any dependency of time of day on any of the behaviours described here, but all flies experiencing light-dark cycles were tested between ZT (zeitgeber time) 2 and ZT 14 (lights are turned on at ZT 0 and off at ZT 12). Flies containing TH-LexA and LexAop-TrpA1 (or flies used as controls for TH>TrpA1 experiments) were raised at 19°C and collected as adults before being stored at 25°C. Detailed genotypes of all strains used in the paper are listed in the supplement.

##### Evaluation of mating

A pair of flies was scored as "mating" when they adopted a stereotyped mating posture for at least 30 seconds. This posture consists of the male mounting the female and propping himself up on her abdomen using his forelegs, while curling his own abdomen and keeping the genitalia in contact. The posture is starkly different from anything exhibited during other naturalistic behaviours and is rarely, if ever, sustained for 30 seconds during unsuccessful attempts to initiate a mating. If the flies are physically pulled apart without disengaging the genitalia (such as if the female falls or if they collide with an obstacle), the male is able to climb back into place. This posture is also maintained in the presence of threats, unless the male elects to terminate the mating differentiating it from a "stuck" phenotype. When stuck, the male dismounts the female, orients himself away from her, and attempts to walk away, but cannot decouple their genitalia. In the rare cases when we see a male become stuck in response to a threat, we record the copulation as having terminated. Occasionally in extremely long (>1hr) mating males, the male will become stuck, possibly because the seminal fluids harden and adhere the flies together. In this case too, the onset of the stuck posture is scored as the end of mating.

**Assessing fertility**

Mating pairs were manually separated by forcefully and repeatedly aspirating the flies through a narrow opening until the mating was forceably disrupted, and then the female fly was collected and placed in isolation in the above-described cornmeal food vials. One week later, the vial was visually inspected for the presence of larva as an indicator of successful fertilization. An exception to this is when the experiment involved DIN/NP2719>CsChrimson males, in which case the separation was performed by using a single two-second pulse of red light.

**Optogenetic stimulation during behaviour**

For CsChrimson experiments: One male and one virgin female fly was placed in each of .86" diameter 1/8" thick acrylic wells sitting 4" above 655 nm LEDs (Luxeon Rebel, Deep Red, LXM3-PD01-0350) driven using 700 mA constant current drivers (LuxDrive BuckPuck, 03021-D-E-700) and passed through frosted collimating optics (Carclo #10124). This spot of light was scattered using a thin diffuser film (Inventables, 23114-01) under the wells to ensure a uniform light intensity of ~0.1 $mW/mm^2$. The LEDs were controlled using an Arduino Mega2560 (Adafruit) running a custom script, which itself was controlled by a Raspberry Pi (either 2 or 3, running Raspbian, a Debian variant). Flies were observed by recording from above using the Raspberry Pi with a Raspberry Pi NoIR camera (Adafruit) and infrared illumination from below using IR LED arrays (Crazy Cart 48-LED CCTV Ir Infrared Night Vision Illuminator reflected off the bottom of the box) while streaming the video to a computer for observation.

For ACR experiments: The set-up was as above except using the green Luxeon Rebel, LXML-PM01-0100, and a pulse-width modulated signal to set the time-average intensity of the light to ~5 $\mu W/mm^2$ (approximately six times brighter than the ambient light) unless otherwise noted.

**Thermogenetics (with and without optogenetics) and heat threats**

A similar device to the one described above was constructed, with the addition of a ¼" thick water bath underneath each well. Room temperature water was continually passed through this bath, except when thermogenetic (TrpA1 or Shibire-ts) or heat threat manipulations occurred, when water of the temperature described in each experiment was used (controlled by a separate stopcock for each well). The LEDs above were driven with 1A BuckPucks controlled by a pulse-width modulated signal selected to ensure the average intensity of illumination is the same as in the other behavioural experiments (~0.1 mW/mm$^2$ for red light, ~ 5 μW/mm$^2$ for green) despite having to pass through the water bath.

**Additional notes about acute optogenetics and heat threat experiments**

In most DIN/NP2719>Chr experiments, flies which did not terminate in response to the light pulse were usually probed with a four-second pulse of light 30-60s after the original light pulse to ensure that they expressed Chrimson. In almost every mating pair, this was sufficient to induce termination, and in the few cases (<0.1%) in which it was insufficient, a subsequent 15 second pulse was likewise insufficient to terminate the mating, suggesting that these flies did not express CsChrimson (likely parental flies), and thus were not included in the reported data.

DIN/NP2719>Shibire-ts experiments were performed using only the second matings of flies whose first mating was of a normal duration (15-30 minutes). This was because a subset of males showed long (>1 hour) mating durations at room temperature, and so this protocol ensured that we only used flies in which expression of Shibire-ts did not induce any developmental defects that may have affected their response to heat threats. All control experiments in any plot containing DIN/NP2719>Shi$^{ts}$ were thus performed using the male fly's second mating. We did not observe any noticeable differences between the first and second matings in control flies, either in terms of heat threat responsiveness or copulation duration, and so express confidence that findings using the second mating are informative about the first mating.

All reported termination probabilities are technically conditional: they are only the subset of flies which persisted in mating until the noted time of the stimulus. For the 15-minute and earlier time points, this accounts for 100% of experimental flies, but data at the 20-minute time point should be considered in this light, rather than as a cumulative termination probability that includes the flies that terminated without intervention.

**Calcium imaging**

Images were acquired using a modified Thorlabs Bergamo II. Samples were excited using a Coherent Chameleon Vision II Ti:Sapphire laser emitting a 920 nm beam and emission was detected using cooled Hamamatsu H7422P-40 GaAsP photomultiplier tubes, with light collected through a 16x water immersion objective (Olympus). The PMT signal was amplified using Becker-Hickl fast PMT amplifiers (HFAC-26) and passed to a PicoQuant TimeHarp 260 photon counting board, which was synchronized to the laser emission by a photodiode (Thorlabs DET110A2) inverted using a fast inverter (Becker-Hickl A-PPI-D). The TimeHarp signal was acquired by custom software (FLIMage, Florida Lifetime Imaging) which was also used to control the microscope. For

intensity imaging, all detected photons within a pixel were summed together, regardless of arrival time relative to the excitation pulse.

Images were processed by simply taking the sum of all photon counts across the image, and then subtracting the background (estimated as the median pixel value, which was almost always 0 photon counts). Because our driver does not label any other neurons in the abdominal ganglion, we did not use ROIs or image segmentation to restrict our analysis to specific pixels. All data were likely entirely due to functioning GCaMP molecules, as opposed to aggregates of

GCaMP or autofluorescence, as their empirically-estimated fluorescence lifetime was always 2.6-2.8 nanoseconds. $\Delta F/F$ was computed as $\frac{\Delta F}{F}(t) = \frac{F(t)-F_0}{F_0}$ where $F_0 = \sum_{i=-6}^{-1} F(i)/6$ with $t = 0$ corresponding to the first frame of optogenetic excitation. Analysis was performed using custom Python3 code.

**Antibodies and immunohistochemistry**

All samples were fixed in PBS with Triton X-100 and 4% paraformaldehyde for 20 minutes, then washed three times with PBS/Triton X-100 for 20 minutes each before application of antibodies. All samples were incubated with the primary antibody for two days, washed three times with PBS/ITriton X-100 for 20 minutes each, incubated with the secondary antibody for two days, then washed three times as before and mounted on coverslips using VectaShield (Vector Labs). The exception is for MCFO staining, in which we followed the protocol of Nern *et al.*[34] Antibodies used are as follows:

Rabbit anti-GFP (A-11122, Invitrogen, 1:1,000 dilution)

Chicken anti-GFP (GFP-1010, Aves Labs, 1:1,000 dilution)

Mouse anti-GFP (A11120, Invitrogen, 1:2,000 dilution)

Rabbit anti-DsRed (323496, Clontech 1:1000 dilution)

Mouse anti-nc82 (Developmental Studies Hybridoma Bank)

Donkey anti-chicken 488 (703-545-155, Jackson ImmunoResearch, 1:400)

Donkey anti-rabbit 488 (A11008, Invitrogen, 1:400 dilution)

Donkey anti-mouse 488 (A21202, Invitrogen, 1:400 dilution)

Donkey anti-rabbit 555 (A-31572, Invitrogen, 1:400 dilution)

Donkey anti-mouse Cy3 (715-166-150, Jackson ImmunoResearch, 1:400 dilution)

Donkey anti-rabbit 647 (711-605-152, Jackson ImmunoResearch, 1:400 dilution)

Donkey anti-mouse 647 (711-605-151, Jackson ImmunoResearch, 1:400 dilution)

**Confocal microscopy**

Confocal images were collected using a Zeiss LSM 710 through a 20x air objective (Olympus PLAN-APOCHROMAT) controlled by Zen software, and analyzed using ImageJ.

**Statistics**

*General framework:* Throughout this manuscript, we take a Bayesian approach to parameter estimation because it more closely corresponds to the inference procedures we are performing, and as such all reported windows and intervals correspond to the mass of the posterior distribution for the inferred model parameter. This is because the Bayesian approach corresponds to inference about the values of descriptors of our model (e.g. in our data, the probability of terminating the mating in response to some stimulus), rather than consistency of a data set with a particular value that the model might take. With this approach, we can make statistical claims about our belief in the magnitude of effects, rather than simply reporting their deviation from that produced by a null hypothesis. We do, however, recognize that the frequentist approach is more commonplace, and so present our data in a manner that is as consistent as possible with typical frequentist reporting and hypothesis testing. We use noninformative priors [35], so this trivially corresponds to the usual Central Limit Theorem statistics in the case of estimating the variability of means, but a slightly different estimator on proportions. Our results and their interpretation do not hinge, in any case, on precise statistical methodology, as our effects tend to be very large, and so this decision is more philosophical than effectual.

*Credible intervals for proportions*: All proportions are modeled as Bernoulli random processes with probability $p$ and presented as the sample estimate $\hat{p}$ for the proportion $p$ ($\hat{p} = x/n$ with $x$ the number of observed successes and $n$ the total number of observations, the maximum likelihood estimate, rather than the maximum posterior estimate, for consistency with standard data presentation). This point is surrounded by a 68% credible interval (selected to be similar to the broadly familiar SEM metric, which is itself a 68% credible interval on the mean under a uniform prior), generated by sampling from the posterior distribution using Markov Chain Monte Carlo (MCMC, Metropolis-Hastings algorithm) with the noninformative Jeffreys prior $\pi(p) = 1/\sqrt{p(1-p)}$ and selecting the 16-84% window of this empirical estimate of the posterior. The Jeffreys prior was selected because it gives a posterior that is in a sense invariant under reparameterizations [36], and thus gives a consistent result between our posterior distributions even when we invert or transform the inference problem. The window generated by this method is also a numerical approximation of a 68% confidence interval with the corresponding frequentist properties.

The Bayesian framework becomes important when we perform inference on the distribution of probability of at least one of two Bernoulli random variables succeeding. If we know exact values for the probability of success of each random variable, the probability $p$ of at least one of the two, with probabilities $p_1$ and $p_2$ respectively, succeeding is simply $p = p_1 + p_2 - p_1 p_2$. But because we only have estimates of the true probabilities, we estimated the probability of at least one success by using our previously generated estimates of the posterior distributions of $p_1$ and $p_2$, and then sampling our estimate of the above quantity $p$ using MCMC. The credible intervals reported correspond to the 16-84% window of the estimated posterior distribution as above, with the reported sample estimate corresponding to $\hat{p} = \widehat{p_1} + \widehat{p_2} - \widehat{p_1 p_2}$. All code for these analyses (and corresponding hypothesis testing) was written using the freely available Python packages `numpy` and `pymc` and is available on Github at https://github.com/CrickmoreRoguljaLabs.

*Hypothesis testing on proportions*: To test the hypothesis that two sample proportions were drawn from the same Bernoulli process, we used Fisher's exact test. When testing whether a sample of size $N$ was consistent with two collections of independent draws from two separately estimated Bernoulli processes, we used a Monte Carlo approach to generate samples of size $N$ from randomly generated Bernoulli processes whose success probability $p$ was selected according to the posterior distribution described above. We then reported the fraction of samples whose number of successes was at least as high as that which we observed as the p-value (a one-tailed test that our experimental proportion was greater than would be expected if generated by the two independent Bernoulli processes estimated elsewhere).

*Credible intervals on means:* We use the standard SEM estimator for variability of sample means $SEM = \hat{\sigma}/\sqrt{N}$ with $\hat{\sigma}^2$ the unbiased estimator of sample variance and $N$ the sample size, which corresponds to the 68% credible interval for the sample mean using the "improper" uniform prior.

*Hypothesis testing on distributions:* We use the nonparametric Mann-Whitney U test on rank sums for differences in distributions of copulation duration. We then correct for multiple comparisons by using the Holm-Bonferroni correction on our criteria for statistical significance (with the number of hypotheses being the number of unique pairs of comparisons, $n(n-1)/2$).

**Modeling**

*General nonlinear dynamical system:*

The model in Figure 5 is formalized using a system of $2n$ differential equations. The first $n$ ($D_i$) correspond to the drive terms, while the second $n$ ($I_i$) correspond to the demotivating neurons:

$$\frac{dD_i}{dt} = \sigma[\delta_i(t) - f(I_i(t))] - D_i(t)$$

$$\frac{dI_i}{dt} = \sigma\left[\sum_j f\left(D_j(t)\right)\left(w_{ij} + \sum_{k\neq j,i} \alpha_{ijk} f(D_k(t))\right)\right] - I_i(t)/\tau_i$$

with $f(x)$ a monotonically increasing nonlinearity bounded below by 0 with $f(x) \to 0$ as $x \to -\infty$ and $\sigma(x)$ a nonlinearity bounded both above and below such that $\sigma(0) = 0$. Each $\delta_i(t)$ is any positive function of time. These correspond to a system of coupled "motivating" neurons, each of whose activity is given by $f(D_i)$ and whose activity increases when their inputs, $\delta_i(t)$, exceed their inhibition $I_i(t)$, and decrease when the opposite is true. The purpose of the negative limit of the nonlinearity $f(x)$ is to ensure no population's activity drops below 0. $\sigma(x)$ reflects the transformation from the strength of synaptic input into changes of activity of the neuron, and requiring that the function saturate imposes the condition that, when a neuron is already driven sufficiently forcefully, additional excitatory drive has minimal effect on the rate of increase in activity. The weights, $w_{ij}$, $\alpha_{ijk}$, and $\tau_i$, are nonnegative constants (or at least vary over timescales much slower than $D$ and $I$, as proposed in **Figure 2** – we do not account for $\tau_i$ changing as a function of $D_i$ in this analysis). This produces a model which, while fairly general, is not the only imaginable way to implement the system. However, we find that this model produces generally interesting phenomena which can be sought and tested experimentally. The parameter values in **Figure 5b and 5c** can found in **Supplementary Table 1**.

The argument of $\sigma$ in the equation for the inhibitory signals is motivated as the lowest-order Taylor expansion for any function of the drives $\{f(D_j(t))\}$ which increases "synergistically" as multiple drives co-vary. Setting the weights $\alpha_{ijk} = 0$ for all $i$, $j$, $k$ corresponds to "independent" (linear) integration. While the illustrations in **Figure 5** use the forms in **Supplementary Table 1** and are approximated using the `NDSolve` function in Mathematica, we prove a few results that are only weakly dependent on the forms of $f$ and $\sigma$ when the system operates in the regime in which the drives compete (i.e. are stimulated at levels comparable to their inhibition) by a series of approximations and linearizations.

First, we note that when one $\delta_i$ is sustained and much stronger than the others, and provided that the other $\delta_j$ are not too strong so that the inhibitory term $f(I_i)$ is drowned out, the system tends to an equilibrium. In this equilibrium, $D_j \approx \sigma\left[-\tau_j f\left(\sigma\left[w_{ji} f(\sigma[\delta_i])\right]\right)\right] < 0$ for all $j \neq i$, while $D_i \approx \sigma[\delta_i] > 0$, corresponding to pursuing drive $i$, as would be expected. Of course, when all drive inputs are strong enough to drown out their inhibitory input, the system converges to the equilibrium $D_i \approx \sigma[\delta_i]$ for all $i$. It is difficult to interpret this scenario, in which every drive is entertained at once.

Clearly the interesting dynamics occur when all of the drives are within the "physiological" range (i.e. within the dynamic range of $f$) and stimulated at somewhat comparable levels. In these conditions the argument of $\sigma$ in the equations for the drives is near zero, and we approximate it with the first order Taylor expansion $\sigma(x) \approx \sigma'(0)x$. We then have a simplified system of equations

$$\frac{dD_i}{dt} = g[\delta_i(t) - f(I_i(t))] - D_i(t)$$

$$\frac{dI_i}{dt} = \sigma\left[\sum_j f(D_j(t))\left(w_{ij} + \sum_{k \neq j,i} \alpha_{ijk} f(D_k(t))\right)\right] - I_i(t)/\tau_i$$

with $g = \sigma'(0)$. We make another simplifying assumption motivated by the findings in **Figure 2**, claiming that the inhibitory system "equilibrates" more slowly than the drive system (is quasi-static) so that $\{D_i\}$ is in equilibrium before $\{I_i\}$ has had time to change much, i.e. we treat $f(I_i(t))$ as approximately constant. This gives a simple linear equation for $dD_i/dt$ whose solution is $D_i(t) = e^{-t/\tau}\left(\int_0^t e^{t'/\tau} g[\delta_i(t') - f(I_i(t'))]dt' + D_i(0)\right)$. For our purposes, we even ignore much of the messy dynamics of this solution and opt for the equilibrium value of $D_i(t) \approx g[\delta_i(t) - f(I_i(t))]$. The question is really to determine the dynamics of $\{I_i\}$.

We now approximate $f(D_i(t))$ by $f_0 + \gamma[\delta_i(t) - f(I_i(t))]$ with $f_0 = f(0)$ and incorporate the coefficient in the first order Taylor expansion into $\gamma$ (i.e. $\gamma = \sigma'(0)f'(0)$). We additionally assert that $w = \sum_{j \neq i} w_{ij} \approx -w_{ii}$ and that $w_{ij} \approx w_{ik}$ when $j, k \neq i$ so that if all drives have equal input, they suppress each other to the same extent (or that ignoring the synergistic effects, each inhibitory input is driven by a value proportional to the average of the opposing drives). When one drive dominates, this average is a large value for every inhibitory population except for the dominant one, for which it is small.

We will use two approaches for approximating the solution to the system. The first examines explicit solutions after linearization but does not allow for detailed analysis of the system's behaviour, while the second employs a perturbative approach to introducing the synergistic term, but while the perturbative analysis permits more thorough analysis, it provides a less exact solution to the dynamics.

*Explicit solution in the pairwise case:*

In the absence of synergy, we have the equations

$$\frac{dI_i}{dt} = \sigma\left[w\gamma(\overline{(\delta - f)}_i - (\delta_i - f(I_i)))\right] - I_i(t)/\tau_i$$

where $\overline{(\delta - f)}_i$ is the average difference of input and inhibition to all of the alternative drive neurons $D_j$ with $j \neq i$. When many drive values are comparable, $\overline{(\delta - f)_i}$ is approximately the same for all $i$, and so we treat it as a constant. When there are only two populations with substantial input, each one experiences a $\overline{(\delta - f)}_i$ to which the other competing input predominantly contributes. All others experience a much stronger average input that forces their inhibitory terms up and thus suppresses the drive, and thus we only consider the competing drives. Using the same first order Taylor expansions around $\sigma(0)$ and $f(0)$, we obtain a solution for the linearized equation for $I_i(t)$

$$I_i(t) \approx \beta_i e^{-\zeta_i t}\left[\int_0^t dt' e^{\zeta_i t'}\left(\overline{(\delta - f)}_i - (\delta_i(t') - f_0)\right)\right]$$

with $\beta_i = \sigma'(0) f'(0) \sum_j w_{ij}$ and $\zeta_i = \beta_i + {}^1/_{\tau_i}$. Thus in order for the drive input $\delta_i$ to even briefly overcome the inhibition from the rest of the circuit,

$$\beta \int_0^t dt'\, e^{\zeta_i t'} \overline{(\delta - f)}_i + f_0 \le D_i(0) + \delta_i + \beta e^{-\zeta_i t}\int_0^t dt'\, e^{\zeta_i t'} \delta_i(t')$$

where the rightmost term generating an apparent boost in the ability to switch behaviours comes from the drive disinhibiting itself by weakening the other demotivating neurons.

If we reintroduce synergistic integration and return to the equation for $dI_i/dt$ in which $f(D_i)$ has been linearized we obtain

$$\frac{dI_i}{dt} = \sigma\left[\gamma\left(\overline{(\delta - f)}_i - \left(\delta_i - f(I_i)\right) + f_0\left[\sum_{j\neq i}\sum_{k\neq i,j}\alpha_{ijk}\left(f_0 + \gamma(\delta_k - f(I_k)\right)\right]\right.\right.$$
$$\left.\left. + \gamma^2\left[\sum_{j\neq i}\sum_{k\neq i,j}\alpha_{ijk}\left(\left(\delta_j - f(I_j)\right)\right)\left((\delta_k - f(I_k))\right)\right]\right)\right] - I_i(t)/\tau_i$$

We once again assume approximate balance to linearize $\sigma$ yet again and obtain

$$I_i(t) \approx \beta_i e^{-\zeta_i t}\left[\int_0^t dt' e^{\zeta_i t'}\left(\overline{(\delta - f)}_i - (\delta_i(t') - f_0) + f_0 \sum_{j\neq i}\sum_{k\neq i,j}\alpha_{ijk}\left(f_0 + \gamma(\delta_k - f_0 - f'(0)I_k\right)\right.\right.$$
$$\left.\left. + \gamma^2\left[\sum_{j\neq i}\sum_{k\neq i,j}\alpha_{ijk}\left(\left(\delta_j - f_0 - f'(0)I_j\right)\right)\left((\delta_k - f_0 - f'(0)I_k)\right)\right]\right)\right]$$

so our new inequality for switching behaviours is

$$\beta_i e^{-\zeta_i t}\left[\int_0^t dt' e^{\zeta\zeta_i}\left(\overline{(\delta - f)}_i + f_0 + f_0 \sum_{j\neq i}\sum_{k\neq i,j}\alpha_{ijk}\left(f_0 + \gamma(\delta_k - f_0 - f'(0)I_k\right)\right.\right.$$
$$\left.\left. + \gamma^2\left[\sum_{j\neq i}\sum_{k\neq i,j}\alpha_{ijk}\left(\left(\delta_j - f_0 - f'(0)I_j\right)\right)\left((\delta_k - f_0 - f'(0)I_k)\right)\right]\right)\right]$$
$$\le D_i(0) + \delta_i + \beta e^{-\zeta_i t}\int_0^t dt'\, e^{\zeta_i t'}\delta_i(t')$$

These additional terms promote switching in favor of behaviour $i$ when negative and stability when positive. If behaviour $i$ is relatively dominant, then $I_j$ and $I_k$ will have high values relative to the $\delta$'s and the system will favor behaviour $i$ even in cases that it would switch if the $\alpha_{ijk}$ were all 0. However, when the two are of the same order, the overall sign of the subtraction may switch and the system will make behaviour $i$ less stable than otherwise. Thus we see that synergistic integration promotes both stability and switching, dependent on the full complement of motivational inputs.

*Perturbative analysis:*

We rewrite the preceding equation in the notation of linear algebra (after some nondimensionalization to get $\boldsymbol{\delta}$ and $\boldsymbol{I}$ to have the same units, i.e. after multiplying by $f'(0)$ and absorbing it into the other constants) as

$$\frac{d\boldsymbol{I}}{dt} = \sigma\big(W(f(\boldsymbol{\delta}-\boldsymbol{I})) + \epsilon\, diag(Af(\boldsymbol{\delta}-\boldsymbol{I})f(\boldsymbol{\delta}-\boldsymbol{I})^T)\big) - \boldsymbol{I}/\tau$$

where $A$ is defined to be the matrix

$$A = \alpha \begin{pmatrix} 0 & 1 & 1 & \dots & 1 & 1 \\ 1 & 0 & 1 & 1 & \dots & 1 \\ 1 & 1 & 0 & 1 & \dots & 1 \\ \vdots & 1 & 1 & \ddots & \dots & 1 \\ 1 & \vdots & \vdots & 1 & \ddots & 1 \\ 1 & 1 & 1 & 1 & 1 & 0 \end{pmatrix} = \alpha(\boldsymbol{1}\boldsymbol{1}^T - \mathbb{I})$$

with $\mathbb{I}$ the identity matrix and $\alpha$ a constant (that may dependent on $n$) so that the diagonal of $Af(\boldsymbol{D})f(\boldsymbol{D})^T$ produces each of the synergistic terms and $W$ is defined to be the matrix of weights

$$W = \frac{w}{n}\begin{pmatrix} -(n-1) & 1\cdots & 1\cdots & 1 \\ 1\cdots & -(n-1) & 1\cdots & 1 \\ 1\cdots & 1\cdots & -(n-1) & 1 \\ 1\cdots & 1\cdots & 1 & -(n-1) \end{pmatrix} = \frac{w}{n}\left(\boldsymbol{1}\boldsymbol{1}^{\boldsymbol{T}} - n\mathbb{I}\right)$$

Now we assume the solution $\boldsymbol{I}(t)$ can be expressed as a perturbation series in the term $\epsilon$ describing the relative contribution of the synergistic term, approximating it as an infinite series of simplified linear cases[37]

$$\boldsymbol{I}(t) \sim \sum_{i=0}^{\infty} \epsilon^i \boldsymbol{I}_i(t) \quad (\epsilon \to 0)$$

and linearize the $\sigma$ and $f$ terms as before:

$$\frac{d\boldsymbol{I}}{dt} = \sigma'(0)\left(W\big(\boldsymbol{f}_0 + \gamma(\boldsymbol{\delta}-\boldsymbol{I})\big) + \epsilon\, diag\left(A\big(\boldsymbol{f}_0+\gamma(\boldsymbol{\delta}-\boldsymbol{I})\big)\big(\boldsymbol{f}_0+\gamma(\boldsymbol{\delta}-\boldsymbol{I})\big)^T\right)\right) - \boldsymbol{I}/\tau$$

We then group the solution series in order of $\epsilon$

$$\frac{d\boldsymbol{I}_0}{dt} = -Z\boldsymbol{I}_{\boldsymbol{0}} + \sigma'(0)W(\boldsymbol{f}_0 + \gamma\boldsymbol{\delta})$$

$$\frac{d\boldsymbol{I}_1}{dt} = -Z\boldsymbol{I}_{\boldsymbol{1}} + diag\big(A\big(\boldsymbol{f}_0 + 2\gamma(\boldsymbol{\delta}-\boldsymbol{I}_0)\big)(\boldsymbol{f}_0)^T\big)$$

$$\frac{d\boldsymbol{I}_2}{dt} = -Z\boldsymbol{I}_{\boldsymbol{2}} + diag[A(-2\boldsymbol{I}_1(\boldsymbol{f}_0+\gamma\boldsymbol{\delta})^T + \gamma^2\boldsymbol{I}_0\boldsymbol{I}_0^T)]$$

and generally

$$\frac{d\boldsymbol{I}_n}{dt} = -Z\boldsymbol{I}_n + diag\left[A\left(-2\gamma\boldsymbol{I}_{n-1}(\boldsymbol{f}_0+\gamma\boldsymbol{\delta})^T + \gamma^2\sum_{j=0}^{n-1}\boldsymbol{I}_j\boldsymbol{I}_{\boldsymbol{n-j-1}}^T\right)\right]$$

where $Z$ combines the linear dynamical terms

$$Z = \left(\frac{\mathbb{I}}{\tau} + \sigma'(0)\gamma W\right)$$

which highlights the need that $\sigma'(0)\gamma w < 1/\tau$, or else the system is unstable. Each of these relations is a driven linear dynamical system, with the drive corresponding to nonlinear terms that are lower orders in the expansion. We can then iteratively solve this system to arbitrary order.

The solution to equations of this form are

$$\boldsymbol{I}_n(t) = e^{-Zt}\int_o^t e^{Zs}\varphi_n(s)ds$$

where $\varphi_n(t)$ is the driving term for each equation. Only $\boldsymbol{I}_0(0)$ and $\boldsymbol{I}_1(0)$ have nontrivial initial conditions, so that

$$\boldsymbol{I}_0(t) = e^{-Zt}\boldsymbol{I}_0(0) + \sigma'(0)\, e^{-Zt}\int_o^t e^{Zs}W(\boldsymbol{f}_0 + \gamma\boldsymbol{\delta}(s))ds$$

The initial condition we will assume is that $\boldsymbol{I}_0(0)$ is at equilibrium, i.e.

$$Z\boldsymbol{I}_0(0) = \sigma'(0)W(\boldsymbol{f}_0 + \gamma\boldsymbol{\delta}(0))$$

or

$$\boldsymbol{I}_0(0) = Z^{-1}\sigma'(0)W(\boldsymbol{f}_0 + \gamma\boldsymbol{\delta}(0))$$

We may then use the Sherman-Morrison formula

$$(A+B)^{-1} = A^{-1} - \frac{1}{1+g}A^{-1}BA^{-1}$$

where $g = trace(BA^{-1})$ to compute $Z^{-1} = \left(\frac{\mathbb{I}}{\tau} + \sigma'(0)\gamma W\right)^{-1}$:

$$Z^{-1} = \tau\mathbb{I} - \frac{1}{1+\sigma'(0)\gamma\tau\, trace(W)}\tau\mathbb{I}\sigma'(0)\gamma W\tau\mathbb{I}$$

$$Z^{-1} = \tau\left(\mathbb{I} - \frac{W}{\frac{1}{\tau\sigma'(0)\gamma} - w(n-1)}\right)$$

so that

$$\boldsymbol{I}_0(0) = \sigma'(0)\tau\left(\mathbb{I} - \frac{W}{\frac{1}{\tau\sigma'(0)\gamma} - w(n-1)}\right)W(\boldsymbol{f}_0 + \gamma\boldsymbol{\delta}(0))$$

We then observe that $W\boldsymbol{f}_0 = 0$ to simplify this equation to

$$\boldsymbol{I}_0(0) = \tau\gamma\sigma'(0)\left(\mathbb{I} - \frac{W}{\frac{1}{\tau\gamma\sigma'(0)} - w(n-1)}\right)W\boldsymbol{\delta}(0)$$

which we further abbreviate by introducing the constants $\beta = \tau\sigma'(0)\gamma$ and $\chi_n = \frac{\beta}{1-\beta w(n-1)}$ to obtain

$$\boldsymbol{I}_0(0) = \beta(\mathbb{I} - \chi_n W)W\boldsymbol{\delta}(0)$$

We may perform a similar analysis to find the initial conditions of $\boldsymbol{I}_1$:

$$\boldsymbol{I}_1(0) = \tau(\mathbb{I} - \chi_n W)\, diag\left(A\left(\boldsymbol{f}_0 + 2\gamma\big(\boldsymbol{\delta}(0) - \boldsymbol{I}_0(0)\big)\right)(\boldsymbol{f}_0)^T\right)$$

$$\boldsymbol{I}_1(0) = \tau(\mathbb{I} - \chi_n W)\left(f_0^2(n-1)\boldsymbol{1} + 2f_0\left(diag(A(\boldsymbol{\delta}(0) - \boldsymbol{I}_0(0))\boldsymbol{1}^T)\right)\right)$$

Now we have the initial conditions and may analyze the dynamics of the system for arbitrary $\boldsymbol{\delta}(t)$.

We assume that one drive begins as dominant (without loss of generality, the first basis vector of $\boldsymbol{D}$) due to a greater value of $\boldsymbol{\delta}_1(0) = \Delta_0 + \Delta_1$ and that all others begin with a fixed drive value $\boldsymbol{\delta}_i(0) = \Delta_{0,}, 1 < i \leq\ n$. The initial conditions of this system are:

$$\boldsymbol{I}(0) = \frac{\beta w \Delta_1}{n}(1 + w\chi_n)(\mathbf{1} - n\boldsymbol{e}_1) + \epsilon f_0 \tau a \left[\left(\left(f_0(n-1) + 2\gamma\left(\Delta_0(n-1) + \beta w \Delta_1 (1 + w\chi_n)\left(n - 2 - \frac{1}{n}\right)\right)(1 + w\chi_n) - \frac{w\chi_n}{n}\Delta_1\right)\mathbf{1} - (1 + w\chi_n)\Delta_1\big(1 + \beta w(1 + w\chi_n)\big)\boldsymbol{e}_1\right]$$

where $\boldsymbol{e}_1$ is the first basis vector and the corresponding drive vector is

$$\boldsymbol{D}(0) = \boldsymbol{\delta}(0) - f_0 - \gamma \boldsymbol{I}(0)$$

This equilibrium goes to $\infty$ as $n$ grows unless $\epsilon a$ decays at least as fast as $1/n$, and so we will assume that to be true. We can discern a few interesting features of the integrative term (the one of order $\epsilon$) from this equilibrium. First, we see that there is a uniform increase to the inhibition experienced by each drive that scales with the dominance of the primary drive. This inhibition is diminished in the dominant drive by the nonlinear interaction term.

If the system then changes so that other drives are incremented by values $\{\Delta_j\}$ then we may analyze the dynamics of the system as described above. To do so, we first need to evaluate the matrix exponential

$$e^{Zt} = e^{\left(\frac{1}{\tau}\mathbb{I} + \sigma'(0)\gamma W\right)t} = e^{(1-\beta)t/\tau \mathbb{I}} e^{\frac{\beta w t}{n\tau}\mathbf{1}\mathbf{1}^{\boldsymbol{T}}}$$

Note that $\left(\mathbf{1}\mathbf{1}^{\boldsymbol{T}}\right)^k = \mathbf{1}\mathbf{1}^{\boldsymbol{T}}\mathbf{1}\mathbf{1}^{\boldsymbol{T}} \cdots \mathbf{1}\mathbf{1}^{\boldsymbol{T}} = \mathbf{1}n^{k-1}\mathbf{1}^{\boldsymbol{T}} = n^{k-1}\mathbf{1}\mathbf{1}^{\boldsymbol{T}}$ for $k \geq 1$. Then

$$e^{\frac{\beta w t}{n\tau}\mathbf{1}\mathbf{1}^{\boldsymbol{T}}} = \mathbb{I} + \sum_{k=1}^{\infty}\frac{\left(\frac{\beta w t}{n\tau}\mathbf{1}\mathbf{1}^T\right)^k}{k!} = \mathbb{I} + \frac{1}{n}\left(\sum_{k=0}^{\infty}\frac{(\beta w t/\tau)^k}{k!} - 1\right)\mathbf{1}\mathbf{1}^T = \mathbb{I} + \frac{(e^{\beta w t/\tau} - 1)}{n}\mathbf{1}\mathbf{1}^T$$

giving us our propagator

$$e^{Zt} = e^{(1-\beta w)t/\tau}\left(\frac{\left(e^{\beta w t/\tau} - 1\right)}{n}\mathbf{1}\mathbf{1}^T - \mathbb{I}\right) = e^{\frac{t}{\tau}}\left(\frac{1 - e^{-\beta w t/\tau}}{n}\mathbf{1}\mathbf{1}^T + e^{-\beta w t/\tau}\mathbb{I}\right)$$

We then find that the zero[th] order solution (noting that $\mathbf{1}^T W = 0$) is:

$$\boldsymbol{I}_0(t) = e^{-Zt}\boldsymbol{I}_0(0) + \sigma'(0)e^{-Zt}\int_0^t e^{(1-\beta w)t'/\tau} W\gamma\boldsymbol{\delta}(t')dt'$$

$$\boldsymbol{I}_0(t) = e^{-Zt}\boldsymbol{I}_0(0) + \sigma'(0)e^{-Zt}\sum_{j=1}^{n}\frac{\Delta_j w}{n}\left(\mathbf{1} - n\boldsymbol{e}_j\right)\int_0^t e^{(1-\beta w)s/\tau}ds$$

$$\boldsymbol{I}_0(t) = e^{-Zt}\boldsymbol{I}_0(0) + \frac{\sigma'(0)\gamma w\left(e^{(1-\beta w)t/\tau} - 1\right)\tau}{(1-\beta w)n}e^{-Zt}\sum_{j=1}^{n}\Delta_j\left(\mathbf{1} - n\boldsymbol{e}_j\right)$$

$$\boldsymbol{I}_0(t) = \frac{\beta w \Delta_1}{n}(1 + w\chi_n)e^{-(1-\beta w)t/\tau}(\mathbf{1} - n\boldsymbol{e}_1) + \frac{\beta w\left(1 - e^{-\frac{(1-\beta w)t}{\tau}}\right)}{(1-\beta w)n}\sum_{j=1}^{n}\Delta_j\left(\mathbf{1} - n\boldsymbol{e}_j\right)$$

Let's take a moment to discuss these dynamics, which are purely linear. The system moves along axes of the form $\mathbf{1} - n\boldsymbol{e}_j$, indicating a shared component (the $\mathbf{1}$) and a single insulating component for each drive $n\boldsymbol{e}_j$ ($\mathbf{1}$ and $\mathbf{1} - \boldsymbol{e}_j$ are eigenvectors of $Z$). The extent to which each drive is protected from the others is proportional to its input drive $\Delta_j$, as is the force it exerts on the rest of the system. As the system moves to the new equilibrium, each inhibitory population is active approximately at the level of $(\sum_k \Delta_k) - n\Delta_j$. If $\Delta_j$ is above the average of all drives, its inhibitory population is relatively quiescent. But all of the drives adjust themselves with the same overall dynamics: moving as a linear system with time constant $\tau/(1-\beta w)$.

The frequency with which we encounter these terms leads us to define $\boldsymbol{\xi}_j = \mathbf{1} - n\boldsymbol{e}_j$ so that we may describe $\boldsymbol{I}_0(t)$ in terms of $\{\boldsymbol{\xi}_j\}$:

$$\boldsymbol{I}_{\mathbf{0}}(t) = \sum_j I_0^j(t)\boldsymbol{\xi}_j = \frac{\beta w}{n}\left(\Delta_1(1+w\chi_n)e^{-\frac{(1-\beta w)}{\tau}t}\boldsymbol{\xi}_1 + \frac{1}{1-\beta w}\left(1-e^{-\frac{(1-\beta w)}{\tau}t}\right)\sum_j \Delta_j\,\boldsymbol{\xi}_j\right)$$

The terms $\boldsymbol{\xi}_j$ are not all linearly independent, with $\sum_j \boldsymbol{\xi}_j = 0$, so we add the vector $\boldsymbol{\xi}_0 = \mathbf{1}$, which we will call $\boldsymbol{\xi}_0$ and now $\{\boldsymbol{\xi}_j\} \cup \boldsymbol{\xi}_0$ forms a basis. These $\boldsymbol{\xi}_j$ serve as something of a proxy for the corresponding drive $D_j$, since they represent the extent to which each inhibitory population avoids some common input (the coefficient of $\boldsymbol{\xi}_j$).

The vectors $\boldsymbol{\xi}_j$ (when we use this notation, $j \neq 0$) have several convenient properties for our analysis. We note that that $A\boldsymbol{\xi}_j = -a\boldsymbol{\xi}_j, diag(\boldsymbol{\xi}_j\mathbf{1}^T) = \boldsymbol{\xi}_j, W\boldsymbol{\xi}_j = -w\boldsymbol{\xi}_j,$ and $Z\boldsymbol{\xi}_j = \frac{1-\beta w}{\tau}\boldsymbol{\xi}_j$. Also note that $Z\boldsymbol{\xi}_0 = \frac{1}{\tau}\boldsymbol{\xi}_0$

To change bases, we observe that $\boldsymbol{e}_j = \frac{1}{n}(\boldsymbol{\xi}_0 - \boldsymbol{\xi}_j)$ so that $\boldsymbol{\delta}(t) = \left(\Delta_0 + \frac{1}{n}\sum_j \Delta_j\right)\boldsymbol{\xi}_0 - \frac{1}{n}\sum_j \Delta_j\,\boldsymbol{\xi}_j$

The next step is to solve for the first order solution of the perturbative expansion:

$$\boldsymbol{I}_1(t) = e^{-Zt}\boldsymbol{I}_1(0) + e^{-Zt}\int_o^t e^{Zs} diag\left(A\left(\boldsymbol{f}_0 + 2\gamma(\boldsymbol{\delta} - \boldsymbol{I}_0(s))\right)(\boldsymbol{f}_0)^T\right)ds$$

We first evaluate the driving force

$$diag\left(A\left(\boldsymbol{f}_0 + 2\gamma(\boldsymbol{\delta} - \boldsymbol{I}_0(t))\right)(\boldsymbol{f}_0)^T\right)$$
$$= f_0 a\left((n-1)\left(f_0 + 2\gamma\left(\Delta_0 + \frac{1}{n}\sum\nolimits_j \Delta_j\right)\right)\boldsymbol{\xi}_0 + \frac{1}{n}\sum\nolimits_j (\Delta_j + nI_0^j(t))\boldsymbol{\xi}_j\right)$$

Then we apply our propagator

$$e^{Zt}\varphi_1(t) = f_0 a\left((n-1)\left(f_0 + 2\gamma\left(\Delta_0 + \frac{1}{n}\sum\nolimits_j \Delta_j\right)\right)e^{t/\tau}\boldsymbol{\xi}_0 + \frac{1}{n}e^{\frac{(1-\beta w)}{\tau}t}\sum\nolimits_j (\Delta_j + nI_0^j(t))\boldsymbol{\xi}_j\right)$$

Performing the integral yields

$$\int_0^t e^{Zs}\varphi_1(s)ds = f_0 a\tau\left((n-1)\left(f_0 + 2\gamma\left(\Delta_0 + \frac{1}{n}\sum\nolimits_j \Delta_j\right)\right)\left(e^{\frac{t}{\tau}} - 1\right)\boldsymbol{\xi}_0\right.$$
$$\left. + \frac{1}{n}\sum\nolimits_j\left(\frac{\Delta_j}{1-\beta w}\left(e^{\frac{(1-\beta w)}{\tau}t} - 1\right) + \frac{n}{\tau}\int_0^t e^{\frac{(1-\beta w)}{\tau}s} I_0^j(s)\,ds\right)\boldsymbol{\xi}_j\right)$$

$$= f_0 a\tau\left((n-1)\left(f_0 + 2\gamma\left(\Delta_0 + \frac{1}{n}\sum_j \Delta_j\right)\right)\left(e^{t/\tau}-1\right)\boldsymbol{\xi}_0 + \frac{1}{n}\sum_j\left(\frac{\Delta_j}{1-\beta w}\left((1+\beta w)\left(e^{\frac{1-\beta w}{\tau}t}-1\right)+\frac{\beta w}{\tau}t\left(\delta_{j,1}-1\right)\right)\right)\boldsymbol{\xi}_j\right)$$

Finally we apply $e^{-Zt}$ to get

$$\boldsymbol{I}_1(t) = e^{-Zt}\boldsymbol{I}_1(0) + f_0 a\tau\left((n-1)\left(f_0 + 2\gamma\left(\Delta_0 + \frac{1}{n}\sum_j \Delta_j\right)\right)\left(1-e^{-t/\tau}\right)\boldsymbol{\xi}_0 + \frac{1}{n}\sum_j\left(\frac{\Delta_j}{1-\beta w}\left((1+\beta w)\left(1-e^{-\frac{1-\beta w}{\tau}t}\right)+\frac{\beta w}{\tau}te^{-\frac{1-\beta w}{\tau}t}\left(\delta_{j1}-1\right)\right)\right)\boldsymbol{\xi}_j\right)$$

where $\delta_{xy}$ is the Kronecker delta function, which is 1 when $x = y$ and 0 otherwise. This solution is somewhat more interesting due to the transient term $te^{-\frac{1-\beta w}{\tau}t}$ inhibiting the non-dominant drives, but still does not show mixing terms of the form $\Delta_j\boldsymbol{\xi}_k$, and so does not reveal how interactions between drives contribute to the system's dynamics. However the transient term is important in that it contributes substantially to the solution early on, when $t < \frac{\tau}{1-\beta w}$. Understanding these interactions requires going to the second order perturbative solution, where the linear zero$^{\text{th}}$ order system has had a chance to move through two layers of processing and the effects of other drives can begin to mix. This term is given by

$$\boldsymbol{I}_2(t) = e^{-Zt}\int_0^t e^{Zs} diag\left[A\left(-2\boldsymbol{I}_1(s)(\boldsymbol{f}_0+\gamma\boldsymbol{\delta})^T + \gamma^2\boldsymbol{I}_0(s)\boldsymbol{I}_0^T(s)\right)\right]ds = e^{-Zt}\int_0^t e^{Zs}\varphi_2(s)ds$$

where the $A$ term forces mixing between the drives. $diag\left(\boldsymbol{\xi}_j\boldsymbol{\xi}_0^T\right) = \boldsymbol{\xi}_j$ and $diag\left(\boldsymbol{\xi}_j\boldsymbol{\xi}_k^T\right) = \boldsymbol{\xi}_j + \boldsymbol{\xi}_k - \boldsymbol{\xi}_0 + n^2\delta_{jk}$, so we can plug in the above formulae to get the driving term:

$$\varphi_2(t) = -a\left[2(n-1)I_1^0(t)\left(f_0+\gamma\left(\Delta_0+\frac{1}{n}\sum_j\Delta_j\right)\right)+\frac{2}{n}\gamma\sum_j\sum_k\Delta_k I_1^k(t)+\gamma^2 I_0^j(t)I_0^k(t)\right]\boldsymbol{\xi}_0 + a\sum_j\left[I_1^j(t)\left(2(-f_0-\gamma\Delta_0)\right)+\gamma^2\left(\sum_k I_0^j(t)I_0^k(t)+n^2\delta_{jk}\right)\right]\boldsymbol{\xi}_j$$

where $I_m^j(t)$ refers to the $j^{\text{th}}$ component of $I_m(t)$ in the $\{\boldsymbol{\xi}_j\}$ basis. This term finally contains the mixing effects of integration by demotivating neurons, evident in the entry of terms relating to $I_0^k(t)$ in the $\boldsymbol{\xi}_j$ component. Now we can enter this term into the integral to get

$$\boldsymbol{I}_2(t) = ae^{-Zt}\left(-\int_0^t e^{\frac{s}{\tau}}\left[2(n-1)I_1^0(s)\left(f_0+\gamma\left(\Delta_0+\frac{1}{n}\sum_j\Delta_j\right)\right)+\frac{2}{n}\gamma\sum_j\sum_k\Delta_k I_1^k(s)+\gamma^2 I_0^j(s)I_0^k(s)\right]ds\boldsymbol{\xi}_0 + \int_0^t e^{\frac{(1-\beta w)}{\tau}s}\sum_j\left[-2I_1^j(s)(f_0+\gamma\Delta_0)+\gamma^2\left(\sum_k I_0^j(s)I_0^k(s)+n^2\delta_{jk}\right)\right]ds\boldsymbol{\xi}_j\right)$$

Now that we have reached the lowest order mixing term, we will focus on the individual drive components $\boldsymbol{\xi}_j$. Recall that $I_0^j(t) = \frac{\beta w}{n}\frac{1}{1-\beta w}\left(1 - e^{-\frac{(1-\beta w)}{\tau}t}\right)\Delta_j$ so that $I_0^j(t)I_0^k(t) = \left(\frac{\beta w}{n(1-\beta w)}\right)^2\left(1 - 2e^{-\frac{(1-\beta w)}{\tau}t} + e^{-2\frac{1-\beta w}{\tau}t}\right)\Delta_j\Delta_k \geq 0$, except for any component involving interaction with $\boldsymbol{\xi}_1$, which will introduce an additional $\Delta_j\Delta_1(1 + w\chi_n)e^{-\frac{1-\beta w}{\tau}t}$ term. Whether a drive is stabilized or destabilized by the interactions between drives is dependent on whether this synergistic component can overcome the $-2I_1^j(s)(f_0 + \gamma\Delta_0)$ term, which is itself suppressive of the corresponding behaviour and in the dominant drive lacks the transient $-\frac{\tau}{1-\beta w}te^{-\frac{1-\beta w}{\tau}t}$. Then if $\frac{\tau}{1-\beta w}\Delta_j > \Delta_1(1 + w\chi_n)$, i.e. $\Delta_j$ is comparable in magnitude to or greater than $\Delta_1$, the drive will act in a switch-promoting capacity to destabilize Drive 1. Otherwise, the greater values of $\Delta_1\Delta_k$ in the synergistic term for Drive 1 will overpower the destabilizing influence of its competitors and the system will behave in a stability-promoting regime. The dependence on $\chi_n$ shows that this regime's size changes with the number of relevant drives, and once $n > 2 + \frac{1}{\beta w}$ switch-promoting is the only paradigm. The $\frac{1}{\beta w} = \frac{1}{\sigma'(0)f'(0)\tau w}$ term reflects the how nonlinear the system is; when the nonlinearities are sharp and the derivatives are high, the supralinearity of the system is dominant and the inhibitory neurons act to strongly amplify any competing drives. In our simulations, the parameters were such that $\frac{1}{\beta w} \approx 8$, and we see in **Extended Data Figure 8** that the stabilizing regime was much smaller and much less effective at suppressing challengers when $n = 10$. Reducing the steepness of the nonlinearities sustained the stabilizing regime even with larger numbers of competing drives.

*Other nonlinear formulations:*

In the following sections, we explore a number of other nonlinear frameworks that incorporate stochasticity. These analyses are purely numerical.

*Probabilistic rate network:*

We implemented a network of 200 neurons per drive with probabilistic inputs at every time step for a total of $2n$ neurons: $n$ motivating and $n$ demotivating. The firing rate vector $\boldsymbol{V}$ obeyed the stochastic differential equation

$$\frac{d\boldsymbol{V}}{dt} = W\sigma(\boldsymbol{V}) + \sigma(\boldsymbol{V}) \circ A\sigma(\boldsymbol{V}) - T\boldsymbol{V} + \mathbf{D}$$

where bolded capitalized symbols represent vectors in $\mathbb{R}^{2n}$ and non-bold capitalized symbols represent matrices in $\mathbb{R}^{2n\times 2n}$. The symbol $\circ$ represents the Hadamard product, i.e. $(\boldsymbol{A} \circ \boldsymbol{B})_i = A_iB_i$. $\boldsymbol{D} = \boldsymbol{\mu} + \epsilon\mathbf{N}$ where the elements of $\mathbf{N}$ at every time step are drawn from a Gaussian random variable with variance 1 and $\epsilon$ is the value in **Supplementary Table 1**. $\mu_j = 0$ for demotivating neurons. $\sigma(x) = \max(0, x)/n$ where $n$ is the total number of motivating neurons in the network. This acted to block input from neurons whose activity was below "threshold." The first half of the components of $\boldsymbol{V}$ correspond to the rates of the motivating neurons, while the second half correspond to the rates of the demotivating neurons. $W$ was defined as 4 block matrices:

$$W = \begin{pmatrix} 0 & D \to M \\ M \to D & 0 \end{pmatrix}$$

with

$$D \to M = \begin{pmatrix} -a \ldots & 0 \ldots & 0 \ldots & 0 \ldots \\ 0 \ldots & -a \ldots & 0 \ldots & 0 \ldots \\ 0 \ldots & 0 \ldots & -a \ldots & 0 \ldots \\ 0 \ldots & 0 \ldots & 0 \ldots & -a \ldots \end{pmatrix} \quad M \to D = \begin{pmatrix} -a \ldots & a \ldots & a \ldots & a \ldots \\ a \ldots & -a \ldots & a \ldots & a \ldots \\ a \ldots & a \ldots & -a \ldots & a \ldots \\ a \ldots & a \ldots & a \ldots & -a \ldots \end{pmatrix}$$

with every element $a$ sampled from an exponential distribution of mean $1$. This corresponds to every demotivating neuron inhibiting only the 100 motivating neurons corresponding to its own specific drive, while being stimulated by every motivating neuron *not* corresponding to its own drive. Motivating neurons protect their demotivating neurons. The supralinearity $A$ took a similar block form:

$$A = \begin{pmatrix} 0 & 0 \\ syn & 0 \end{pmatrix}$$

with the block matrix

$$syn = \begin{pmatrix} 0 \ldots & a \ldots & a \ldots & a \ldots \\ a \ldots & 0 \ldots & a \ldots & a \ldots \\ a \ldots & a \ldots & 0 \ldots & a \ldots \\ a \ldots & a \ldots & a \ldots & 0 \ldots \end{pmatrix}$$

This matrix produced supralinear integration of all other drives at the demotivating neurons alone. The time-constant matrix $T$ was defined as

$$T_{ij} = \begin{cases} 1 \text{ for the first half of components (motivating neurons, } i \leq n) \\ \dfrac{1}{1.0 + \log(1 + e^{V_k})} \text{ with } k = i - n \text{ (demotivating neurons, } i > n) \end{cases}$$

for $i = j$ and $0$ otherwise. This let the time constant of demotivating neurons grow as their corresponding motivating neurons increased in strength.

The system was run for 1000 time steps, with inputs to Drives 2 and 3 increasing by the value described in **Table 1** during steps 500 – 600. The plots in **Extended Data Figure 8b** show the outcomes of 100 separate runs of the system. The plots in **Extended Data Figure 8c** represent an individual run in each of the stabilizing, switching, and no-tertiary-drive conditions. "Drive output" was generated by taking the softmax of the mean of each motivating neuron population. The system was considered to have "switched" behaviours when the drive output value of Drive 1 was no longer the greatest value and never succeeded the now-dominant drive thereafter (i.e. $t_{switch} = \sup_t(D_1(t) > D_i \; \forall \; i \neq 1)$).

We observed that, as the number of drives in the system increased, we needed to lower the relative noise of each drive's input in order to see a strong stabilizing regime, though there generally was a stabilizing regime to some extent. This emerges from the perturbative analysis, but was exacerbated in the presence of greater noise. We speculate that the random noise onto the demotivating neurons ensured that they were always operating near the destabilizing regime even in the absence of manipulation of the tertiary drive.

*Spiking neurons:*

We implemented a network of $2n$ spiking neurons connected as in the rate network, with $W$ and $A$ as before, except with $W$ multiplied by a factor of $W^*$ and $A$ by a factor of $A^*$ (see **Supplementary Table 1**). The vector of neuronal voltages $\boldsymbol{V}$ obeys the differential equation

$$\frac{d\boldsymbol{V}}{dt} = W\boldsymbol{r} + \boldsymbol{s} \circ A\boldsymbol{s} + \boldsymbol{D} - T\boldsymbol{V}$$

with

$$T_{ij} = \begin{cases} \tau \text{ for the first half of components (motivating neurons, } i \leq n) \\ \dfrac{\beta\tau}{1.0 + \dfrac{\log(1+e^{3V_k})}{3}} \text{ with } k = i - n \text{ (demotivating neurons, } i > n) \end{cases}$$

when $i = j$ and 0 otherwise.

A neuron $i$ is said to spike when $V_i > \theta$, after which $V_i$ is set to $v_r$. $\boldsymbol{r}$ and $\boldsymbol{s}$ correspond to "postsynaptic currents" after a neuron fires a spike, obeying the differential equations:

$$\frac{d\boldsymbol{r}}{dt} = -\frac{\boldsymbol{r}}{\tau_r} + \sum_{t_{spike}} \boldsymbol{\delta}\,(t - t_{spike}) \quad \frac{d\boldsymbol{s}}{dt} = -T\boldsymbol{s} + \sum_{t_{spike}} \boldsymbol{\delta}\,(t - t_{spike})$$

with each component $\delta_i$ of $\boldsymbol{\delta}$ a Dirac delta function whenever neuron $i$ fires a spike. This gives a different decay rate for the nonlinear effects than the linear ones. The input vector $\boldsymbol{D}(t) = \boldsymbol{\mu}(t) + \epsilon\mathbf{N}$ as before, except the noise is applied to the input current of all neurons.

Spiking neural networks can be notoriously tricky to control. We found that our results did not hinge critically on particular parameter values, but that either the stabilizing or switching condition could be more difficult to elicit depending on the parameters. Networks with a greater $W^*$ tended to stabilize the dominant drive, while networks with a greater $A^*$ generally operated in the switching regime. The values listed in **Supplementary Table 1** seemed to elicit both behaviours without a great deal of hand-tuning the input levels, and so we use these to demonstrate both phenomena in the Figures.

**Temporal integration:**

*Estimation of parameters and their variance:* The instantaneous probability of termination, given that a fly has not yet terminated the mating by time $t$, was modeled as

$$p(t_{term} = t \mid \neg\, t_{term} < t) = p_0\tau(1 - e^{-\frac{t}{\tau}})$$

with $p_0$ the intensity of the stimulus and $\tau$ the time constant of integration, two model parameters we wish to fit. Each moment in time gives an approximate estimate of this value, which can be pooled together using the cumulative distribution $\sigma(t) = p(t_{term} \leq t)$, which can be derived by noting that

$$\frac{d\sigma}{dt} = p(t_{term} = t \mid \neg t_{term} < t)\big(1 - \sigma(t)\big) = p_0\tau(1 - e^{-\frac{t}{\tau}})(1 - \sigma(t))$$

this can trivially be solved to yield

$$-\log(1 - \sigma) = p_0\tau\left(t + \tau e^{-\frac{t}{\tau}}\right) + C$$

with $C$ the constant of integration. Because $\sigma(0) = 0$, we can immediately see that $C = -p_0\tau^2$ and solve to get

$$\log(1 - \sigma(t)) = p_0\tau\left(\tau(1 - e^{-\frac{t}{\tau}}) - t\right)$$

finally yielding

$$\sigma(t) = 1 - e^{-p_0\tau\left(t - \tau\left(1 - e^{-t/\tau}\right)\right)}.$$

The probability $p_{p_0,\tau}(x)$ of any particular observation $x$, given parameters, $p_0$ and $\tau$ is then

$$p_{p_0,\tau}(x) = \frac{d\sigma}{dt}\bigg|_{t=x} = p_0\tau\big(1 - e^{-x/\tau}\big)e^{-p_0\tau\left(x - \tau\left(1 - e^{-x/\tau}\right)\right)}$$

We can now attempt to fit our data to this model to obtain estimates of the parameters $p_0$ and $\tau$. One form of estimate is the maximum likelihood estimate, the values of the parameters "least surprised" by the data (those which predict the greatest likelihood of generating the data set). The likelihood $\mathcal{L}_{\{x\}}(p_0, \tau)$ of any particular data set $\{x\}$, assuming all samples are independent, is

$$\mathcal{L}_{\{x\}}(p_0, \tau) = \prod_x p_{p_0,\tau}(x)$$

and we choose as our estimates $\widehat{p_0}, \hat{\tau} = \operatorname{argmax}_{p_0,\tau} \mathcal{L}_{\{x\}}(p_0, \tau)$.

Any values $\widehat{p_0}$ and $\hat{\tau}$ that maximize $\mathcal{L}_{\{x\}}(p_0, \tau)$ also maximize $\log \mathcal{L}_{\{x\}}(p_0, \tau)$, or

$$\log \mathcal{L}_{\{x\}}(p_0, \tau) = \sum_x \log\big(p_{p_0,\tau}(x)\big)$$

However, our data is subject to an additional constraint: the stimulation is terminated at some upper bound value $u$. Thus, rather than consisting of termination times $\{x\}$, the data takes the form $\{x\} \cup \{\emptyset\}^{n_{per}}$ with $n_{per}$ the number of data points that persevere through the threat. The probability of observing $\emptyset$ is the probability that the corresponding sample $x$ would be greater than $u$, or $1 - \sigma(u)$. Then we can then account for this constraint in the likelihood function by re-writing $\mathcal{L}_{\{x\}}(p_0, \tau)$ as

$$\mathcal{L}_{\{x\}}(p_0, \tau) = \left(\prod_{x\in\{x\}} p_{p_0,\tau}(x)\right)\big(1 - \sigma(u)\big)^{n_{per}}$$

and thus

$$\log \mathcal{L}_{\{x\}}(p_0, \tau) = \sum_x \log\big(p_{p_0,\tau}(x)\big) + n_{per}\log\big(1 - \sigma(u)\big)$$

We can then maximize this expression to obtain our estimates $\widehat{p_0}$ and $\hat{\tau}$.

To do so, we take the gradient of $\log \mathcal{L}_{\{x\}}(p_0, \tau)$ and find where it equals 0.

$$\frac{\partial \log \mathcal{L}_{\{x\}}(p_0, \tau)}{\partial p_0} = \sum_{x\in\{x\}} \frac{\partial \log\big(p_{p_0,\tau}(x)\big)}{\partial p_0} + \frac{n_{per}\partial \log\big(1 - \sigma(u)\big)}{\partial p_0}$$

$$= \left[\sum_{x\in\{x\}} \frac{\partial}{\partial p_0}\left(\log\left(p_0\tau\big(1 - e^{-x/\tau}\big)e^{-p_0\tau\left(x-\tau\left(1-e^{-x/\tau}\right)\right)}\right)\right)\right] - n_{per}\tau\left(u - \tau\left(1 - e^{-\frac{u}{\tau}}\right)\right)$$

$$= \left[\sum_{x\in\{x\}} \frac{\partial}{\partial p_0}\left(\log(p_0\tau) + \log\big(1 - e^{-x/\tau}\big) - p_0\tau\left(x - \tau\big(1 - e^{-x/\tau}\big)\right)\right)\right] - n_{per}\tau\left(u - \tau\left(1 - e^{-\frac{u}{\tau}}\right)\right)$$

$$\frac{\partial \log \mathcal{L}_{\{x\}}(p_0, \tau)}{\partial p_0} = \left[\sum_{x\in\{x\}} \frac{1}{p_0} - \tau\left(x - \tau\left(1 - e^{-\frac{x}{\tau}}\right)\right)\right] - n_{per}\tau\left(u - \tau\left(1 - e^{-\frac{u}{\tau}}\right)\right)$$

Similarly,

$$\frac{\partial \log \mathcal{L}_{\{x\}}(p_0, \tau)}{\partial \tau} = \sum_{x\in\{x\}} \frac{\partial \log\big(p_{p_0,\tau}(x)\big)}{\partial \tau} + \frac{n_{per}\partial \log\big(1 - \sigma(u)\big)}{\partial \tau}$$

Splitting into two pieces:

$$\frac{n_{per}\partial \log\big(1 - \sigma(u)\big)}{\partial \tau} = n_{per}p_0\left(2\tau\big(1 - e^{-u/\tau}\big) - u\big(1 + e^{-u/\tau}\big)\right)$$

and

$$\sum_{x\in\{x\}} \frac{\partial \log\left(p_{p_0,\tau}(x)\right)}{\partial \tau} = \sum_{x\in\{x\}} \frac{\partial}{\partial \tau}\left(\log(p_0\tau) + \log\left(1 - e^{-\frac{x}{\tau}}\right) - p_0\tau\left(x - \tau\left(1 - e^{-\frac{x}{\tau}}\right)\right)\right)$$

$$= \sum_{x\in\{x\}} \left( \frac{1}{\tau} - \frac{x e^{-x/\tau}}{\tau^2(1+e^{-x/\tau})} - p_0 \left(2\tau\left(e^{-\frac{x}{\tau}}-1\right) + x\left(1+e^{-x/\tau}\right)\right)\right)$$

$$= \sum_{x\in\{x\}} \left( \frac{\tau\left(1+p_0\tau(2\tau - x)\right) - (x+\tau+4p_0\tau^3)e^{-\frac{x}{\tau}} + p_0\tau^2 e^{-\frac{2x}{\tau}}(x+2\tau)}{\tau^2(1-e^{-x/\tau})}\right)$$

At the maximum likelihood values $\widehat{p_0}$ and $\hat{\tau}$, both of these are 0, so

$$0 = \left[\sum_{x\in\{x\}} \frac{1}{\widehat{p_0}} - \hat{\tau}\left(x - \hat{\tau}\left(1 - e^{-\frac{x}{\hat{\tau}}}\right)\right)\right] - n_{per}\hat{\tau}\left(u - \hat{\tau}\left(1 - e^{-\frac{u}{\hat{\tau}}}\right)\right)$$

or

$$\frac{n_{term}}{\widehat{p_0}} = \left[\sum_{x\in\{x\}} \hat{\tau}\left(x - \hat{\tau}\left(1 - e^{-\frac{x}{\hat{\tau}}}\right)\right)\right] + n_{per}\hat{\tau}\left(u - \hat{\tau}\left(1 - e^{-\frac{u}{\hat{\tau}}}\right)\right)$$

where $n_{per}$ is the number of flies which terminate the mating in response to the stimulus. If $n$ is the total number of flies in the experiment, so that $\frac{n_{term}}{n} = p_{term}$ and $\frac{n_{per}}{n} = 1 - p_{term}$ then we have

$$\widehat{p_0} = \frac{p_{term}/\hat{\tau}}{(1-p_{term})\left(u - \hat{\tau}\left(1 - e^{-\frac{u}{\hat{\tau}}}\right)\right) + \frac{\sum_{x\in\{x\}}\left(x - \hat{\tau}\left(1 - e^{-\frac{x}{\hat{\tau}}}\right)\right)}{n}}$$

which is the exact maximum likelihood estimate for $p_0$ given $\tau$.

Similarly, we have (after dividing by $n$) that

$$0 = (1-p_{term})\widehat{p_0}\left(2\hat{\tau}\left(1 - e^{-\frac{u}{\hat{\tau}}}\right) - u\left(1 + e^{-\frac{u}{\hat{\tau}}}\right)\right)$$
$$+ \frac{1}{n}\sum_{x\in\{x\}} \left( \frac{1}{\hat{\tau}} - \frac{x e^{-x/\hat{\tau}}}{\hat{\tau}^2(1+e^{-x/\hat{\tau}})} - p_0 \left(2\hat{\tau}\left(e^{-\frac{x}{\hat{\tau}}}-1\right) + x\left(1+e^{-x/\hat{\tau}}\right)\right)\right)$$

We were unable to find an analytical solution to this equation, and so to estimate $\hat{\tau}$ we performed grid search to maximize the log-likelihood, for each value using the analytical value for $\widehat{p_0}$ given the current estimate for $\tau$. We used a step size of 0.001, and searched parameters ranging from 0.01 to 40.

Likewise, for the model in which there is no $\tau$, the maximum likelihood estimate for $p_{0,null}$ is

$$\widehat{p_{0,null}} = \frac{p_{term}}{(1-p_{term})u + \frac{1}{n}\sum_{x\in\{x\}} x}$$

To estimate the variance of the parameter fits, we use the Fisher Information and Cramér-Rao bound, which says

$$Cov(\boldsymbol{\theta}) \geq \mathcal{I}(\boldsymbol{\theta})^{-1}$$

for any consistent estimator of a parameter vector $\boldsymbol{\theta}$ where $\mathcal{I}(\theta)$ is the Fisher Information matrix defined as

$$\mathcal{I}(\theta)_{ij} = -\mathbb{E}\left[\left(\frac{\partial^2 \log p(\{x\}|\boldsymbol{\theta})}{\partial\theta_i\partial\theta_j}\right)\right]$$

For our model,

$$\mathcal{I}\begin{pmatrix}\hat{\tau}\\ \widehat{p_0}\end{pmatrix} = -\mathbb{E}\left[\begin{pmatrix}\frac{\partial^2 \log p(\{x\}|\hat{\tau},\widehat{p_0})}{\partial\tau^2} & \frac{\partial^2 \log p(\{x\}|\hat{\tau},\widehat{p_0})}{\partial\tau\partial p_0}\\ \frac{\partial^2 \log p(\{x\}|\hat{\tau},\widehat{p_0})}{\partial\tau\partial p_0} & \frac{\partial^2 \log p(\{x\}|\hat{\tau},\widehat{p_0})}{\partial p_0^2}\end{pmatrix}\right]$$

The bottom right element is simply

$$\frac{\partial^2 \log p(\{x\}|\hat{\tau},\widehat{p_0})}{\partial\tau^2} = -\frac{n_{term}}{\widehat{p_0}^2}$$

so that the expectation is $\mathbb{E}\left[\frac{\partial^2 \log p(\{x\}|\hat{\tau},\widehat{p_0})}{\partial\tau^2}\right] = -\frac{n\sigma(u)}{\widehat{p_0}^2}$

The diagonal terms are

$$\frac{\partial^2 \log p(\{x\}|\hat{\tau},\widehat{p_0})}{\partial\tau\partial p_0} = n_{per}\left(2\hat{\tau}\left(1-e^{-u/\hat{\tau}}\right) - u\left(1+e^{-\frac{u}{\hat{\tau}}}\right)\right) + \sum_{x\in\{x\}} 2\hat{\tau}\left(1-e^{-\frac{x}{\hat{\tau}}}\right) - x\left(1+e^{-\frac{x}{\hat{\tau}}}\right)$$

The expectation of the terms on the left (those resulting from the truncation) is

$$n(1-\sigma(u))\left(2\hat{\tau}\left(1-e^{-u/\hat{\tau}}\right) - u\left(1+e^{-\frac{u}{\hat{\tau}}}\right)\right)$$

The top left term of the Fisher Information is a little messy. The contribution from the values above the cutoff $u$ is

$$n(1-\sigma(u))p_0\left(\frac{(u^2+2u\tau)e^{-\frac{u}{\tau}}}{\tau^2} - 2\left(1-e^{-u/\tau}\right)\right)$$

while the contribution from the values within the range of the assay (without taking the expectation) are

$$\sum_{x\in\{x\}} \frac{e^{-x/\tau}}{\tau^4\left(e^{\frac{x}{\tau}}-1\right)^2}\Big[-p_0\tau^2(x^2+2x\tau+2\tau^2) + e^{\frac{3x}{\tau}}\tau^2(2p_0\tau-1)$$
$$+ e^{\frac{x}{\tau}}\tau(-2x-\tau+2p_0x^2\tau+4p_0x\tau^2+6p_0\tau^3)$$
$$- e^{2x/\tau}\left(-2\tau^2+6p_0\tau^4+2x\tau(p_0\tau-1)+x^2(1+p_0\tau^2)\right)\Big]$$

For those elements of the Fisher Information for which we did not find a closed form solution of the expectation, we used the value at the maximum likelihood estimate (essentially assuming that the probability density of those values is peaked near our estimate, **Extended Data Figure 3c**). We then computed the Fisher Information and took the inverse to find the Cramér-Rao bound, which we used as our estimate of the covariance matrix. The plotted variances of the parameters correspond to the diagonal of this matrix.

The standard error of $p_{0,null}$, likewise, is

$$SE(p_{0,null}) = \left(\frac{p_{0,null}}{\sqrt{n}}\right)\left(\frac{1}{\sqrt{(1-e^{-p_{0,null}u}) - \left(up_{0,null}e^{-up_{0,null}}\right)^2}}\right)$$

Because the maximum likelihood estimate is asymptotically normally distributed with variance given by the Cramér-Rao bound, for statistical comparisons of model fits we performed a Welch's t-test for unequal variances and unequal sample sizes.

*Termination probability of a stimulus and its relation to the model parameters:*

The above analysis is of a stimulation that begins at time $t = 0$ and is sustained forever. **Figure 2b** focuses on stimuli that last a finite duration. To analyze the case of a short stimulus, especially one much shorter than $\tau$, we replaced the expression for $p(t_{term} = t \mid \neg\, t_{term} < t)$ with $p(t_{term} = t \mid \neg\, t_{term} < t) = f(t)$ where

$$f(t) = \begin{cases} p_0\tau\left(1 - e^{-\frac{t}{\tau}}\right) & t < \mu \\ p_0\tau\left(1 - e^{-\frac{\mu}{\tau}}\right)e^{\frac{-(t-\mu)}{\tau}} & t \geq \mu \end{cases}$$

with $\mu$ the duration of the stimulus. The peak of this function is $f(\mu) = p_0\tau\left(1 - e^{-\frac{\mu}{\tau}}\right)$. Then to plot the change in the peak of this function as $\mu$ and $\tau$ are adjusted, we plotted $\partial_\tau f(\mu)$ and $\partial_\mu f(\mu)$:

$$\partial_\mu f(\mu) = \ p_0 e^{-\mu/\tau} \quad \partial_\tau f(\mu) = p_0\left(1 - \ \left(\frac{\mu}{\tau} - 1\right)\ e^{-\mu/\tau}\right)$$

For the plot, we divided both equations by the maximal value of each, which is $p_0$ for $\partial_\mu f(\mu)$ and $p_0(1 - e^{-2})$ for $\partial_\tau f(\mu)$. The integral of $f(t)$, as below, is $p_0\tau\mu$, which was used to plot the "differential change in integrated output."

To analyze how this translates to the termination probability, we have to again solve the differential equation

$$\frac{d\sigma}{dt} = p(t_{term} = t \mid \neg t_{term} < t)\big(1 - \sigma(t)\big)$$

$$\frac{d\sigma}{dt} = f(t)\big(1 - \sigma(t)\big)$$

whose solution is

$$-\log\big(1 - \sigma(t)\big) = \int_0^t f(t')dt' + C$$

or

$$\sigma(t) = \ 1 - e^{-\int_0^t f(t')dt'}$$

as before. We wish to evaluate $\Phi = \lim_{t\to\infty} \sigma(t)$, which we can find from $\int_0^t f(t')dt'$.

$$\int_0^t f(t')dt' = \int_0^\mu f(t')dt' + \int_\mu^t f(t')dt'$$

$$= p_0\tau\left[\int_0^\mu \left(1 - e^{-\frac{t'}{\tau}}\right)dt' + \left(1 - e^{-\frac{\mu}{\tau}}\right)\int_\mu^t e^{\frac{-(t'-\mu)}{\tau}}dt'\right]$$

$$= p_0\tau\left[\mu - \tau\left(1 - e^{-\frac{\mu}{\tau}}\right) + \tau\left(1 - e^{-\frac{\mu}{\tau}}\right)\left(1 - e^{\frac{-(t-\mu)}{\tau}}\right)\right]$$

so then

$$\Phi = 1 - \lim_{t\to\infty} e^{-p_0\tau\left[\mu-\tau\left(1-e^{-\frac{\mu}{\tau}}\right)+\tau\left(1-e^{-\frac{\mu}{\tau}}\right)\left(1-e^{\frac{-(t-\mu)}{\tau}}\right)\right]}$$
$$= 1 - \lim_{t\to\infty} e^{-p_0\tau\left[\mu-\tau\left(e^{\frac{\mu}{\tau}}-1\right)\left(e^{\frac{-t}{\tau}}\right)\right]} = 1 - e^{-p_0\tau\mu}$$

If $p_0\tau\mu \ll 1$ (as is the case for our short pulses, where $\mu \approx 1$ sec, $p_0\tau \ll 1$ sec$^{-1}$) then

$$\Phi \approx p_0\tau\mu$$

so if $\mu \ll \tau$, then $\frac{d\Phi}{d\tau} \ll \frac{d\Phi}{d\mu}$ and the termination probability grows much more quickly with changes in $\mu$ than changes in $\tau$.

However, when $\mu \gg \tau$, as in our sustained stimuli and some grooming stimuli, then $\frac{d\Phi}{d\tau} \gg \frac{d\Phi}{d\mu}$ and the termination probability grows much more quickly with changes in $\tau$ than changes in $\mu$.

*Robustness of the fitting algorithm:*

The cumulative distribution consists of 60 data points (albeit with correlational structure), while the model has only two parameters, making it relatively robust to overfitting. We evaluate the ability of our statistical model to capture temporal integration in **Extended Data Figure 5**. In this figure, we generate synthetic data sets following the distribution of the model as described above (by sampling from a uniform distribution between 0 and 1 and then inverting the cumulative distribution function), varying the model parameters, and then applied our maximum likelihood estimate. The inverse cumulative distribution is

$$\sigma^{-1}(z) = \tau\left(1 + W\left(\frac{(1-z)^{\frac{1}{p_0\tau^2}}}{e}\right)\right) - \frac{\log(1-z)}{p_0\tau}$$

where $W(x)$ is the Lambert W-function, so that $W(xe^x) = x$, evaluated on the main branch (which is well-defined, since $0 \le (1-z)^{\frac{1}{p_0\tau^2}} \le 1$).

*Measuring augmentation:*

The data was assumed to follow a model in which the termination probability $p_2(t)$ of a second pulse $t$ units of time after an initial pulse (with termination probability $p_1$) is potentiated by a factor $a_t$.,so that $p_2(t) = p_1 a_t$. If two pulses are applied in succession, then the probability of terminating the mating $p_{term}$ follows the relation

$$p_{term}(t) = p_1 + p_2(t) - p_1 p_2(t)$$

which may be rewritten as

$$p_{term}(t) = p_1\big(1 + a_t(1 - p_1)\big)$$

To infer $a_t$, we performed two sets of experiments – one in which a single pulse was provided, to estimate $p_1$, and one in which two pulses were provided, to use our estimate of $p_1$ to estimate $a_t$. The likelihood was modeled as arising from the following probabilistic model:

$$x_1 \sim Binomial(n_1, p_1)$$
$$x_2 \sim Binomial\left(n_2, p_1\big(1 + a_t(1 - p_1)\big)\right)$$

with $n_1$ and $n_2$ fixed and corresponding to the sample size of the single pulse and paired pulse experiments respectively. We then estimated the posterior probability distribution of $a_t$ by using Hamiltonian Monte Carlo sampling with a log-normal prior on $a_t$ so that

$$\log a_t \sim \mathcal{N}(0,1)$$

and the Jeffrey's prior on $p_1$ as above. The analysis code was written using the TensorFlow-Probability package `edward2`. The data presented in the graphs correspond to the 16%-84% credible interval and the mean posterior value.

**Table 1: Modeling parameters**

| Figure | Parameter | Value |
|---|---|---|
| **Figure 5b** | | |
| Top row | $n$ | 3 |
| | $f(x)$ | $1.5\log(1 + e^{x/2})$ |
| | $\sigma(x)$ | $1.5\tan^{-1}(x)$ |
| | $w_{ij}$ | 2 for $i = j$, 1 otherwise |
| | $\alpha_{ijk}$ | 2, except in independent, where all are 0 |
| | $\delta_i(t)$ | $\delta_1(t) = .65$,<br>$\delta_2(t) = 0.1 + 0.65[\mu(t - 20)]$<br>$\delta_3(t) = 0.1 + .5[\mu(t - 20)]$ |
| | $\tau_i$ | All 2 |
| | Initial conditions | $D_i(0) = I_i(0) = 0$ |
| Bottom row | $n$ | 3 |
| | $f(x)$ | $1.5\log(1 + e^{x/2})$ |
| | $\sigma(x)$ | $1.5\tan^{-1}(x)$ |
| | $w_{ij}$ | -2 for $i = j$, 1 otherwise |
| | $\alpha_{ijk}$ | 2, except in independent, where all are 0 |
| | $\delta_i(t)$ | $\delta_1(t) = 1.5$,<br>$\delta_2(t) = 0.1 + 2.45[\mu(t - 20)]$<br>$\delta_3(t) = 0.1 + 2[\mu(t - 20) - \mu(t - 40)]$ |
| | $\tau_i$ | All 2 |
| | Initial conditions | $D_i(0) = I_i(0) = 0$ |
| **Figure 5c** | | |
| Weakly dominant drive | $\delta_i(t)$ | $\delta_1(t) = .65$,<br>$\delta_2(t) = 0.1 + a[\mu(t - 20)]$<br>$\delta_3(t) = 0.1 + b[\mu(t - 20)]$<br>(a and b are stepped from 0.3 to 1.0 in increments of 0.01) |
| Strongly dominant drive | $\delta_i(t)$ | $\delta_1(t) = 1.5$,<br>$\delta_2(t) = 0.1 + a[\mu(t - 20)]$<br>$\delta_3(t) = 0.1 + b[\mu(t - 20)]$<br>(a and b are stepped from 0.0 to 3.0 in increments of 0.05) |
| **Extended Data Figure 7** | | |
| 3 drives | Neurons per drive | 100 motivating, 100 demotivating |
| | Mean drive 1 input | 1.1 |
| | Baseline drive level | 0.0 |
| | Mean drive 2 input (during increase) | 1.15 |
| | Mean drive 3 input (during increase) | None: 0.0 |

| | | |
|---|---|---|
| | | Stabilizing: 0.25 |
| | | Switching: 1.13 |
| | Input noise | 0.25 |
| 10 drives | Neurons per drive | 100 motivating, 100 demotivating |
| | Mean drive 1 input | 1.9 |
| | Baseline drive level | 0.0 |
| | Mean drive 2 input (during increase) | 1.95 |
| | Mean drive 3 input (during increase) | None: 0.0 |
| | | Stabilizing: 0.25 |
| | | Switching: 1.93 |
| | All other drives: | 0.0 |
| | Input noise | 0.1 |
| **Extended Data Figure 8** | | |
| | Mean drive 1 input | 1.9 |
| | Baseline drive 2 and 3 input | 1.0 |
| | Drive 2 during increase | 2.05 |
| | Drive 3 during increase | None: 0.0 |
| | | Stabilizing: 1.85 |
| | | Switching: 1.95 |
| | $\tau_r$ | 2.0 |
| | $\theta$ | 10.0 |
| | $v_r$ | -1.0 |
| | $\beta$ | 2.0 |
| | Noise level | 1.25 |
| | $W^*$ | 8.0 |
| | $A^*$ | $\sqrt{10}$ |

(Note: $\mu(x)$ is the Heaviside step function, equal to 0 when $x < 0$ and 1 when $x \geq 0$).

**Table 2: Genotypes and number of flies per experiment**

| Figure | Label | Genotype | Condition | N |
|---|---|---|---|---|
| **Figure 1** | | | | |
| a) | DIN>Chr-tdTomato | w-;NP2719-Gal4, repo-Gal80/+;UAS-CsChrimson-tdTomato/+ | | |
| | DIN>MCFO | pBPhsFlp2::PEST; NP2719-Gal4, repo-Gal80/+; pJFRC210-10XUAS-FRT>STOP>FRT-myr::smGFP-OLLAS, pJFRC201-10XUAS-FRT>STOP>FRT-myr::smGFP-HA, pJFRC240-10XUASFRT>STOP>FRT-myr::smGFP-V5-THS-10XUAS-FRT>STOP>FRT-myr::smGFP-FLAG/+ (MCFO-2 from Nern *et al.*, heat shocked twice for 15 minutes each at 37°C as an adult fly, separated by one day, and dissected at least three days later to allow for expression of the tags) | | |
| | DIN>SytGFP,Denmark | w-;NP2719-Gal4,repo-Gal80/+;UAS-SytGFP,UAS-Denmark/+ | | |
| b) | DIN>ACR1 | w-;NP2719-Gal4/+;UAS-GtACR1-eYFP/+ | No light | 8 |
| | | | Light | 9 |
| | | | No light, no retinal | 6 |
| | | | Light, no retinal | 9 |
| | DIN>GFP | w-;NP2719-Gal4/+;UAS-GFP/+ | No light | 8 |
| | | | Light | 9 |
| c) | All flies | w-;NP2719-Gal4+;UAS-GtACR1-eYFP/+ | No light | 19 |
| | | | Off at 5 | 11 |
| | | | Off at 20 | 11 |
| | | | Off at 30 | 18 |

| | | | | |
|---|---|---|---|---|
| | | | On at 20 | |
| d) | DIN>ACR1 | w-;NP2719-Gal4+;UAS-GtACR1-eYFP/+ | No light | 17 |
| | | | Light | 15 |
| | DIN>GFP | w-;NP2719-Gal4/+;UAS-GFP/+ | No light | 8 |
| | | | Light | 6 |
| | +/ACR1 | w-;+/+;UAS-GtACR1-eYFP/+ | No light | 14 |
| | | | Light | 12 |
| e) | DIN>ACR1 | w-;NP2719-Gal4/+;UAS-GtACR1-eYFP/+ | Light + heat threat | 14 |
| | | | No light | 11 |
| | | | No heat | 10 |
| | | | Tonic | 14 |
| f) | DIN>Chr | w-;NP2719-Gal4/+; UAS-CsChrimson-tdTomato | No light | 89 |
| | No retinal | | 1 minute | 11 |
| | | | 5 minutes | 14 |
| | | | 10 minutes | 16 |
| | | | 15 minutes | 22 |
| | | | 20 minutes | 11 |
| | 2 sec light pulse | | 1 minute | 35 |
| | | | 5 minutes | 34 |
| | | | 10 minutes | 20 |
| | | | 15 minutes | 37 |
| | | | 20 minutes | 14 |
| | 10 minutes into mating | | | 35 |
| | 15 minutes into mating | | | 40 |
| **Figure 2** | | | | |
| a) | DIN>Chr | w-; NP2719-Gal4/+; UAS-CsChrimson-tdTomato/+ | 10 min, 500 ms | 51 |
| | | | 10 min, 1 sec | 39 |
| | | | 15 min, 500 sec | 47 |
| | | | 15 min, 1 sec | 46 |
| c-e) | DIN>Chr | w-;NP2719-Gal4/+;UAS-CsChrimson-tdTomato/+ | 10 min, low intensity | 107 |
| | | | 10 min, medium intensity | 116 |
| | | | 10 min, high intensity | 87 |
| | | | 15 min, low intensity | 126 |
| | | | 15 min, medium | 110 |

| | | | | |
|---|---|---|---|---|
| | | | intensity | |
| | | | 15 min, high intensity | 85 |
| **Figure 3** | | | | |
| a) | DIN>Tnt, Crz>Chr | w-; NP2719-Gal4/UAS-TntG; Crz-LexA/LexAop-CsChrimson-tdTomato | No light, 5 min | 10 |
| | | | No light, 10 min | 8 |
| | | | Light, 5 min | 14 |
| | | | Light, 10 min | 6 |
| | DIN>GFP, Crz>Chr | w-;NP2719-Gal4/UAS-CD8-GFP;Crz-LexA/LexAop2-Chr-tdTomato | Light / 5 minutes | 16 |
| | | | Light / 10 minutes | 12 |
| | | | No light / 5 minutes | 10 |
| | | | No light / 10 minutes | 18 |
| b) | DIN>Tnt, Crz>Chr | w-;NP2719-Gal4/UAS-tntG;Crz-LexA/LexAop2-Chr-tdTomato | Light | 10 |
| | | | No light | 9 |
| | DIN>GFP, Crz>Chr | w-;NP2719-Gal4/UAS-CD8-GFP;Crz-LexA/LexAop2-Chr-tdTomato | Light | 9 |
| | | | No light | 9 |
| c) | DIN>Chr, Crz>Kir2.1 | w-; NP2719-Gal4/LexAop-Kir2.1; Crz-LexA/UAS-CsChrimson-tdTomato | | 90 |
| | DIN>Chr, Crz>GFP | w-; NP2719-Gal4/LexAop-GFP; Crz-LexA/UAS-CsChrimson-tdTomato | | 91 |
| d) | DIN>Chr, TH>TrpA1 | w-; NP2719-Gal4/LexAop-TrpA1; TH-LexA/UAS-CsChrimson-tdTomato | Warmth | 108 |
| | | | No warmth | 119 |
| e,f,g) | DIN>Chr, jGCaMP7f | w-; NP2719-Gal4, Repo-Gal80/+; UAS-jGCaMP7f/UAS-CsChrimson-tdTomato | | 9 |
| **Figure 4** | | | | |
| a) | Groom>Chr | w-;R45G01-LexA/+;LexAop2-CsChrimson-tdTomato | 1 minute | 4 |
| | | | 5 minutes | 7 |

| | | | | |
|---|---|---|---|---|
| | | | 10 minutes | 20 |
| | | | 15 minutes | 8 |
| | Groom>Chr, DIN>Tnt | w-;R45G01-LexA,NP2719-Gal4/UAS-tntG;LexAop2-CsChrimson-tdTomato/+ | 1 minute | 7 |
| | | | 5 minutes | 4 |
| | | | 10 minutes | 7 |
| | | | 15 minutes | 7 |
| | Groom>Chr, DIN>GFP | w-;R45G01-LexA,NP2719-Gal4/UAS-CD8-GFP;LexAop2-CsChrimson-tdTomato/+ | 1 minute | 2 |
| | | | 5 minutes | 8 |
| | | | 10 minutes | 25 |
| | | | 15 minutes | 15 |
| b) | Groom>Chr | w-; R45G01-LexA/+ ; LexAop2-CsChrimson-tdTomato/+ | Heat | 26 |
| | | | Light | 30 |
| | | | Both | 45 |
| c) | 5 minutes | w-;NP2719-Gal4/+;UAS-Chr-tdTomato/+ | 41°C / Light | 31 |
| | | | 41°C / Heat | 16 |
| | | | 41°C / Both | 28 |
| | | | 37°C / Light | 38 |
| | | | 37°C / Heat | 12 |
| | | | 37°C / Both | 49 |
| | 10 minutes | w-;NP2719-Gal4/+;UAS-Chr-tdTomato/+ | Light | 41 |
| | | | Heat | 49 |
| | | | Both | 47 |
| | 15 minutes | w-;NP2719-Gal4/+;UAS-Chr-tdTomato/+ | Light | 35 |
| | | | Heat | 39 |
| | | | Both | 38 |
| d) | | w-; Split-Gal4-p65.AD/+;Split-Gal4-Gal4.DBD/UAS-CsChrimson-tdTomato | | 328 |
| e) | | w-; Split-Gal4-p65.AD/+;Split-Gal4-Gal4.DBD/UAS-CsChrimson-tdTomato | | 310 |
| f) | $DI_{Ag}$>Chr | w-;R27E07-p65.AD/+;R20F03-Gal4.DBD/UAS-CsChrimson-tdTomato | Heat | 37 |
| | | | Light | 35 |
| | | | Both | 43 |

| | | | | |
|---|---|---|---|---|
| g) | DIN>ACR1, Groom>Chr | w-; NP2719-Gal4, R45G01-LexA/UAS-GtACR1-eYFP; UAS-CsChrimson-tdTomato/+ | DINs silenced during grooming | 30 |
| | | | Heat + green, no grooming | 38 |
| | | | Heat + grooming | 37 |
| | | | Grooming alone | 36 |
| **Figure 5** | | | | |
| e) | SS02547>Chr | w-; R42B02-p65.AD/+;VT059427-Gal4.DBD/UAS-CsChrimson-tdTomato | 41°C heat threat | 32 |
| | | | 500 ms pulse, 10 min | 42 |
| | | | Both (500 ms pulse, 10 min) | 31 |
| | | | 10 sec pulse | 31 |
| | | | Both (10 sec pulse, 10 min) | 28 |
| | | | 37°C heat threat | 40 |
| | | | 500 ms pulse, 10 min | 45 |
| | | | Both (500 ms pulse, 10 min) | 39 |
| | | | 10 sec pulse | 39 |
| | | | Both (10 sec pulse, 10 min) | 39 |
| **Extended Data Figure 1** | | | | |
| a) | DIN>$Shi^{ts}$ | w-;NP2719-Gal4/NP2719-Gal4;UAS-Shibire-ts/UAS-Shibire-ts | 41°C | 22 |
| | | | 23°C | 9 |
| | DIN | w-;NP2719-Gal4/NP2719-Gal4;+/+ | 41°C | 20 |
| | | | 23°C | 12 |
| | $Shi^{ts}$ | w;+/+;UAS-Shibire-ts/UAS-Shibire-ts | 41°C | 12 |

| | | | | |
|---|---|---|---|---|
| | | | 23°C | 6 |
| b) | All flies | w-;NP2719-Gal4/NP2719-Gal4;UAS-Shibire-ts/UAS-Shibire-ts | No threat | 19 |
| | | | 1 min 41°C | 13 |
| | | | 30°C constant | 7 |
| **Extended Data Figure 2** | | | | |
| a) | | w-;NP2719-Gal4/+; UAS-CsChrimson-tdTomato/+ | Green | 36 |
| | | w-;NP2719-Gal4/+; UAS-CsChrimson-tdTomato/+ | No green | 38 |
| b) | | w-;NP2719-Gal4/+;UAS-CsChr-tdTomato/+ | No light | 89 |
| | | | No ret | 11 |
| | | | Courtship | 7 |
| | | | Tonic | 30 |
| | | | Mating | 9 |
| c) | | w-;NP2719-Gal4/+;UAS-CsChr-tdTomato/+ | | 30 |
| **Extended Data Figure 3** | | | | |
| b) | As in Figure 2c | | | |
| **Extended Data Figure 4** | | | | |
| a) | DIN>Chr | w-; NP2719-Gal4/+; UAS-CsChrimson-tdTomato/+ | 10 min, single pulse | 53 |
| | | | 15 min, single pulse | 56 |
| | | | 10 min, 0.5 s ISI | 70 |
| | | | 10 min, 1 s ISI | 72 |
| | | | 10 min, 5 s ISI | 79 |
| | | | 10 min, 10 s ISI | 88 |
| | | | 10 min, 20 s ISI | 67 |
| | | | 10 min, 40 s ISI | 69 |
| | | | 15 min, 0.5 s ISI | 65 |
| | | | 15 min, 1 s ISI | 64 |
| | | | 15 min, 5 s | 75 |

| | | | | |
|---|---|---|---|---|
| | | | ISI | |
| | | | 15 min, 10 s ISI | 71 |
| | | | 15 min, 20 s ISI | 67 |
| | | | 15 min, 40 s ISI | 81 |
| b) | Groom>Chr | w-;R45G01-LexA/+;LexAop2-CsChrimson-tdTomato | 3 sec, 5 min | 25 |
| | | | 3 sec, 10 min | 57 |
| | | | 3 sec, 15 min | 43 |
| | | | 6 sec, 5 min | 26 |
| | | | 6 sec, 10 min | 43 |
| | | | 6 sec, 15 min | 39 |
| d) | Groom>Chr | w-;R45G01-LexA/+;LexAop2-CsChrimson-tdTomato | Single pulse, 10 min | 57 |
| | | | 5 sec ISI, 10 min | 36 |
| | | | 10 sec ISI, 10 min | 41 |
| | | | 20 sec ISI, 10 min | 39 |
| | | | Single pulse, 15 min | 43 |
| | | | 5 sec ISI, 15 min | 42 |
| | | | 10 sec ISI, 15 min | 35 |
| | | | 20 sec ISI, 15 min | 31 |
| e) | $AG_{Desc}$>Chr | w-;R27E07-p65.AD/+;R20F03-Gal4.DBD/UAS-CsChrimson-tdTomato | 10 min, 700 ms pulse | 49 |
| | | | 1 sec ISI, 10 min | 53 |
| | | | 5 sec ISI, 10 min | 47 |
| | | | 10 sec ISI, 10 min | 41 |
| | | | 20 sec ISI, 10 min | 55 |
| | | | 15 min, 300 | 54 |

| | | | | |
|---|---|---|---|---|
| | | | ms pulse | |
| | | | 1 sec ISI, 15 min | 35 |
| | | | 5 sec ISI, 15 min | 37 |
| | | | 10 sec ISI, 15 min | 61 |
| | | | 20 sec ISI, 15 min | 47 |
| f) | Groom>Chr, DIN>ACR1 | w-;NP2719-Gal4, R45G01-LexA/UAS-GtACR1-eYFP;LexAop2-CsChrimson-tdTomato/+ | Red only, one pulse | 55 |
| | | | Red + green, one pulse | 34 |
| | | | Two red pulses, no green | 44 |
| | | | Two red pulses + green | 47 |
| | Groom>Chr, DIN>GFP | w-;NP2719-Gal4, R45G01-LexA/UAS-GFP;LexAop2-CsChrimson-tdTomato/+ | Red only, one pulse | 42 |
| | | | Red + green, one pulse | 39 |
| | | | Two red pulses, no green | 41 |
| | | | Two red pulses + green | 44 |
| g) | Groom>Chr, DIN>ACR1 | w-;NP2719-Gal4, R45G01-LexA/UAS-GtACR1-eYFP;LexAop2-CsChrimson-tdTomato/+ | Single pulse | 41 |
| | | | Green in between | 48 |
| | Groom>Chr, DIN>GFP | w-;NP2719-Gal4, R45G01-LexA/UAS-GFP;LexAop2-CsChrimson-tdTomato/+ | Single pulse | 41 |
| | | | Green in between | 51 |
| **Extended Data Figure 5** | | | | |

| | | | | |
|---|---|---|---|---|
| a) | Left and middle panels | w-; NP2719-Gal4, Repo-Gal80/LexAop-GFP; Crz-LexA/UAS-myr-tdTomato | | |
| | Right panel | w-; NP2719-Gal4, Repo-Gal80/LexAop-SytGDP::HA; Crz-LexA/UAS-myr-tdTomato | | |
| b) | DIN>Chr, Crz>Kir | w-; NP2719-Gal4/LexAop-Kir2.1; Crz-LexA/UAS-CsChrimson-tdTomato | | 29 |
| c) | DIN>Chr, Crz>Kir2.1 | w-; NP2719-Gal4/LexAop-Kir2.1; Crz-LexA/UAS-CsChrimson-tdTomato | | 90 |
| | DIN>Chr, Crz>GFP | w-; NP2719-Gal4/LexAop-GFP; Crz-LexA/UAS-CsChrimson-tdTomato | | 91 |
| d) | DIN>Chr | w-; NP2719-Gal4/+; UAS-CsChrimson-tdTomato | | 91 |
| f) | DIN>Chr-tdT, TH>myr-GFP | w-; NP2719-Gal4, Repo-Gal80/LexAop-myr-GFP; TH-LexA/UAS-CsChrimson-tdTomato | | |
| g) | TH>ACR1 | w-;+/+;TH-Gal4/UAS-GtACR1-eYFP | No light | 10 |
| | | | Light | 11 |
| | No retinal | | No light | 10 |
| | | | Light | 9 |
| h) | TH>ACR1 | w-;+/+;TH-Gal4/UAS-GtACR1-eYFP | No light | 26 |
| | | | Light | 31 |
| | No retinal | | No light | 17 |
| | | | Light | 20 |
| | | | No warmth | 71 |
| i) | DIN>Chr, TH>TrpA1 | w-; NP2719-Gal4/LexAop-TrpA1; TH-LexA/UAS-CsChrimson-tdTomato | Warmth | 108 |
| | | | No warmth | 119 |
| l) | DIN>Chr | w-; NP2719-Gal4/+; UAS-CsChrimson-tdTomato | Warmth | 88 |
| | | | No warmth | 71 |
| m) | DIN>Chr, jGCaMP7f | w-; NP2719-Gal4, Repo-Gal80/+; UAS-jGCaMP7f/UAS-CsChrimson-tdTomato | | 7 |
| **Extended Data Figure 10** | | | | |
| a) | SS02547>Chr | w-; R42B02-p65.AD/+;VT059427-Gal4.DBD/UAS- | 41°C heat threat | 32 |

| | | | | |
|---|---|---|---|---|
| | | CsChrimson-tdTomato | | |
| | | | 500 ms pulse, 10 min | 42 |
| | | | Both (500 ms pulse, 10 min) | 31 |
| | | | 10 sec pulse | 31 |
| | | | Both (10 sec pulse, 10 min) | 28 |
| | | | 37°C heat threat | 40 |
| | | | 500 ms pulse, 10 min | 45 |
| | | | Both (500 ms pulse, 10 min) | 39 |
| | | | 10 sec pulse | 39 |
| | | | Both (10 sec pulse, 10 min) | 39 |
| b) | SS01559>Chr | w- ; R29F12-p65.AD/+ ; R88C07-Gal4.DBD/UAS-CsChrimson-tdTomato | 41°C heat threat | 38 |
| | | | 500 ms pulse, 10 min | 30 |
| | | | Both (500 ms pulse, 10 min) | 34 |
| | | | 10 sec pulse | 19 |
| | | | Both (10 sec pulse, 10 min) | 32 |
| c) | $AG_{Desc}$>Chr | w-;R27E07-p65.AD/+;R20F03-Gal4.DBD/UAS-CsChrimson-tdTomato | 41°C heat threat | 38 |
| | | | 500 ms pulse, 10 min | 36 |
| | | | Both (500 ms pulse, 10 min) | 36 |
| | | | 1 sec pulse | 37 |
| | | | Both (1 sec pulse,10 min) | 38 |

| | | | | |
|---|---|---|---|---|
| | | | 37°C heat threat | 35 |
| | | | 500 ms pulse, 10 min | 29 |
| | | | Both (500 ms pulse, 10 min) | 34 |
| | | | 1 sec pulse | 29 |
| | | | Both (1 sec pulse, 10 min) | 16 |
| d) | SS01570>Chr | w-; VT012639-p65.AD/+; VT034795-Gal4.DBD/UAS-CsChrimson-tdTomato | 41°C heat threat | 39 |
| | | | 500 ms pulse, 10 min | 36 |
| | | | Both (500 ms pulse, 10 min) | 38 |
| | | | 10 sec pulse | 45 |
| | | | Both (10 sec pulse, 10 min) | 41 |
| | | | 37°C heat threat | 38 |
| | | | 500 ms pulse, 10 min | 37 |
| | | | Both (500 ms pulse, 10 min) | 37 |
| | | | 10 sec pulse | 41 |
| | | | Both (10 sec pulse, 10 min) | 40 |
| e) | Groom>Chr | w-; R45G01-LexA/+ ; LexAop2-CsChrimson-tdTomato/+ | 41°C heat threat | 66 |
| | | | 500 ms pulse, 10 min | 47 |
| | | | Both (500 ms pulse, 10 min) | 48 |
| | | | 3 sec pulse | 32 |
| | | | Both (3 sec pulse, 10 | 49 |

| | | | | |
|---|---|---|---|---|
| | | | min) | |
| | | | 37°C heat threat | 42 |
| | | | 500 ms pulse, 10 min | 40 |
| | | | Both (500 ms pulse, 10 min) | 33 |
| | | | 3 sec pulse | 38 |
| | | | Both (3 sec pulse, 10 min) | 33 |

**Tabulated p-values and statistical tests performed**

| **Figure + Test used** | **Null hypothesis** | **p-value (significant values in red at alpha = 0.05 using Holm-Bonferroni correction for number of comparisons) Statistical significance indicated in red** |
|---|---|---|
| **Figure 1** | | |
| **b)** Mann-Whitney U-test (groups numbered from left to right: DIN>ACR No light = 1, DIN>ACR Light = 2, DIN>ACR No light no retinal = 3, DIN>ACR Light no retinal = 4, DIN>GFP No light = 5, DIN>GFP Light = 6) | No difference between copulation durations | 1-2: 0.00062, 1-3: 0.271, 1-4: 0.1936, 1-5: 0.96, 1-6: 0.384, 2-3: 0.0018, 2-4: 0.00042, 2-5: 0.00062, 2-6: 0.00042, 3-4: 0.952, 3-5: 0.093, 3-6: 0.059, 4-5: 0.0159, 4-6: 0.034, 5-6: 0.267 |
| **c)** Mann-Whitney U-test (groups numbered from left to right as above) | No difference between copulation durations | 1-2: 0.453, 1-3: 0.711, 1-4: <0.00001, 1-5: 0.0168, 2-3: 0.8181, 2-4: 0.0003, 2-5: 0.05, 3-4: <0.00001, 3-5: 0.05, 4-5: 0.023 |
| **d)** Fisher's exact test, light vs. no light. | No difference between termination probabilities | DIN>ACR1: <0.0001, DIN>GFP: 1.0, +/ACR: 1.0 |
| **e)** Mann-Whitney U test (groups numbered from left to right as above) | No difference between copulation durations | 1-2: 0.00008, 1-3: 0.575, 2-3: 0.00028 |
| **f)** Fisher's exact test. Groups numbered as 1 = DIN>Chr, 2 = No light, 3 = No retinal | No difference in termination probabilities within a timepoint | *Time: 1 minute:* 1-2: <0.0001, 1-3: <0.0001, 2-3: 1.0<br>*Time: 5 minutes:* 1-2: <0.0001, 1- |

| | | |
|---|---|---|
| | | 3: <0.0001, 2-3: 1.0<br><br>*Time: 10 minutes:* 1-2: <0.0001, 1-3: <0.0001, 2-3: 1.0<br><br>*Time: 15 minutes:* 1-2: <0.0001, 1-3: <0.0001, 2-3: 1.0<br><br>*Time: 20 minutes:* 1-2: <0.0001, 1-3: <0.0001, 2-3: 1.0 |
| **F)** Kolmogorov-Smirnov test (data on right) | No difference between distributions of termination times | 0.055 |
| **Figure 2** | | |
| **a)** Kolmogorov-Smirnov test within each pulse width | No difference between distributions of responses based on time into mating | *500 ms pulse:* 0.582<br><br>*1 sec pulse:* 0.882 |
| **d)** Welch's t-test. 10 Minutes, low intensity = 1, medium = 2, high = 3, 15 minutes low intensity = 4, medium = 5, high = 6 | No difference between the $\tau$ parameters across conditions | 1-2: 0.101, 1-3: 0.535, 1-4: 0.0152, 1-5: 0.0059, 1-6: 0.057, 2-3: 0.186, 2-4: 0.019, 2-5: 0.00005, 2-6: 0.0292, 3-4: 0.017, 3-5: <0.000001, 3-6: 0.00619, 4-5: 0.036, 4-6: 0.2219, 5-6: 0.147 |
| **d)** Welch's t-test. 10 Minutes, low intensity = 1, medium = 2, high = 3, 15 minutes low intensity = 4, medium = 5, high = 6 | No difference between the $p_0$ parameters across conditions | 1-2: <0.000001, 1-3: <0.000001, 1-4: 0.4376, 1-5: <0.000001, 1-6: 0.000001, 2-3: <0.000001, 2-4: <0.000001, 2-5: 0.0000158, 2-6: <0.000001, 3-4: <0.000001, 3-5: <0.000001, 3-6: 0.867, 4-5: <0.000001, 4-6: <0.000001, 5-6: <0.000001 |
| **Figure 3** | | |
| **a)** Fisher's exact test, comparisons within a time point (across genotypes) Genotypes numbered as (DIN>Tnt, Light at 30 s = 1, No light = 2, DIN>GFP, Light at 30 s = 3, No light = 4) | No difference between the termination probabilities | *Time: 5 minutes*: 1-2: 1.0, 1-3: 1.0, 1-4: 0.0002, 2-3: 1.0, 2-4: <0.0001, 3-4: 0.0002<br><br>*Time: 10 minutes:* 1-2: 1.0, 1-3: 0.0075, 1-4: 0.0034, 2-3: 0.0034, 2-4: 0.0015, 3-4: 0.669 |
| **b)** Mann-Whitney U test, conditions numbered from left to right | No difference between the distribution of copulation durations | 1-2: 0.617, 1-3: 0.00008, 1-4: 0.00008, 2-3: 0.0002, 2-4: 0.0002, 3-4: 0.829 |
| **c)** Welch's t-test | No difference between | 0.569 |

| | | |
|---|---|---|
| | the $\tau$ parameters across conditions | |
| **c)** Welch's t-test | No difference between the $p_0$ parameters across conditions | <0.000001 |
| **d)** Welch's t-test | No difference between the $\tau$ parameters across conditions | <0.000001 |
| **d)** Welch's t-test | No difference between the $p_0$ parameters across conditions | <0.000001 |
| **f)** Wilcoxon rank-sum test (Pre-DA) | No difference between paired responses pre-DA | 0.0039 |
| **f)** Wilcoxon rank-sum test (Post-DA) | No difference between ratio of residuals post-DA | 0.1289 |
| **Figure 4** | | |
| **a)** Fisher's exact test, groups numbered (Groom>Chr = 1, DIN>GFP = 2, DIN>Tnt = 3) | No difference between termination probability at fixed time point. | *Time*: *1 minute*: 1-2: 1.0, 1-3: 1.0, 2-3: 1.0,<br>*Time: 5 minutes*: 1-2: 1.0, 1-3: 1.0, 2-3: 1.0,<br>*Time: 10 minutes*: 1-2: 0.0792, 1-3: 0.292, 2-3: 0.0252,<br>*Time: 15 minutes:* 1-2: 1.0, 1-3: 0.0002, 2-3: 0.0002 |
| **b)** Posterior probability of a termination probability as large as, or greater than, the observed data given the estimated independent distribution (estimated by the separate light and heat data) | No difference between termination probabilities and that predicted by independent treatments | 0.0091 |
| **c)** Posterior probability of a termination probability as large as, or greater than, the observed data given the estimated independent distribution (estimated by the separate light and heat data) | No difference between termination probabilities and that predicted by independent treatments | *Time: 5 min:* 41°C: 0.4556, 37°C: 0.920<br>*Time: 10 min*: 0.00003<br>*Time: 15 min:* 0.0011 |

| | | |
|---|---|---|
| **f)** Posterior probability of a termination probability as large as, or greater than, the observed data given the estimated independent distribution (estimated by the separate light and heat data) | No difference between termination probabilities and that predicted by independent treatments | 0.0006 |
| **g)** Fisher's exact test, DINs silenced during grooming vs. Heat + Grooming | No difference in termination probabilities between treatments | 0.0009 |
| **Figure 5** | | |
| **e)** Posterior probability of a termination probability as large as or greater than, the observed data given the estimated independent distribution (estimated by the separate light and heat data) | No difference between termination probabilities and that predicted by independent treatments (blue = stabilizing, magenta = switching) | *Time: 10 minutes:* 500 ms pulse: 0.9997, 10 sec pulse: 0.000067<br><br>*Time: 15 minutes:* 500 ms pulse: 0.9982, 10 sec pulse: 0.595 |
| **Extended Data Figure 1** | | |
| **a)** Fisher's exact test (groups numbered DIN>Shi$^{ts}$=1, DIN-Gal4 = 2, +/Shi$^{ts}$=3) | No difference between termination probabilities | 41°C: 1-2: <0.0001, 1-3: <0.0001, 2-3: 0.6833<br><br>Mechanical: 1-2: 1.0, 1-3: 0.475, 2-3: 0.5088 |
| **b)** Mann-Whitney U test, conditions numbered from left to right | No difference between the distribution of copulation durations | 1-2: 0.803, 1-3: 0.00014, 2-3: 0.00036 |
| **Extended Data Figure 2** | | |
| **b)** Mann-Whitney U test, conditions numbered from left to right | No difference between the distribution of copulation durations | 1-2: .258, 1-3: .9601, 1-4: <0.0001, 1-5: <0.0001, 2-3: .4965, 2-4: <0.0001, 1-5: <0.0001, 3-4: <0.0001, 3-5: <0.0001, 4-5: <0.0001 |
| **c)** F test | Likelihood of a slope at least as large as that observed if the true slope were 0. | 0.7609 |
| **Extended Data Figure 4** | | |
| **a)** Posterior probability of a | No difference between | *Time: 10 minutes:* 0.5 sec: |

| | | |
|---|---|---|
| termination probability as large as, or greater than, the observed data given the estimated independent distribution (estimated by the single pulse data) | termination probabilities and that predicted by independent pulses | <0.0001, 1 sec: <0.0001, 5 sec: <0.0001, 10 sec: 0.176, 20 sec: 0.790, 40 sec: 0.652<br><br>*Time: 15 minutes*: 0.5 sec: <0.0001, 1 sec: <0.0001, 5 sec: <0.0001, 10 sec: <0.0001, 20 sec: 0.036, 40 sec: 0.797 |
| **b)** Fisher's exact test, within each genotype, computed across time points (5 min = 1, 10 min = 2, 15 min = 3) | No difference between response to the light pulse across time into mating. | *3 sec pulses:* 1-2: 0.0075, 1-3: 0.0025, 2-3: 0.643<br><br>*6 sec pulses:* 1-2: 0.0023, 1-3: <0.0001, 2-3: 0.0146 |
| **d)** Posterior probability of a termination probability as large as, or greater than, the observed data given the estimated independent distribution (estimated by the single pulse data) | No difference between termination probabilities and that predicted by independent pulses | *Time: 10 minutes:* 5 sec ISI: 0.967, 10 sec ISI: 0.969, 15 sec ISI: 0.9155<br><br>*Time: 15 minutes:* 5 sec ISI: 0.00083, 10 sec ISI: 0.211, 15 sec ISI: 0.828 |
| **e)** Posterior probability of a termination probability as large as, or greater than, the observed data given the estimated independent distribution (estimated by the single pulse data) ($AG_{Desc}$>Chr) | No difference between termination probabilities and that predicted by independent pulses | *Time: 10 minutes*: 1 sec ISI: 0.0051, 5 sec ISI: 0.580, 10 sec ISI: 0.268, 20 sec ISI: 0.329<br><br>*Time: 15 minutes:* 1 sec ISI: 0.0002, 5 sec ISI: <0.0001, 10 sec ISI: 0.011, 20 sec ISI: 0.172 |
| **f)** Posterior probability of a termination probability as large as, or greater than, the observed data, given the estimated independent distribution (estimated either using two red pulses, or one red and one green pulse) | No difference between termination probabilities and that predicted by the pulses acting independently | *DIN>ACR1:* Red only: 0.067, + Green: 0.586<br><br>*DIN>GFP:* Red only: 0.371, +Green: 0.554 |
| **g)** Fisher's exact test comparing the two genotypes | No difference between termination probabilities | 0.4192 |
| **Extended Data Figure 6** | | |
| **g)** Mann-Whitney U test, conditions numbered from left to right | No difference between the distribution of copulation durations | 1-2: 0.357, 1-3:0.9681, 1-4: 0.778, 2-3: 0.289, 2-4: 0.497, 3-4: 0.653 |
| **h)** Fisher's exact test, comparing within columns (Light to no light) | No difference between termination probabilities | With retinal: 0.017<br><br>No retinal: 0.68 |

| | | |
|---|---|---|
| **l)** Welch's t-test | No difference between the $\tau$ parameters across conditions | 0.00133 |
| **l)** Welch's t-test | No difference between the $p_0$ parameters across conditions | <0.000001 |
| **Extended Data Figure 7** | | |
| **b)** Mann-Whitney U test, groups numbered from left to right | No difference between the distributions of the time of switch. | *Left panel:* 1-2: <0.00001, 1-3: <0.00001, 2-3: <0.00001<br>*Right panel:* 1-2: <0.00001, 1-3: <0.00001, 2-3: <0.00001 |
| **Extended Data Figure 8** | | |
| **f)** Mann-Whitney U test, groups numbered from left to right | No difference between the distributions of the time of switch. | 1-2: 0.0044, 1-3: <0.00001, 1-3: <0.00001 |
| **Extended Data Figure 10** | | |
| **a)** Posterior probability of a termination probability as large as or greater than, the observed data given the estimated independent distribution (estimated by the separate light and heat data) | No difference between termination probabilities and that predicted by independent treatments (blue = stabilizing, magenta = switching) | *Time: 10 minutes:* 500 ms pulse: 0.9997, 10 sec pulse: 0.000067<br>*Time: 15 minutes:* 500 ms pulse: 0.9982, 10 sec pulse: 0.595 |
| **b)** Posterior probability of a termination probability as large as or greater than, the observed data given the estimated independent distribution (estimated by the separate light and heat data) | No difference between termination probabilities and that predicted by independent treatments (blue = stabilizing, magenta = switching) | *Time: 10 minutes*: 500 ms pulse: 0.9989, 10 sec pulse: >0.99999 |
| **c)** Posterior probability of a termination probability as large as or greater than, the observed data given the estimated independent distribution (estimated by the separate light and heat | No difference between termination probabilities and that predicted by independent treatments (blue = stabilizing, magenta = switching) | *Time: 10 minutes:* 500 ms pulse: 0.6841, 1 sec pulse: 0.0001<br>*Time: 15 minutes:* 500 ms pulse: 0.0082, 1 sec pulse: <0.000001 |

| | | |
|---|---|---|
| data) | | |
| **d)** Posterior probability of a termination probability as large as or greater than, the observed data given the estimated independent distribution (estimated by the separate light and heat data) | No difference between termination probabilities and that predicted by independent treatments (blue = stabilizing, magenta = switching) | *Time: 10 minutes:* 500 ms pulse: 0.9862, 10 sec pulse: 0.99997<br>*Time: 15 minutes:* 500 ms pulse: 0.9800, 10 sec pulse: 0.152 |
| **e)** Posterior probability of a termination probability as large as or greater than, the observed data given the estimated independent distribution (estimated by the separate light and heat data) | No difference between termination probabilities and that predicted by independent treatments (blue = stabilizing, magenta = switching) | *Time: 10 minutes:* 500 ms pulse: 0.7788, 3 sec pulse: 0.0001<br>*Time: 15 minutes:* 500 ms pulse: 0.693, 3 sec pulse: 0.0012 |

1066
1067
